# A typology of Australian terrestrial bird communities

**DOI:** 10.1101/2025.02.03.635991

**Authors:** Martine Maron, Karlina Indraswari, Jonathan Mills-Anderson, Courtney B. Melton, Helen J. Mayfield, Hugh Possingham, James Radford, April E. Reside, Andrew Bennett, Allan Burbidge, Michael Clarke, Rohan H. Clarke, Robert Davis, Teresa Eyre, Amanda Freeman, Michelle Gibson, Birgita Hansen, Angie Haslem, Jacinta Humphrey, Nigel Jackett, Bryony Palmer, Alex Kutt, Nicholas P. Leseberg, Richard Loyn, Alex Maisey, Golo Maurer, Paul McDonald, Helenna Mihailou, Richard Noske, Fred Rainsford, Julian Reid, Doug Robinson, Katherine Selwood, Jeremy S. Simmonds, Rebecca Spindler, Daniella Teixeira, Ayesha Tulloch, Eric Vanderduys, Simon Verdon, David Watson, James Watson, Hannah Fraser

## Abstract

**Aim:** Increasing interest in holistic measurement of the response of fauna communities to interventions requires suitable community condition metrics. However, the development of such metrics is hindered by the absence of broad-scale typologies at suitable spatial and ecological resolutions. We aimed to derive a preliminary typology of terrestrial bird communities for Australia, based on bird co-occurrence data, and describe and map the likely distribution of each community type across the continent.

**Location:** Mainland Australia, continental islands

**Time period:** 1973-2022

**Major taxa studied:** Aves

**Methods:** We used fine-scale co-occurrence data from standard 2-ha surveys in BirdLife Australia’s citizen-science database. After filtering to reduce bias, we used hierarchical clustering followed by iterative consultation with experts to identify reliably distinct and recognisable terrestrial bird communities across Australia. We used Maxent to model the likely distributions of each community, and developed community descriptions based on each community’s composition and distribution.

**Results:** The resultant typology included 29 reliably distinct and recognisable bird communities with major clusters corresponding with seven broad geographical regions. The distributions of bird communities did not correspond tightly to the boundaries of major vegetation groups, with most communities occurring across multiple vegetation types.

**Main Conclusions:** Our preliminary typology of bird communities provides a standard classification at a continental scale. It newly defines distinct bird communities as entities for which condition benchmarks can be established to allow assessment of their conservation status and monitoring of change over time.

Refinement will enable cryptic communities in areas with sparse data to be identified. The method could be translated to other regions where adequate coverage of data in the form of standardised surveys of fauna are available. Vast biodiversity datasets delivered through citizen science programs provide the opportunity to develop such typologies for fauna communities, as a precursor to developing targeted and informative community condition metrics.

## 1. INTRODUCTION

The new Global Biodiversity Framework has shifted conservation focus from slowing the rate of biodiversity decline towards actively improving the health of biodiversity holistically over time (Xu et al. 2021; GBF 2022). Tracking such ecosystem-level change requires a set of measures that are meaningful for all taxa and the ecological communities of which they are a part (Hughes & Grumbine 2023). We currently monitor only a small fraction of even the most threatened species, and terrestrial ecosystem condition monitoring is limited; where it occurs, it focuses primarily on plant communities and vegetation structure (Lindenmayer et al. 2020) or relies on remotely-sensed indices (e.g. forest integrity index: Grantham et al. 2020). Because vegetation condition may not accurately reflect faunal community condition, methods to measure the overall condition, and monitor trends in condition, of terrestrial fauna communities remain a critical gap, (Redford 1992; Wilkie et al. 2011). A more complete set of indicators of biodiversity health that includes fauna community condition is urgently needed to understand where ecological communities are in good condition and where they are degraded. This in turn informs the need for conservation interventions, to slow, halt and restore faunal community condition, whilst providing a tool to assess the performance of conservation interventions (Nicholson et al. 2021).

Defaunation can be cryptic (Dirzo et al. 2014). For example, the Asian songbird crisis severely depleted bird populations within extensive forest areas due to intensive trapping (Harris et al. 2017; Lees & Yuda 2022), hunting and snaring has depleted mammal communities across vast areas of forest in west Africa (Brashares et al. 2004) and most terrestrial ecosystems within Australia have lost a suite of ‘critical weight range’ mammals (e.g. 35g – 5.5kg) due to novel predation pressure exerted by introduced cats and foxes (McKenzie et al. 2007; Woinarski et al. 2011). Furthermore, factors such as the extent and connectivity of available habitat, at landscape, bioregional or indeed continental scales, can override local habitat condition, such that ostensibly good habitat is no longer occupied by certain species (Radford et al. 2005).

An understanding of fauna community condition is required in many situations, such as impact assessments, fauna community threat assessments, or evaluating effectiveness of conservation management. Sites at which the vegetation is moderately degraded or disturbed can nevertheless support fauna assemblages in very good condition (Ives et al. 2016; Selwood et al. 2018); conversely, structurally and compositionally intact vegetation communities can be fauna depauperate (Redford 1992; Wilkie et al. 2011). In these cases, the standard approach of focussing on vegetation condition serves as a blunt tool. For these reasons, measures more reliably linked to the condition of fauna communities are needed.

A barrier to the development of metrics that directly indicate fauna community condition is the lack of broad-scale typologies of fauna communities at appropriate spatial and ecological resolution. In the terrestrial realm, information used to distinguish ecosystem types tends to focus on soils, terrain, and vegetation (Sattler & Williams 1999; Parkes et al. 2003; Eyre et al. 2015; Keith et al. 2022). At broad scales, vegetation-based ecosystem classifications can be informative proxies for fauna community patterns (Thomson et al. 2009; McAlpine et al. 2016). However, at finer resolutions, the spatial distribution of faunal communities do not necessarily align with those of vegetation communities; in particular, similar faunal communities may occur in multiple vegetation types (Kikkawa 1968) and quite different faunal species compositions can occupy similar vegetation formations either side of biogeographic barriers (Godinho & Da Silva 2018).

While broad biogeographic divisions have long been developed for fauna (Kikkawa & Pearse 1969; Spencer × Horn 1994; Ebach 2012; Hermogenes De Mendonça & Ebach 2020), their resolution is of limited use for understanding expected occurrence of a particular assemblage of species at a given site. Such bioregionalisations provide information on the potential species pool of wide regions, but applications that involve measuring site-level fauna condition require classifications of communities that can be directly translated to the site scale. However, research identifying distinct faunal community types based on site-level co-occurrence of species generally focuses within particular landscapes or regions (Burbidge et al. 2000; Pavey & Nano 2009), with the result that the criteria that distinguish one community from another are not consistent among studies exploring different communities in different regions.

Here, we document the first step required to underpin continental-scale faunal community condition metrics - the development of a data-driven, fauna-based typology of communities. We demonstrate this novel approach for Australia’s terrestrial bird communities. We aim to derive this classification from the site-level co-occurrence patterns of bird species, independently of existing bioregionalisations or other environmental classifications and typologies, to reveal the major communities of birds directly, without relying on proxies. In doing so, we provide a typology that lays the foundation for enabling monitoring and assessment of bird communities as ecological entities.

The ideal foundation for a community typology is a dataset covering the entire area of interest and comprising lists of all species detected at a site at a given point in time. Such data enable the development of a typology independent of vegetation or bioregional classifications, driven solely by co-occurrence patterns of the fauna. Thanks largely to citizen science initiatives, site-level bird survey data are among the most widely available and comprehensive faunal data both in Australia and globally (Troudet et al. 2017; La Sorte × Somveille 2020).

We used fine-scale co-occurrence data from BirdLife Australia’s Birdata database (BirdLife Australia 2022a) to develop an empirically-derived bird community typology for Australia. The database contains more than 410,000 surveys conducted using a standardised method, the 2-ha, 20-min count (Loyn 1986). We applied hierarchical clustering to a large sample of these surveys, followed by iterative consultation with experts, to identify 29 distinct and recognisable terrestrial bird communities across Australia. We used MaxEnt to model the likely distributions of each community, and developed community descriptions based on species that typify the community and summaries of the vegetation groups in which each community is commonly detected. This yielded a preliminary set of distinct, recognisable terrestrial bird communities based on consistent criteria for community classification. Emergence of faunal typologies such as this is a first step in developing metrics to measure and track their condition directly, rather than relying on vegetation as a proxy.

## 2. METHODS

### 2.1 Bird co-occurrence data

In developing our typology, we aimed to define bird communities based on distinct patterns of species co-occurrence, rather than imposing boundaries based on vegetation types or geographic regions. We used site-scale data, collected using a standard 2-ha 20-min survey method, from BirdLife Australia’s Birdata database with surveys from 1973 to 2022, with the vast majority recorded since 1998. Birdata works with the Working List of Australian Birds taxonomy, version 2 (BirdLife Australia 2022b), so that is what is used in this manuscript. It contains hundreds of thousands of bird surveys contributed by thousands of citizen scientists. Surveys are performed and contributed by citizen scientists each year across all parts of Australia.

This typology was developed for terrestrial bird communities only. Bird communities occurring in aquatic ecosystems (e.g. wetlands), shorelines and intertidal zones, the marine environment, and offshore islands were deliberately excluded because birds that typify such environments are much more likely to be surveyed using a range of methods which vary in effort and area surveyed. Terrestrial communities had the advantage (for our analysis) of a commonly-used, standard sampling unit - the 2-ha 20-min survey (Loyn 1986). In this way, we ensured that variation in site-level sampling method or effort was unlikely to be an important driver of the clustering results, which we might have expected had we also included surveys using other methods. Although conceptually, community typology development can be extended to non-terrestrial communities, the approach taken will need to differ.

### 2.2 Data filtering process

We applied a set of filters to the full dataset (Figure 1):

**Figure 1.**
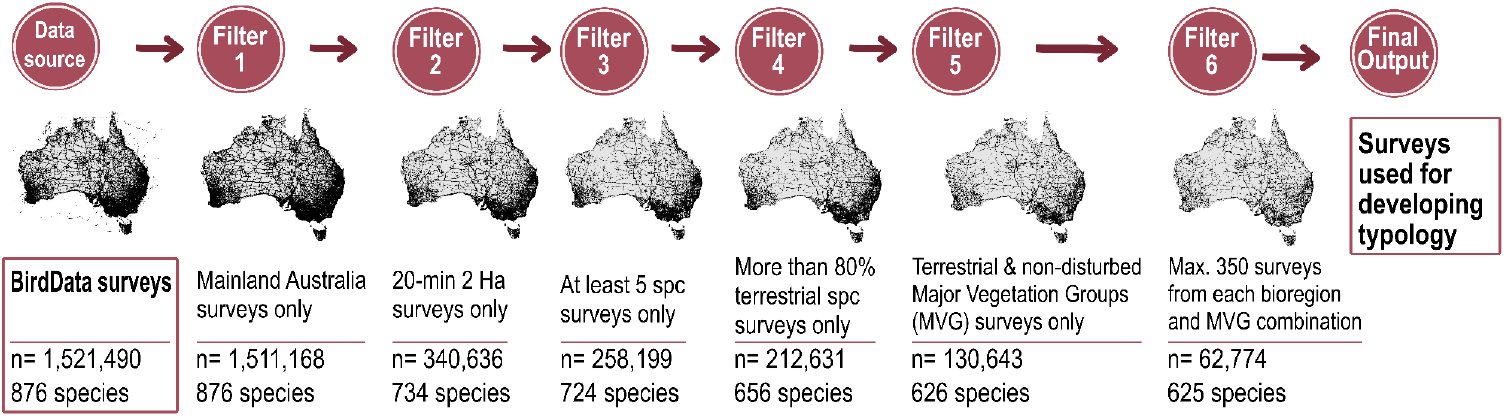
Filtering phases of the Birdata survey data (BirdLife Australia 2022) used to develop bird community typology. Shown are the number of individual bird surveys and number of species included in them after each filtering step, yielding the final sample of 62,774 surveys used for clustering analysis.

1. We excluded surveys done outside of mainland Australia and its continental islands.
2. We excluded surveys that used methods other than the standardised 2-ha 20-min survey method to reduce the chance that the typology reflected different survey methods (Watson 2003) rather than different underlying patterns of co-occurrence of birds. The 2-ha 20-min survey approach is the most commonly used approach for terrestrial bird surveys in Birdata.
3. We excluded surveys with <5 species recorded to improve the chance that surveys we retained contained adequate information about the community from which they were derived.
4. We excluded surveys that included >20% non-terrestrial species (see supplementary material A1 for a classification of terrestrial species).
5. We excluded surveys that were mapped in locations listed as aquatic (Sea and estuaries), highly disturbed (Cleared, non-native vegetation, buildings), or have unclassified or unknown data according to Australia’s National Vegetation Information System (NVIS Technical Working Group 2017).
6. Last, we reduced spatial bias in sample location by stratifying the sites included according to bioregion and vegetation. To do this, we overlaid the Interim Biogeographic Regionalisation for Australia (IBRA) system (DCCEEW 2024) with the Major Vegetation Groups (MVGs, hereafter ‘vegetation groups’), following the approach of Carwardine et al. (2008). This resulted in 1124 distinct combination groups, and we randomly sampled a maximum of 350 surveys for each combination. This resulted in the final sample of 62,774 surveys (Figure 1).

### 2.3 Clustering analysis and expert consultation

We used a non-supervised clustering approach, which allows the data to fall into clusters based on the bird composition of survey lists alone without pre-set rules (Bishop 2006; Miyamoto 2022). We used presence-only data to reduce the influence of very common or abundant species, which tend to contribute little to distinctions among communities (Hirzel et al. 2002; Wilson 2012). We applied hierarchical agglomerative clustering (HAC) on a Euclidean distance matrix using the ward.D2 method in R to group surveys based on similarities in species composition. Euclidean distance was chosen as our aim was to explore potential clusters with no prior assumption of existing patterns (Everitt et al. 2011; Murtagh & Legendre 2014). The ward.D2 algorithm was chosen for its strength in clustering, and that it is less sensitive to outliers (Legendre × Legendre 2012; Murtagh & Legendre 2014). The clustering resulted in a dendrogram depicting major groupings through to minor clusters, with different heights/similarity levels in the dendrogram corresponding to different resolutions of a candidate community typology.

Interpreting unsupervised learning outputs (such as our hierarchical cluster outputs) could be challenging as it required strong qualitative understanding of the data. We iteratively explored and refined our results with ornithological experts. The consultation assisted in identifying the level of similarity at which recognisable and distinct bird communities emerged, and also in identifying clusters that were likely to represent non-naturally occurring bird communities (i.e.: disturbed or degraded communities).

To do this, we consulted with ornithologists and experts (including co-authors on this paper) iteratively. The initial consultation (in May 2022) presented 28 experts with summary data about terminal clusters resulting from truncations of the dendrogram at different heights, ranging from 5 to 50 clusters. The extent to which these ‘candidate communities’ reflected bird communities that were both recognisably distinct to experts, and adequate in capturing expert-judged variation in bird communities across the continent, drove the decision to truncate the dendrogram at a height which yielded 39 clusters as candidate communities. Of these, experts judged that 11 were likely representative of degraded communities, leaving 28 main communities in the candidate typology.

We next sought to refine and corroborate the communities through further consultation with experts, ensuring that the expert group included at least one expert from each of the 28 communities we identified in the previous step. For this consultation, we prepared more detailed information describing each community, excluding the 11 degraded communities which were not considered a best-on-offer site for the typology.

First, we prepared maps of the locations of all surveys classified into each candidate community. Second, we summarised the reporting rate for the 50 most commonly-occurring species in each community (the proportion of surveys classified into that community in which a species occurred). Third, we summarised for each community the proportion of surveys located within each Major Vegetation Group. Finally, we calculated ‘importance value’ indicating the contribution of each species to the distinctiveness of a community within each region by developing a predictive model using random forest, a powerful predictive model algorithm increasingly used for species distribution modelling (Valavi et al. 2021).

We calculated importance scores for species present within each of the seven regions, such that a species’ importance score indicated its contribution to distinctions among communities within the region (not among all communities in the typology). Not all species contribute to distinctions among communities; some are ubiquitous, others equivalently rare among communities. To identify the species that were most important in differentiating communities within each region, we first removed uninformative species using the feature (or variable) selection approach using the Boruta function from the Boruta package in R (Kursa & Rudnicki 2009). Following the removal of these species, we extracted species importance values using the varImp function using the caret and randomForest packages in R (Breiman et al. 2002). Species could contribute to community distinctions through either their presence or absence, and importance values derived from random forest models did not provide this information. Therefore, we additionally used partial dependence plots (using the edarf package (Jones & Linder, 2016) in R) to determine whether a species’ presence or absence was important. Positive values indicated that a species’ presence is important for distinguishing the community from others in the region, negative values indicated that its absence from a community was distinctive, and values closer to zero indicated the species contributed little to differentiating communities. We retained the raw importance values for our analysis and for the community descriptions in the Supplementary Material S1, however we rescaled the importance values (ranging from -1 to 1) when we consulted with experts to aid interpretation.

This set of information about the 28 candidate communities was presented to our larger group of experts at a workshop in December 2022 with 32 participants, as well as in individual follow-up sessions with experts who could not attend. We sought feedback on whether major communities appeared to be missing or merged into single candidate communities, as well as whether the candidate communities were recognisably distinct from one another. Our aim was to identify a ‘Goldilocks’ degree of similarity such that clusters identified based on a lower similarity cutoff were considered by experts as too generic and encompassing multiple distinct community types, but those that emerged based on a higher similarity level (i.e., further splitting the clusters) were too similar for experts to clearly identify as distinct from one another. This round of expert consultation concluded that one of the 28 candidate communities also represented a degraded state, but that three of them should be further split at a marginally lower height on the dendrogram, each into two sub-groups, resulting in a set of 30 tentative communities representing the Australian Bird Community Typology at this step.

### 2.4 Refining and characterising communities

Despite the summary information about each of the 30 tentative communities representing recognisably distinct assemblages, it was evident that many individual surveys comprised primarily widespread species that were common to many communities. Such generic surveys are unhelpful in summarising community types and often resulted in implausible spatial outliers when examining the distribution of a community. We sought to further refine the community descriptions and distributions by excluding these as outliers. We did this by feeding all surveys back into our Random Forest model and extracting information on the probability of each survey being classified into the community it was allocated to. We excluded all surveys that were classified with lower than 60% probability for the main clusters (that were not split further into two), and 50% for the six sub-group communities. This resulted in 17,521 surveys retained across the 30 clusters. This step to retain only surveys that were classified with high confidence reduced considerably the incidence of spatial outliers. Post-outlier removal, we re-calculated species reporting rate, species importance scores, and proportion of surveys classified into different Major Vegetation Groups. An example is shown in Figure 2.

**Figure 2.**
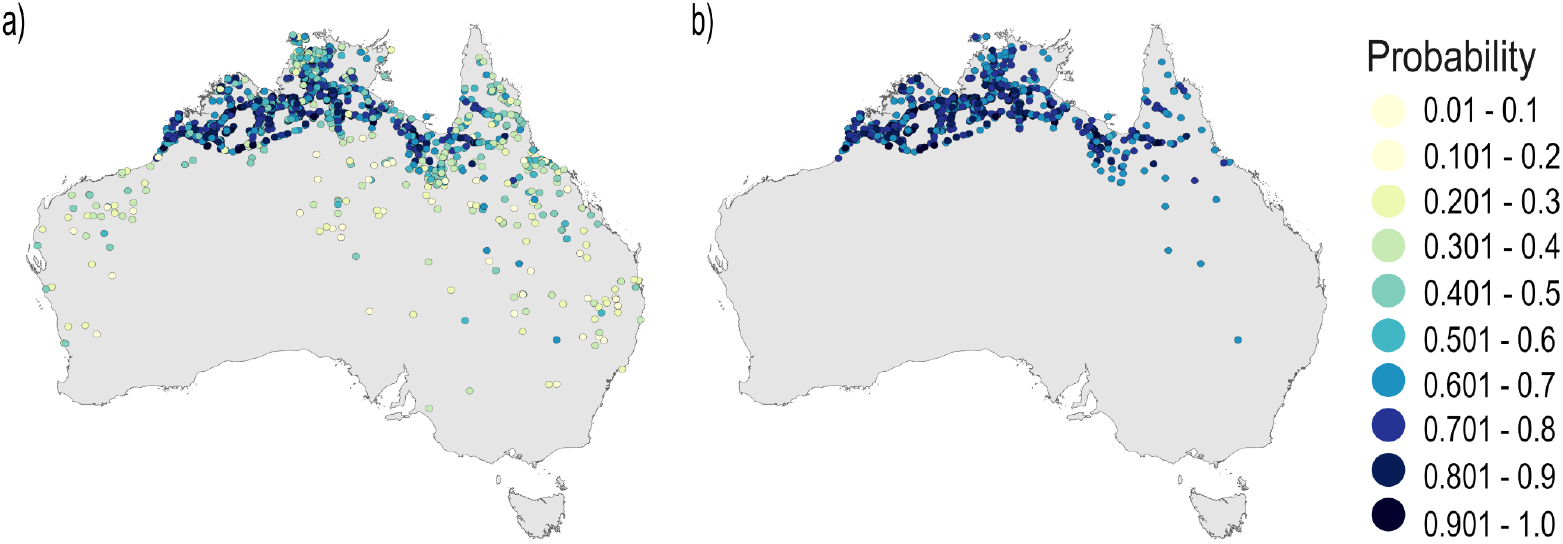
Example of community refinement through outlier exclusion. Figure 2a shows the distribution of all surveys classified into a particular community; b) shows the distribution of only those surveys classified into that community with Pr >0.6.

Following the exclusion of outliers, some of our identified communities retained <100 high-confidence surveys. We retained these sparse communities only where they occurred in areas with sparse survey effort, and experts judged them to be clearly recognisable as distinct communities. For example, Arid Woodland occurred mainly in the Great Victoria Desert, a relatively sparsely-surveyed area, and was retained despite only 68 surveys being classified with high confidence. We removed one cluster at this stage because it only had 19 high-confidence surveys and we considered this insufficient information to define a community. The typology resulting from this stage was again shared with relevant experts, finalising the typology consisting of 29 communities formed from 17,502 surveys. Finally, we ran the random forest model again to analyze the change in our typology’s distinctness. The accuracy of predictive model classification increased from 82% before outlier removal to 98% post-outlier removal.

### 2.5 Identifying community distribution

We used MaxEnt (Phillips et al. 2024) to model the distribution of each of the resultant 29 bird communities. MaxEnt compares the probability densities of a set of spatial environmental variables at georeferenced data locations (in this case, survey locations after outliers were excluded) against a random background sample, following the principle of maximum entropy to make the fewest assumptions, to generate probability distributions of locations suited for the community (Elith et al. 2011). MaxEnt is commonly employed to predict species distributions (e.g., Reside et al. 2010) and, albeit less commonly, to predict community distributions (e.g., Raney & Leopold 2018).

Environmental variables were sourced from WorldClim, TERN, and Geoscience Australia. From WorldClim we used four variables on precipitation (Annual and seasonal precipitation, along with precipitation on the driest and wettest quarter) and four on temperature (minimum temperature, maximum temperature, temperature seasonality, and mean annual temperature). From TERN we used information on total vegetation cover. From Geoscience Australia, we used information on elevation. We also created a distance to water layer by calculating the Euclidean distance from the Digital Earth Australia Waterbodies v2 layer from Geoscience Australia. Further details on the environmental variables used, including selection methods, can be found in supplementary material S3.

## 3. RESULTS

### 3.1 Major clusters

At its coarsest level, the clustering algorithm revealed major splits between terrestrial bird communities of south eastern Australia and those occurring across the rest of the continent; within this second large group, the major split was between communities of the tropical north and those of the arid inland and south-west. These broad splits corresponded broadly to the Bassian, Torresian and Eyrean biogeographic regions (Burbidge 1960; Spencer & Horn 1994), with one group of clusters within the latter comprising mainly degraded communities (Figure 3).

**Figure 3.**
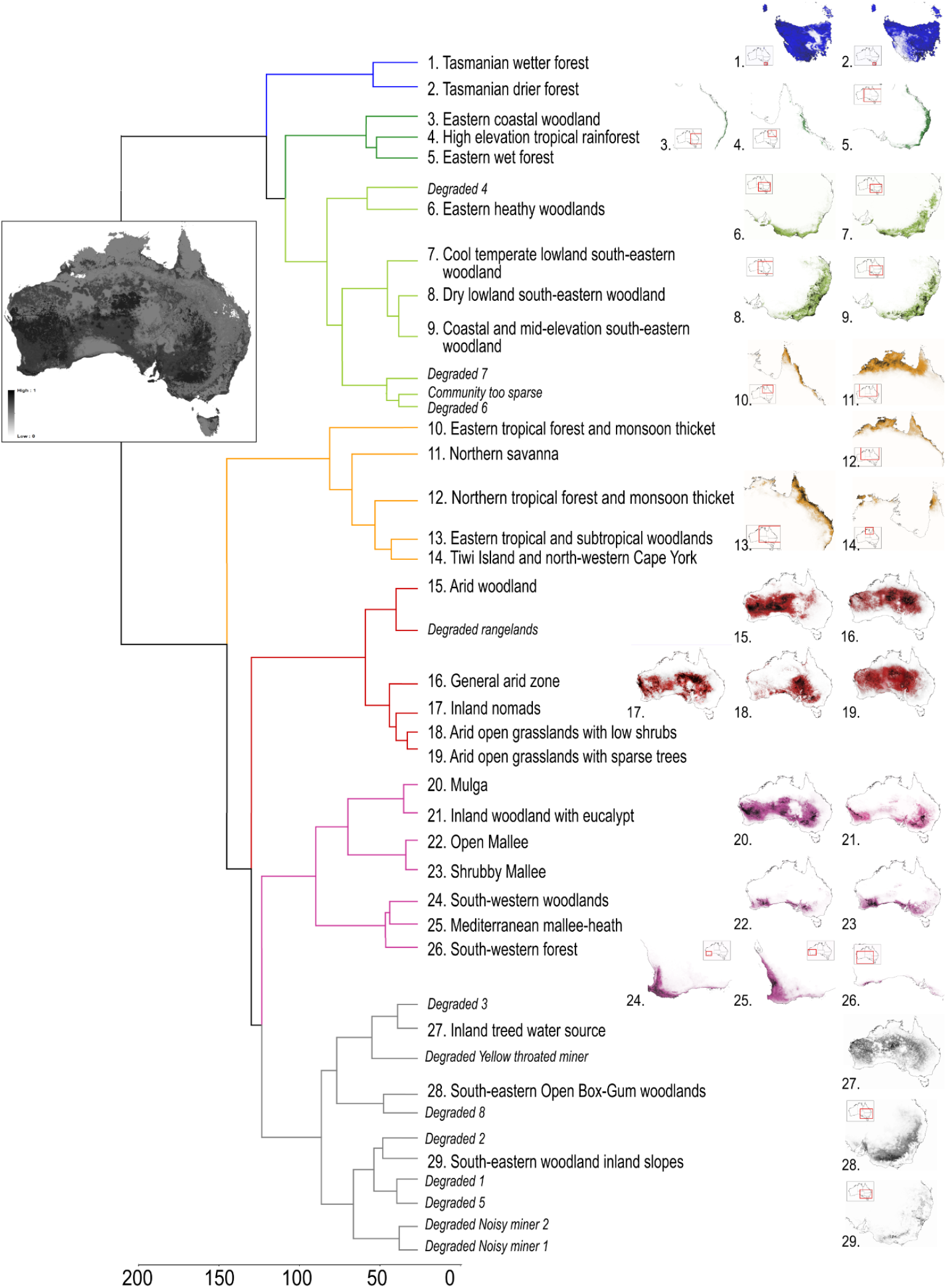
The dendrogram representing the relationships among the community types identified based on clustering. The inset map of Australia shows the stacked distribution of all communities retained in the typology, with darker areas indicating where more communities potentially occur. Colours distinguish higher level groupings representing seven regions. Degraded communities are listed in the dendrogram but are not included in the final typology.

With the input of experts, we identified a level of dissimilarity that split the dataset into 39 clusters, which yielded reliably recognisable bird communities. These 39 clusters fell into seven major groups, which we refer to as ‘regions’ as they reflected broad geographic patterns: Tasmania; eastern wet forests and coastal heath; south-eastern woodland; northern Australia; arid zone; inland eastern woodland; and south-western Australia, mallee and mulga (Figure 3, Table 1, Supplementary material S2).

**Table 1.**
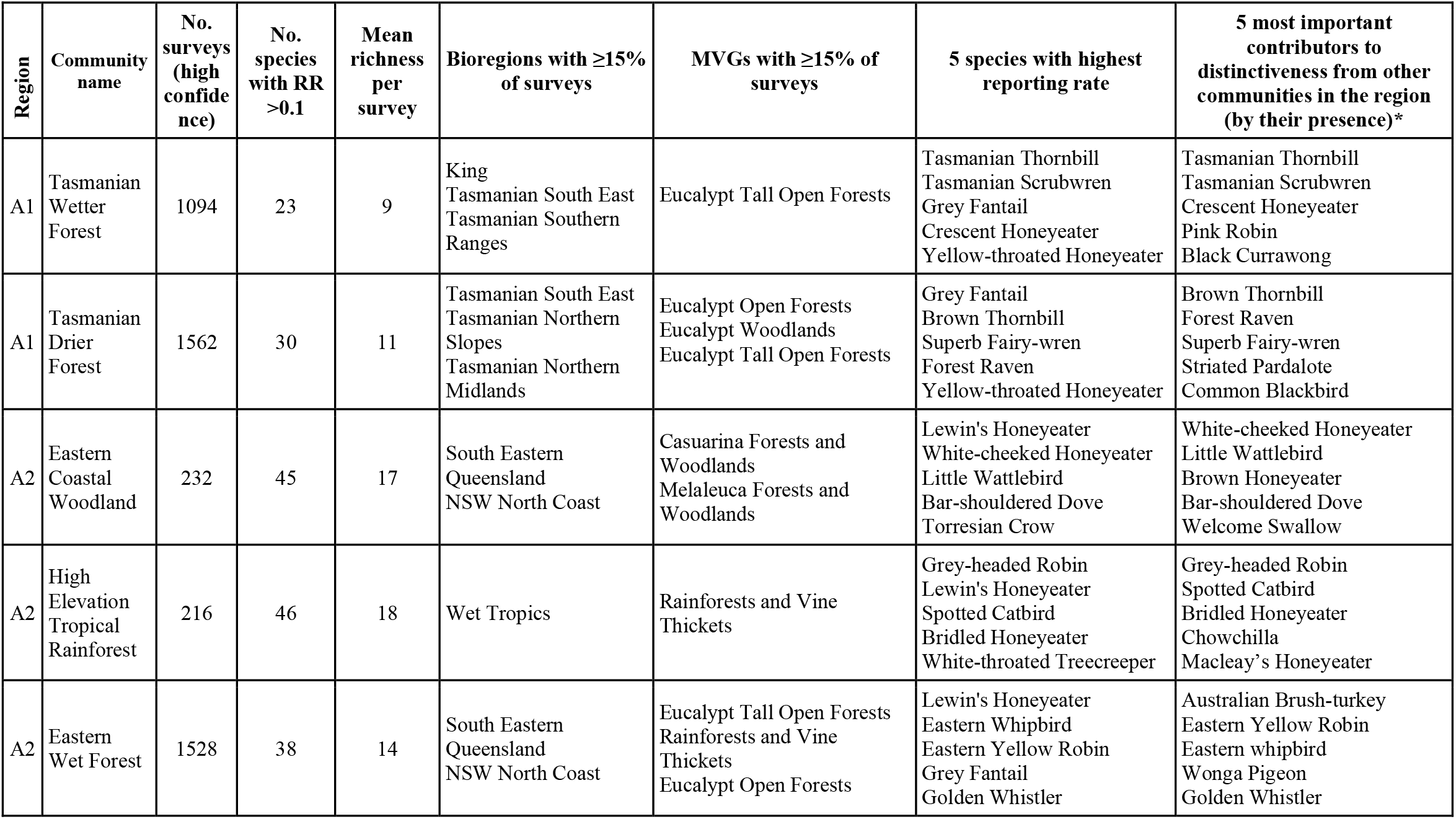
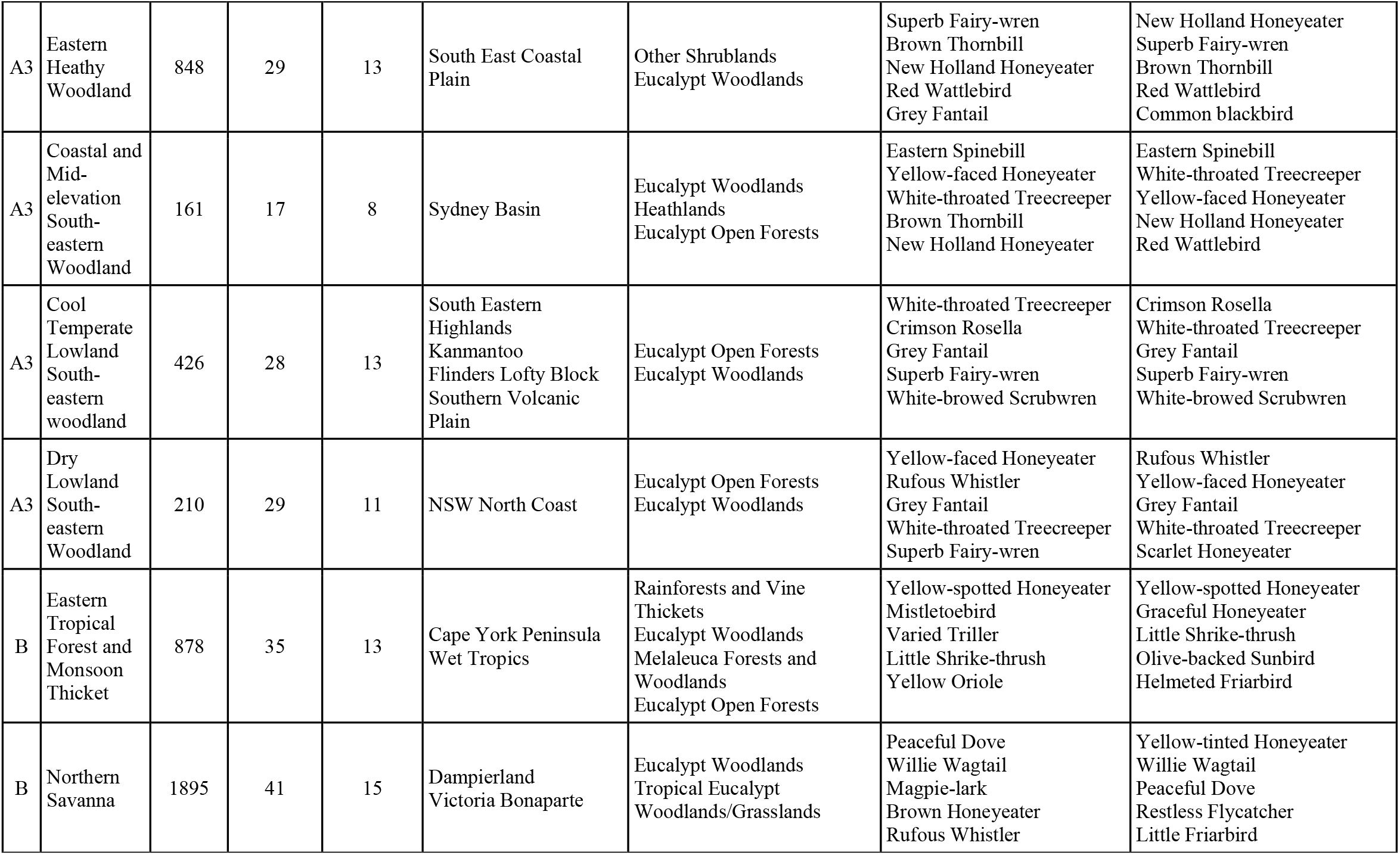
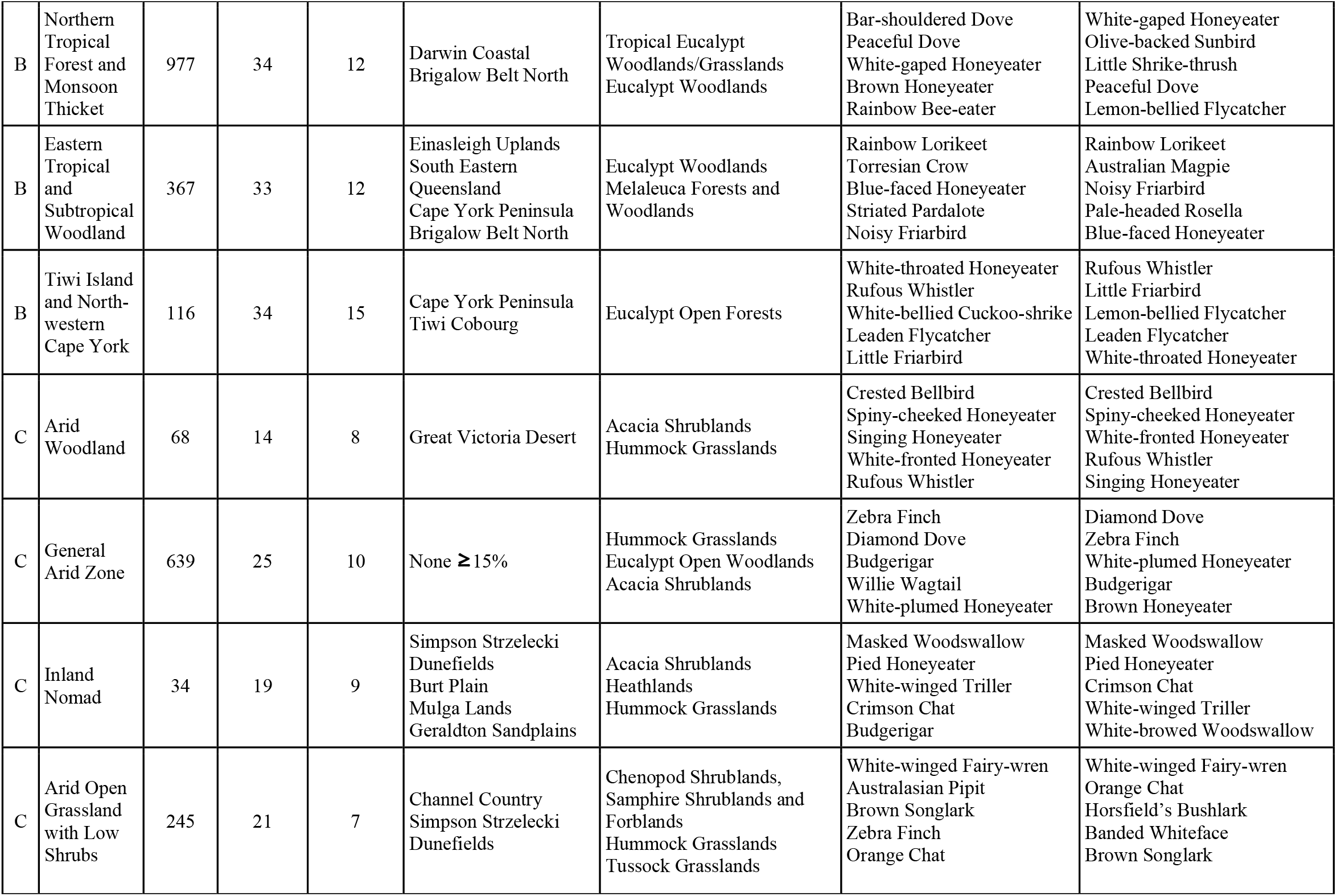
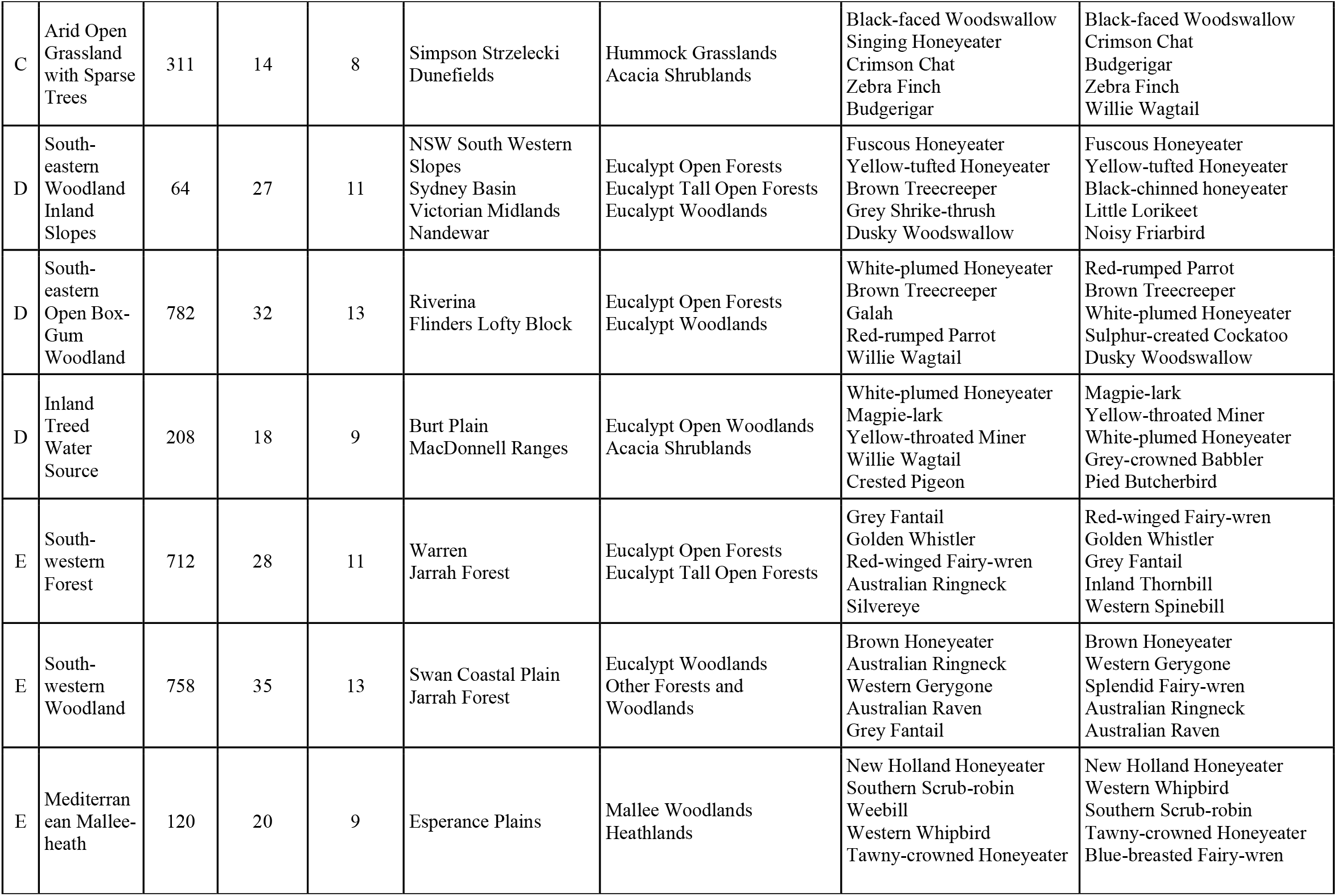
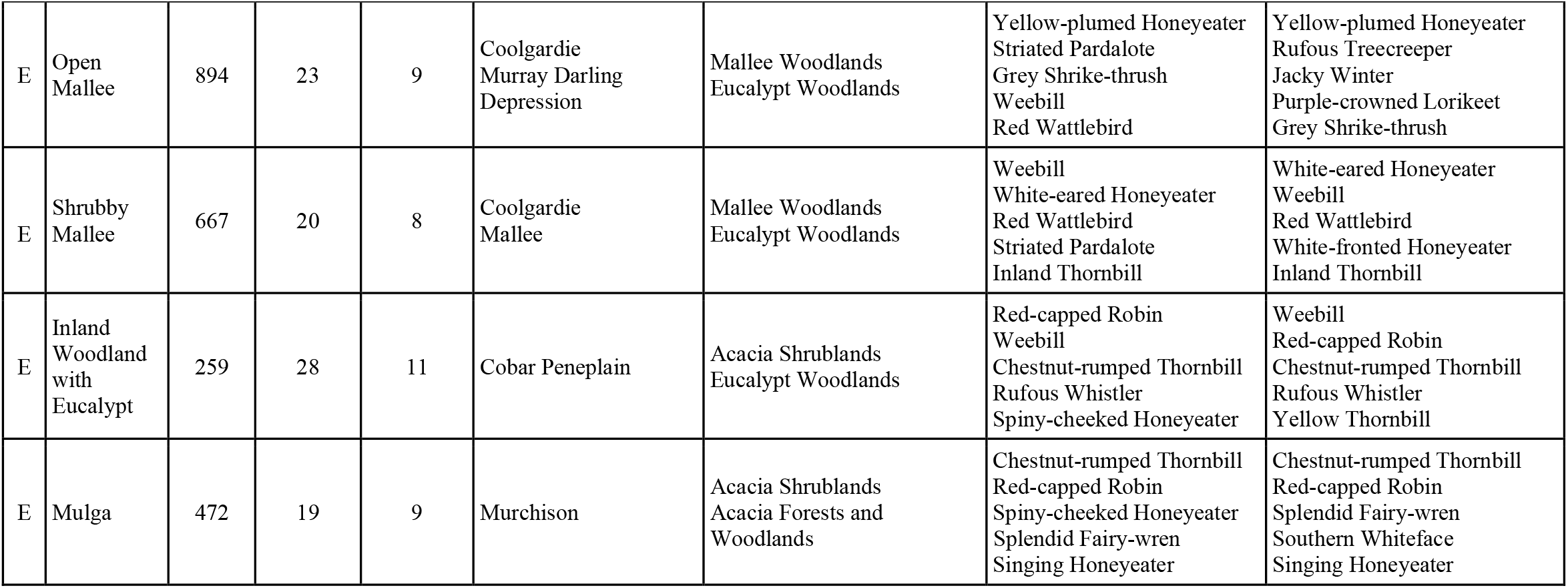
Summary of major terrestrial bird communities and their main characteristics, including the IBRA bioregions and vegetation groups (MVGs) in which each community typically occurred. RR = reporting rate. MVG = Major Vegetation Group. Note that within the Arid Open Grasslands with Low Shrubs community only 4 species were important contributors to distinctiveness. The communities are described in greater detail in Supplementary Material S2.

**Table 1**. SEE END OF MANUSCRIPT

### 3.2 Exclusion of ‘degraded’ and sparse clusters

Of the 39 clusters, twelve were considered by experts to represent groups of very generic survey lists, or strongly anthropogenically altered or degraded examples of bird communities. These clusters were excluded from the typology. Excluded clusters predominantly comprised very widespread and abundant species, or those often associated with highly modified landscapes, such as introduced species or native generalists such as Australian Magpie (RR 0.52), Galah (RR 0.39), and Magpie-lark (RR 0.31), but no or very few distinctive species. Surveys classified into these eleven clusters (n = 12,403) could arise due either to the site supporting a highly simplified or degraded bird community, or simply through the particular timing and/or location of the survey failing to detect species that would, at other times, be present.

The next step of refinement of the typology involved the exclusion of surveys that were not classified into any community with high confidence. This step removed 32,850 surveys, resulting in 17,521 surveys. This smaller and more refined subset of surveys provided a stronger basis for the description of distinct communities, without dilution due to the inclusion of surveys that may not represent that community. At this stage, one cluster was represented by too few high-confidence surveys to retain, resulting in a total of 29 clusters that we judged were distinct, not highly disturbed, and able to be characterised with high confidence. This set of 29 clusters, described using 17,502 surveys, became the basis of the communities in the preliminary typology, and were used for subsequent descriptive analysis and MaxEnt modelling of communities.

Differences among community types in the number of surveys classified with high confidence reflected both the distinctiveness of the communities, as well as variation in the initial number of surveys in each. For example, communities of the south-eastern Australian woodland were well-surveyed, but there were several closely-related communities in this region, and as a result, they only ended up with a modest number of surveys classified with high confidence. Conversely, although *High Elevation Tropical Rainforest* was a small-extent community with relatively few surveys, 78% (216) of the surveys originally classified into that community were classified with high confidence. Greater uncertainty exists about the composition and distinctiveness of communities that are informed by a smaller number of high confidence bird surveys.

### 3.3 Community distribution and naming

The modelled distribution of the 29 communities identified extended across all bioregions of Australia, with most bioregions including more than one community type (Figure 3). Supplementary Material S2 provides a summary of each region’s community types, including their modelled distributions, reporting rates of the most common species, the proportion of surveys that occurred in different vegetation groups, and ordinations showing the multivariate relationships among communities within each region.

The locations of surveys associated with a particular community type corresponded to some extent with particular vegetation groups. In some cases, such as *High Elevation Tropical Rainforest*, this was a very close correspondence, with almost all surveys being located within areas mapped as rainforests and vine thickets (Supplementary Material S2). However, in most cases, a particular community occurred across multiple dominant vegetation groups, and some vegetation groups supported multiple bird communities. For example, surveys classified as belonging to the *Eastern Wet Forest* bird community were located within vegetation groups representing a range of wetter forest types including rainforests and vine thickets, Eucalypt tall open forests, and Eucalypt open forests. Eucalypt woodlands, a widespread vegetation group, included surveys that were classified into at least 11 different bird community types.

Despite the lack of close alignment between vegetation group boundaries and most community types, fine-scale vegetation type and features typically inhabited by each community was generally evident to experts based on distributions and species composition. As such, most of the working names we assigned to communities reflected a combination of vegetation type, specific habitat features, and/or geographic region.

A notable exception was the *Inland Nomad* bird community, a small but very widespread community identified within the arid zone of Australia. This community was dominated by a large number of highly nomadic species, and was considered likely to reflect a temporary influx of species which could occur almost anywhere in the arid zone following suitable seasonal conditions.

## 4. DISCUSSION

Our analysis develops a comprehensive typology of bird communities, representing patterns of local co-occurrence of species, at a continental scale. The 29 terrestrial bird communities we identified describe reliably distinct and recognisable groupings of birds that co-occur, mostly spanning large geographic areas and a range of vegetation groups. Modelled distributions of the communities suggest expected distributions of at least one community type across all parts of mainland Australia. Our typology represents a first step toward developing fauna-focussed metrics of community condition, to enable evaluation and tracking of condition over time at scales from individual properties to regions to entire continents. We propose that this method can be applied to a range of terrestrial faunal groups to generate community typologies, complementary to those that exist for plant communities.

### 4.1 Relationships with vegetation communities

It has long been recognised that the boundaries of plant community types often do not align with those of distinct bird communities (Kikkawa 1968; Recher et al. 1991). Australia’s national vegetation typology (Major Vegetation Groups: MVGs) did not always map neatly to the distributions of bird communities we identified. For example, the *Eastern Heathy Woodland* bird community occurred across 21 different vegetation groups with a maximum of 22% of surveys included in a single vegetation group (other shrublands). In a few cases, a particular community fell largely within a single MVG; for example, 86% of surveys classified as belonging to the *High Elevation Tropical Rainforest* bird community were within the rainforests and vine thickets vegetation group. However, other communities also occurred in this MVG. Many birds are more sensitive to vegetation structure than floristic composition, and sub-strata are particularly important in driving their patterns of local occurrence (Tassicker et al. 2006; Munro et al. 2011). This means that from the perspective of birds, variation within a Major Vegetation Group can be more influential on their local occurrence than variation among them.

At a fine resolution, the boundaries of bird communities may correspond with those of vegetation communities. To identify this alignment, however, we would need to consider the vegetation community types at a much finer scale, and this typology differs among jurisdictions across Australia. For example, the state of Queensland uses a system of Regional Ecosystems, of which there are 1,435 nested within 98 Broad Vegetation Groups (Neldner et al. 2023); however, our analysis identified 21 distinct major bird communities occurring within Queensland. This demonstrates the value of a distinct fauna typology, as assuming a one-to-one relationship between even Queensland’s Broad Vegetation Groups (for example) and bird communities would result in an unnecessarily complex typology. Nevertheless, further work to match finer-resolution vegetation types identified under each of the State vegetation mapping exercises to bird communities (in most cases, a many-to-one relationship) could enable at least an initial indication of likely bird community distributions at a fine scale. Such matching may also enable alignment of bird community types with function-based ecosystem classifications, such as the IUCN Global Ecosystem Typology (Keith et al. 2022)

The classification of ecological communities requires that continuous gradients and temporally dynamic systems reflective of natural systems are reduced to discrete, fixed classes. As such, the resultant typology will necessarily be a highly simplified and subjective representation of nature, imposed by humans to align with human perceptions. This is perhaps most problematic for irruptive bird communities such as those from the arid zone (Pascoe et al. 2021) and northern Australia (Reside et al. 2010). Such typologies also imperfectly reflect ecotones or successional processes through which one community may shift to another over time, following disturbance or lack thereof (Baker et al. 2002; Serong & Lill 2012). Nevertheless, the simplification of complex nature into simplified units is necessary for myriad applications.

### 4.2 Comprehensiveness of the typology

While we are confident that the communities identified represent genuine and distinct community types, more work is required to develop a fully comprehensive typology. Some bird communities are less well surveyed due either to access constraints or smaller extent, and lack the quantity of data that would allow our model to distinguish them. Although the citizen science data we used in our analysis is extensive, the majority of the Australian land mass is sparsely populated and access is challenging, meaning that some areas are poorly represented. So, although some quite localised communities (e.g. *Tiwi Islands and Cape York*) were detected, others will certainly have been missed. Potential examples include small but biogeographically distinct areas such as the Iron and McIlwraith Ranges of far northern Cape York (Johnson × Hooper 1973), the *Banksia* woodlands of coastal south-west Western Australia (Crisp et al. 2001), and the eastern coastal heaths. However, now that we have identified a level of multivariate dissimilarity that consistently yields recognisably distinct bird communities, a targeted approach can be used to evaluate the distinctiveness of additional communities that may have been missed in the original typology due to relevant surveys having been excluded or too sparse. This can be assisted by encouragement of citizen-science efforts to contribute further surveys in areas noted as likely to support such cryptic communities.

We excluded clusters that were deemed not to represent ‘naturally-occurring’ community types. Some of these were excluded as the cluster comprised surveys that were too generic to represent a distinct and recognisable bird community. The 2-ha 20-min surveys we used represent small snapshots of the bird community occupying a particular place at a particular time, and so surveys from one location can vary considerably from hour to hour, let alone from year to year, even in the absence of any local environmental change (Field et al. 2002). Such snapshots can be dominated by a small number of widespread generalist species, even in sites that at other times would yield a richer and more distinct set of species. On the other hand, some sites are persistently dominated by generalists, species commensal with humans, or introduced species. Such assemblages, although they may be accurate representations of the bird community of a site (e.g. in urban areas and artificial habitats), were considered not representative of a naturally-occurring community type, and the 12 clusters apparently representing these were excluded from the typology, given our focus on naturally-occurring bird communities. These judgements are, by necessity, subjective and follow a concept of ‘best on offer’, in that we recognise we do not and can not know the state of communities historically (McNellie et al. 2020).

Our focus on ‘naturally-occurring’ communities was intended to distinguish those representative of what may exist in the absence of intensive pressures recently introduced to the landscape, such as industrial agriculture, urbanisation, and introduced species (Ford et al. 2001). This is quite different to implying that such communities existed in the absence of human intervention, as all of Australia’s ecosystems have been shaped by human practices for millennia. It is therefore important to recognise that what we considered ‘naturally-occurring’ is based on the impressions of contemporary experts familiar with examples of bird communities occurring in what were considered less-disturbed ecosystems. In reality, of course, all of Australia’s ecosystems have been altered substantially in the 200 years since European colonization and associated replacement of Indigenous land management practices (Woinarski & Legge 2013; Fletcher et al. 2021). The extent of these ecosystem-level disruptions is still poorly-recognised and understood, hence our focus on a contemporary ‘best on offer’ conceptualisation of bird community types as a proxy for naturally-occurring communities.

We see our typology as the necessary first step to enable the development of benchmarks of community condition. This, in turn, will enable the assessment of conservation status, tracking of trends, evaluation of management interventions, and reporting of environmental change at multiple scales - from sites and properties, to regions and nationally. The test of a useful typology is therefore not whether it is ‘correct’, but whether it is fit for purpose: yielding categories distinguished on a consistent basis, which users of the typology can recognise reliably. Over the coming years, our proposed typology will doubtless be refined as it is tested through application.

### 4.3 Limitations of the typology

Despite efforts to minimise it, data bias unavoidably influenced our typology. There is a strong spatial bias in the locations most likely to be surveyed across Australia (Barry & Elith 2006), and this will have contributed to variation in sampling adequacy among communities. Indeed, the number of bird surveys that informed each community varied substantially, despite our attempts to improve the spatial evenness of survey effort.

For example, there were very few bird surveys available for the *Inland Nomad* community, relative to the nearby *Northern Savanna* community. As we did not know a priori the spatial distributions of the different communities, we attempted to control for the regional bias in the number of bird surveys by ensuring that no combination of bioregion and vegetation group (MVG) was represented by more than 350 surveys, retaining every survey in combinations with fewer than this number. However, some combinations remained poorly sampled, and so communities local to those areas may have been missed, such as those outlined above.

The dataset we used also contains relevant temporal bias. Northern Australia has fewer surveys during the wet season, as does the arid zone during the hotter summer months. Although Australia’s terrestrial bird fauna is not as dominated by long-distance migrations as northern hemisphere avifaunas (Dingle 2008), such biases are likely to result in reduced detection of summer migrants. While this is unlikely to have led to entire communities being missed, their description and specification are likely to be affected, so future work to examine temporal variation in community composition, particularly in the arid zone and tropical north, will be valuable.

### 4.4 Extension to other taxa and regions

To our knowledge, this is the first continent-wide study exploring fauna-based communities based on co-occurrence, rather than distribution, data. It provides an approach that can be applied in other regions and to other taxa beyond birds. However, our specific method for deriving the typology relied upon extensive citizen science data that used a sampling approach standardised for effort and area, and which is likely to capture a substantial proportion of species within the taxonomic group of interest that are present at the time of the sample. A similar approach could be used in other instances where there is adequate data available for example other structured citizen science programs, potentially including acoustic monitoring of birds, bats or frogs, or collecting eDNA data for aquatic species.

For taxa such as mammals and reptiles, no single survey method is likely to capture a large proportion of species present at a site, and so moderately comprehensive site-level inventories are less likely to be available across the spatial extents required to develop broadscale typologies. For such taxa, approaches will likely rely more heavily on modelled distributions and habitat associations as well as expert elicitation, similar to the approach of Fraser et al. (2018), rather than focussing on co-occurrence. Our method also benefited from the relatively small size of the individual sampling units (2 ha), which meant that it was less likely that the sampled area contained multiple different bird community types, as might be the case if we had used, for example, data from 5-km area searches (another survey method available in Birdata). As such, citizen science programs that use larger units for the aggregation of data (e.g. Southern Africa’s Bird Atlas Program uses 5 × 5 minute ‘pentads’; (Brooks et al. 2022) will require a different approach to the identification of community types.

### 4.5 Applications of a faunal typology

As threats to biodiversity increase in severity and pervasiveness, developing a suite of metrics that allow us to track the condition of broad classes of biodiversity is an increasingly important endeavour. New global commitments by both governments and corporates to achieving ‘nature positive’ outcomes have spurred interest in comparable measurements of biodiversity state and change (Locke et al. 2021; DCCEEW 2022). This is needed to both understand impacts and dependencies on nature, and to measure benefits from positive actions, to inform emerging ‘biodiversity credit’ markets (Yunyue et al. 2024; Swinfield et al. 2024). To answer this call, ecologists need to be able to provide a suite of metrics that are simple but not simplistic, and comparable across locations but sensitive to ecological differences. We argue that such a balance is intermediate between an hypothetical ‘single metric’ of biodiversity condition, and the complexity and expense of tracking every species individually. A small suite of metrics that capture the condition of communities of broad taxa, alongside individual attention to threatened species, would help strike this balance.

The development of this typology was motivated by the need to develop metrics that provide information about the condition of faunal communities, beyond a species-by-species focus and without relying on vegetation condition as a surrogate. The need for a suite of indicators of condition for a given ecosystem is particularly evident when the health of a given fauna community is not well predicted from metrics based on vegetation structure and composition. This occurs especially where the threats that impact fauna community condition are related to direct human disturbance, hunting, and conflict, disease, introduced predators and competitors, or where faunal community responses (e.g., to climate change) substantially lag or lead those of the vegetation community (Redford 1992; Shoo et al. 2005; Wilkie et al. 2011; Woinarski et al. 2015; Scheele et al. 2019).

The typology we have developed enables recognition of fauna community types at the site level, based largely on the combination of species observed. From here, we seek to develop a set of metrics that indicate the condition of each one, in a manner similar to Fraser et al. (2018). This involves identifying the attributes of examples of each community that represent ‘good’ versus ‘poor’ condition examples, with the aim to develop a series of condition metrics able to be calculated from standard samples, in much the same manner as plant community condition metrics. Such metrics are usually standardised relative to their reference condition, representing what is considered “intact” or “best-on-offer” (Thorpe & Stanley 2011; McNellie et al. 2020). These metrics will then enable comparable, standardised tracking and reporting on bird community condition through time, exploring broad correlates of community condition, identifying community types that may satisfy criteria for listing as threatened, and augmenting environmental accounts from the level of individual properties to the national level.

### 4.6 Conclusions

Our empirically-derived typology of the faunal communities of an entire continent demonstrates the value of coordinated, high-quality citizen science efforts. The typology itself is preliminary, but presents an immediate basis for developing metrics of bird community condition for the majority of the terrestrial bird communities of Australia. Ultimately, we hope our approach can initiate similar efforts for other regions and taxa. This work represents a first step to support the development of comprehensive and comparable monitoring of the state and trajectory of faunal communities alongside vegetation, to build a more comprehensive understanding of ecosystem health.

## DATA AVAILABILITY STATEMENT

Summary data for all communities including importance score analysis and reporting rates are provided as Supplementary Material S1. Raw data used for analysis are available under license from BirdLife Australia; a version in which species are anonymised is provided as part of this submission for reference during review, and a final version will be uploaded for public access under a DOI.

## ACKNOWLEDGEMENTS

We thank the Australian Research Council (LP230100179), Bush Heritage Australia, BirdLife Australia, the Queensland Department of Environment, Science and Innovation, and Accounting for Nature for contributing funding and support for this project. Thank you to Stephen Garnett, Monica Awasthy, Andrea Fullagar, Mike Newman, Matthew Taylor, Barry Baker, Richard Major, Chrissy Elmer, Angie Haslem, and Iain Campbell for valuable advice, feedback and discussions.

## SUPPLEMENTARY MATERIAL S1

Summary data comparing reporting rates for all species in each community and importance scores indicating species that distinguish communities within regions accessible here (.xlsx format): https://drive.google.com/drive/folders/1iaYigK0hB_PT9cy1Z98gT9rcNS4BBJX3?usp=sharing

## REFERENCES

Baker J, French K, Whelan RJ. 2002. The edge effect and ecotonal species: bird communities across a natural edge in southeastern Australia. Ecology 83:3048–3059.

Barry S, Elith J. 2006. Error and uncertainty in habitat models. Journal of Applied Ecology 43:413–423.

BirdLife Australia. 2022. Birdata. Occurrence dataset. BirdLife Australia, Melbourne.

Bishop CM. 2006. Pattern recognition and machine learning. Springer, New York.

Brashares JS, Arcese P, Sam MK, Coppolillo PB, Sinclair ARE, Balmford A. 2004. Bushmeat Hunting, Wildlife Declines, and Fish Supply in West Africa. Science 306:1180–1183.

Breiman L, Cutler A, Liaw A, Wiener M. 2002, April 1. randomForest: Breiman and Cutlers Random Forests for Classification and Regression. Available from https://CRAN.R-project.org/package=randomForest x(accessed November 4, 2024).

Brooks M et al. 2022. The African Bird Atlas Project: a description of the project and BirdMap data-collection protocol. Ostrich 93:223–232.

Burbidge N. 1960. The phytogeography of the Australian region. Australian Journal of Botany 8:75–211

Carwardine J, Wilson KA, Watts M, Etter A, Klein CJ, Possingham HP. 2008. Avoiding Costly Conservation Mistakes: The Importance of Defining Actions and Costs in Spatial Priority Setting. PLoS ONE 3:e2586.

Crisp MD, Laffan S, Linder HP, Monro A. 2001. Endemism in the Australian flora. Journal of Biogeography 28:183–198.

DCCEEW. 2022. Nature Positive Plan: better for the environment, better for business. Department of Climate Change, Energy, the Environment and Water, Canberra.

DCCEEW. 2024. Interim Biogeographic Regionalisation for Australia (IBRA). Available from https://www.google.com/url?q=https://datasets.seed.nsw.gov.au/dataset/8e242336-7d10-4630-ae81-e1b6e7464f3c&sa=D&source=docs&ust=1730687306101184&usg=AOvVaw1-jUlS2T_Jph_PytdQk3-6.

Dingle H. 2008. Bird migration in the southern hemisphere: a review comparing continents. Emu - Austral Ornithology 108:341–359.

Dirzo R, Young HS, Galetti M, Ceballos G, Isaac NJB, Collen B. 2014. Defaunation in the Anthropocene. Science 345:401–406.

Ebach MC. 2012. A history of biogeographical regionalisation in Australia. Zootaxa 3392. Available from https://mapress.com/zt/article/view/zootaxa.3392.1.1 x(accessed December 18, 2024).

Elith J, Phillips SJ, Hastie T, Dudík M, Chee YE, Yates CJ. 2011. A statistical explanation of MaxEnt for ecologists: Statistical explanation of MaxEnt. Diversity and Distributions 17:43–57.

Everitt B, Landau S, Leese M, Stahl D. 2011. Cluster analysis. 5th ed. Wiley, Chichester.

Eyre TJ, Kelly AL, Nelder VJ, Wilson BA, Ferguson DJ, Laidlaw MJ, Franks AJ. 2015. BioCondition: A condition assessment framework for terrestrial biodiversity in Queensland. Assessment Manual. Version 2.2. Information Technology, Innovation and Arts, Brisbane.

Field SA, Tyre AJ, Possingham HP. 2002. Estimating bird species richness: How should repeat surveys be organized in time? Austral Ecology 27:624–629.

Fletcher MS, Hall T, Alexandra AN. 2021. The loss of an indigenous constructed landscape following British invasion of Australia: An insight into the deep human imprint on the Australian landscape. Ambio 50.

Ford HA, Barrett GW, Saunders DA, Recher HF. 2001. Why have birds in the woodlands of Southern Australia declined? Biological Conservation 97:71–88.

GBF. 2022. Kunming-Montreal Global biodiversity framework. raft decision submitted by the President, Conference of the parties to the convention on biological diversity, 15th meeting - part 2. Monteal, Canada. Available from https://www.cbd.int/article/cop15-final-text-kunming-montreal-gbf-221222.

Godinho Mbdc, Da Silva FR. 2018. The influence of riverine barriers, climate, and topography on the biogeographic regionalization of Amazonian anurans. Scientific Reports 8:3427.

Grantham HS et al. 2020. Anthropogenic modification of forests means only 40% of remaining forests have high ecosystem integrity. Nature Communications 11:5978.

Harris JBC et al. 2017. Measuring the impact of the pet trade on Indonesian birds. Conservation Biology 31:394–405.

Hermogenes De Mendonça L, Ebach MC. 2020. A review of transition zones in biogeographical classification. Biological Journal of the Linnean Society 131:717–736.

Hirzel AH, Hausser J, Chessel D, Perrin N. 2002. Ecological-niche factor analysis: how to compute habitat-suitability maps without absence data? Ecology 83:2027–2036.

Hughes AC, Grumbine RE. 2023. The Kunming-Montreal Global Biodiversity Framework: what it does and does not do, and how to improve it. Frontiers in Environmental Science 11:1281536.

Ives CD et al. 2016. Cities are hotspots for threatened species. Global Ecology and Biogeography 25:117– 126.

Johnson H, Hooper N. 1973. The Birds of the Iron Range Area of Cape York Peninsula. Australian Bird Watcher 5:80–95.

Jones ZM, Linder FJ. 2016. edarf: Exploratory Data Analysis using Random Forests. The Journal of Open Source Software 1:92.

Keith DA et al. 2022. A function-based typology for Earth’s ecosystems. Nature 610:513–518.

Kikkawa J. 1968. Ecological association of bird species and habitats in eastern Australia; similarity analysis. The Journal of Animal Ecology:143–165.

Kikkawa J, Pearse K. 1969. Geographical distribution of land birds in Australia - A numerical analysis. Australian Journal of Zoology 17:821.

Kursa MB, Rudnicki WR. 2009, December 6. Boruta: Wrapper Algorithm for All Relevant Feature Selection. Available from https://CRAN.R-project.org/package=Boruta x(accessed November 4, 2024).

La Sorte FA, Somveille M. 2020. Survey completeness of a global citizen-science database of bird occurrence. Ecography 43:34–43.

Lees AC, Yuda P. 2022. The Asian songbird crisis. Current Biology 32:R1063–R1064.

Legendre P, Legendre L. 2012. Numerical ecologyThird English edition. Elsevier, Amsterdam.

Lindenmayer D, Woinarski J, Legge S, Southwell D, Lavery T, Robinson N, Scheele B, Wintle B. 2020. A checklist of attributes for effective monitoring of threatened species and threatened ecosystems. Journal of Environmental Management 262:110312.

Locke H et al. 2021. A Nature-Positive World: The Global Goal for Nature. Available from https://library.wcs.org/doi/ctl/view/mid/33065/pubid/DMX3974900000.aspx.

Loyn RH. 1986. The 20-minute search - a simple method for counting forest birds. Corella 10:58–60.

McAlpine C et al. 2016. Integrating plant- and animal-based perspectives for more effective restoration of biodiversity. Frontiers in Ecology and the Environment 14:37–45.

McKenzie NL et al. 2007. Analysis of factors implicated in the recent decline of Australia’s mammal fauna. Journal of Biogeography 34:597–611.

McNellie MJ, Oliver I, Dorrough J, Ferrier S, Newell G, Gibbons P. 2020. Reference state and benchmark concepts for better biodiversity conservation in contemporary ecosystems. Global Change Biology 26:6702–6714.

Miyamoto S. 2022. Theory of Agglomerative Hierarchical Clustering. Springer Singapore, Singapore. Available from https://link.springer.com/10.1007/978-981-19-0420-2 x(accessed November 4, 2024).

Munro NT, Fischer J, Barrett G, Wood J, Leavesley A, Lindenmayer DB. 2011. Bird’s Response to Revegetation of Different Structure and Floristics—Are “Restoration Plantings” Restoring Bird Communities? Restoration Ecology 19:223–235.

Murtagh F, Legendre P. 2014. Ward’s Hierarchical Agglomerative Clustering Method: Which Algorithms Implement Ward’s Criterion? Journal of Classification 31:274–295.

Nicholson E et al. 2021. Scientific foundations for an ecosystem goal, milestones and indicators for the post-2020 global biodiversity framework. Nature Ecology & Evolution 5:1338–1349.

NVIS Technical Working Group. 2017. Australian Vegetation Attribute Manual: National Vegetation Information System, Version 7.0. Department of the Environment and Energy, Canberra. Available from https://www.dcceew.gov.au/sites/default/files/documents/australian-vegetation-attribute-manual-v70.pdf.

Parkes D, Newell G, Cheal D. 2003. Assessing the quality of native vegetation: The ‘habitat hectares’ approach: HABITAT HECTARES. Ecological Management & Restoration 4:S29–S38.

Pascoe BA, Pavey CR, Morton SR, Schlesinger CA. 2021. Dynamics of bird assemblages in response to temporally and spatially variable resources in arid Australia. Ecology and Evolution 11:3977–3990.

Pavey CR, Nano CEM. 2009. Bird assemblages of arid Australia: Vegetation patterns have a greater effect than disturbance and resource pulses. Journal of Arid Environments 73:634–642.

Phillips S J, Dudík M, Schapire RE. 2024. Maxent software for modeling species niches and distributions. Available from http://biodiversityinformatics.amnh.org/open_source/maxent/. x(accessed November 4, 2024).

Radford JQ, Bennett AF, Cheers GJ. 2005. Landscape-level thresholds of habitat cover for woodland-dependent birds. Biological Conservation 124:317–337.

Raney PA, Leopold DJ. 2018. Fantastic Wetlands and Where to Find Them: Modeling Rich Fen Distribution in New York State with Maxent. Wetlands 38:81–93.

Recher HF, Kavanagh RP, Shields JM, Lind P. 1991. Ecological association of habitats and bird species during the breeding season in southeastern New South Wales. Australian Journal of Ecology 16:337– 352.

Redford KH. 1992. The Empty Forest. BioScience 42:412–422.

Reside AE, VanDerWal JJ, Kutt AS, Perkins GC. 2010. Weather, Not Climate, Defines Distributions of Vagile Bird Species. PLoS ONE 5:e13569.

Sattler P, Williams R. 1999. The conservation status of Queensland’s bioregional ecosystems. Environmental Protection Agency, Queensland Government, Brisbane.

Scheele BC et al. 2019. Amphibian fungal panzootic causes catastrophic and ongoing loss of biodiversity. Science 363:1459–1463.

Selwood KE, McGeoch MA, Clarke RH, Mac Nally R. 2018. High-productivity vegetation is important for lessening bird declines during prolonged drought. Journal of Applied Ecology 55:641–650.

Serong M, Lill A. 2012. Changes in bird assemblages during succession following disturbance in secondary wet forests in south-eastern Australia. Emu - Austral Ornithology 112:117–128.

Shoo LP, Williams SE, Hero J-M. 2005. Climate warming and the rainforest birds of the Australian Wet Tropics: Using abundance data as a sensitive predictor of change in total population size. Biological Conservation 125:335–343.

Spencer B, Horn WA. 1994. Report on the work of the Horn Scientific Expedition to Central AustraliaNew facsimile ed. Corkwood Press, Bundaberg, Qld.

Swinfield T, Shrikanth S, Bull JW, Madhavapeddy A, Zu Ermgassen Sose. 2024. Nature-based credit markets at a crossroads. Nature Sustainability 7:1217–1220.

Tassicker AL, Kutt AS, Vanderduys E, Mangru S. 2006. The effects of vegetation structure on the birds in a tropical savanna woodland in north-eastern Australia. The Rangeland Journal 28:139.

Thomson JR, Moilanen AJ, Vesk PA, Bennett AF, Nally RM. 2009. Where and when to revegetate: a quantitative method for scheduling landscape reconstruction. Ecological Applications 19:817–828.

Thorpe AS, Stanley AG. 2011. Determining appropriate goals for restoration of imperilled communities and species. Journal of Applied Ecology 48:275–279.

Troudet J, Grandcolas P, Blin A, Vignes-Lebbe R, Legendre F. 2017. Taxonomic bias in biodiversity data and societal preferences. Scientific Reports 7:9132.

Valavi R, Elith J, Lahoz-Monfort JJ, Guillera-Arroita G. 2021. Modelling species presence-only data with random forests. Ecography 44:1731–1742.

Watson DM. 2003. The ‘standardized search’: An improved way to conduct bird surveys. Austral Ecology 28:515–525.

Wilkie DS, Bennett EL, Peres CA, Cunningham AA. 2011. The empty forest revisited: The empty forest revisited. Annals of the New York Academy of Sciences 1223:120–128.

Wilson JB. 2012. Species presence/absence sometimes represents a plant community as well as species abundances do, or better. Journal of Vegetation Science 23:1013–1023.

Woinarski JCZ et al. 2011. The disappearing mammal fauna of northern Australia: context, cause, and response: Disappearing mammal fauna of north Australia. Conservation Letters 4:192–201.

Woinarski JCZ, Burbidge AA, Harrison PL. 2015. Ongoing unraveling of a continental fauna: Decline and extinction of Australian mammals since European settlement. Proceedings of the National Academy of Sciences 112:4531–4540.

Woinarski JCZ, Legge S. 2013. The impacts of fire on birds in Australia’s tropical savannas. Emu - Austral Ornithology 113:319–352.

Xu H, Cao Y, Yu D, Cao M, He Y, Gill M, Pereira HM. 2021. Ensuring effective implementation of the post-2020 global biodiversity targets. Nature Ecology & Evolution 5:411–418.

Yunyue P, Tong J, Xiaoquan Z, The Nature Conservancy Beijing Representative Office, Beijing, 100600. 2024. Biodiversity credits: Concepts, principles, transactions and challenges. Biodiversity Science 32:23300.

