## Supplementary material for "A typology of Australian terrestrial bird communities": Supplementary Material S2 Community descriptions V3.docx

**Table of contents**

Region A1: Tasmania	2

Tasmanian Drier Forest	4

Tasmanian Wetter Forest	6

Region A2: Eastern wet forests and coastal heath	8

Eastern Coastal Woodland	10

Eastern Wet Forest	12

High Elevation Tropical Rainforest	15

Region A3: South-eastern woodland	17

Eastern Heathy Woodland	19

Coastal and Mid-Elevation South-eastern Woodland	21

Cool Temperate Lowland South-eastern Woodland	23

Dry Lowland, South-eastern Woodland	25

Region B: Northern Australia	27

Eastern Tropical Forest and Monsoon Thicket	29

Eastern Tropical and Subtropical Woodland	31

Northern Tropical Forest and Monsoon Thicket	33

Northern Savanna	35

Tiwi Islands and Cape York	37

Region C: Arid zone	39

Arid Open Grassland with Low Shrubs	40

Arid Open Grassland with Sparse Trees/Tall Shrubs	42

Arid Western Woodland/Tall Shrubland	44

General Arid Zone	46

Inland Nomads	49

Region D: Inland eastern woodland	51

Inland Treed Water Source	53

South-eastern Open Box-Gum Woodland	55

South-eastern Woodland Inland Slopes	58

Region E: South-western Australia, Mallee, Mulga	61

South-western Woodland	66

Mediterranean Mallee-heath	69

Open Mallee	71

Shrubby Mallee	74

Inland Woodland with Eucalypt	77

Mulga	79

### **Region A1: Tasmania**

Two bird communities that occur only on the island of Tasmania were identified: the *Tasmanian Wetter* and *Tasmanian Dr*ier forest bird communities. These communities often occur in close proximity (Figure S1.1), with their distribution influenced by factors such as elevation, aspect and fire history (Wood et al. 2011; Lunn et al. 2018).

Tasmania’s climate is characterized by cool to mild, wet winters and warm, dry summers, resulting in a gradient of vegetation types that support the two identified bird communities. The wetter forest bird community is generally found in the western and southern regions, coinciding with areas of higher annual rainfall, and vegetation is typically dense and closed-canopied (but not exclusively in area of vegetation consistent with the Specht classification of Wet Forest (Specht 1970). These habitats tend to support a subset of bird species that frequent moist, shaded environments including the Tasmanian Scrubwren and Pink Robin. Conversely, the drier forest communities occur more in northern, eastern and central Tasmania, where open forests and woodlands prevail, and species that prefer lower canopy cover and a more open habitat structure are more common (but not exclusively in area of vegetation consistent with the Specht classification of Dry Forest; [[Specht 1970])](https://www.zotero.org/google-docs/?LWpvid).

The Tasmanian region and the two represented communities are unique due to the presence of a high number of endemic species, including the Tasmanian Scrubwren and the Yellow-throated Honeyeater, which contribute to the distinctiveness of its avifauna. These high rates of endemism likely reflect significant recent speciation following the end of the last Ice Age when sea levels rose isolating Tasmania from the Australian mainland around 8-12,000 years ago (Rldpath & Moreau 1966). Common and widespread species such as the Grey Shrike-thrush, Superb Fairy-wren, and Grey Fantail can be found in both communities, highlighting their adaptability to a range of habitat conditions across the island.

An additional feature of Tasmanian bird communities is the prevalence of regional migrants, and Bass Strait migrants. Regional migrants, particularly altitudinal migrants, move away from higher elevation areas (e.g. >800 m) to lower altitudes including near-coastal settings in the austral winter (Rldpath & Moreau 1966; Abbott 1972; Brereton & Taylor 2000). For Bass Strait migrants, some or all of the Tasmanian populations of these species migrate to the Australian mainland for variable periods over winter (Abbott 1972). Examples of obligate Bass Strait migrants include Satin Flycatcher, Dusky Woodswallow and Pallid Cuckoo, whilst examples of partial Bass Strait migrants (only some of the population migrates) include Flame Robin, Silvereye and Grey Fantail. Bass Strait migration, particularly the significant over-water crossing of several hundred kilometers, is also a relatively recent phenomenon that will have been influenced by sea-level rise 8-12,000 years ago (Rldpath & Moreau 1966).

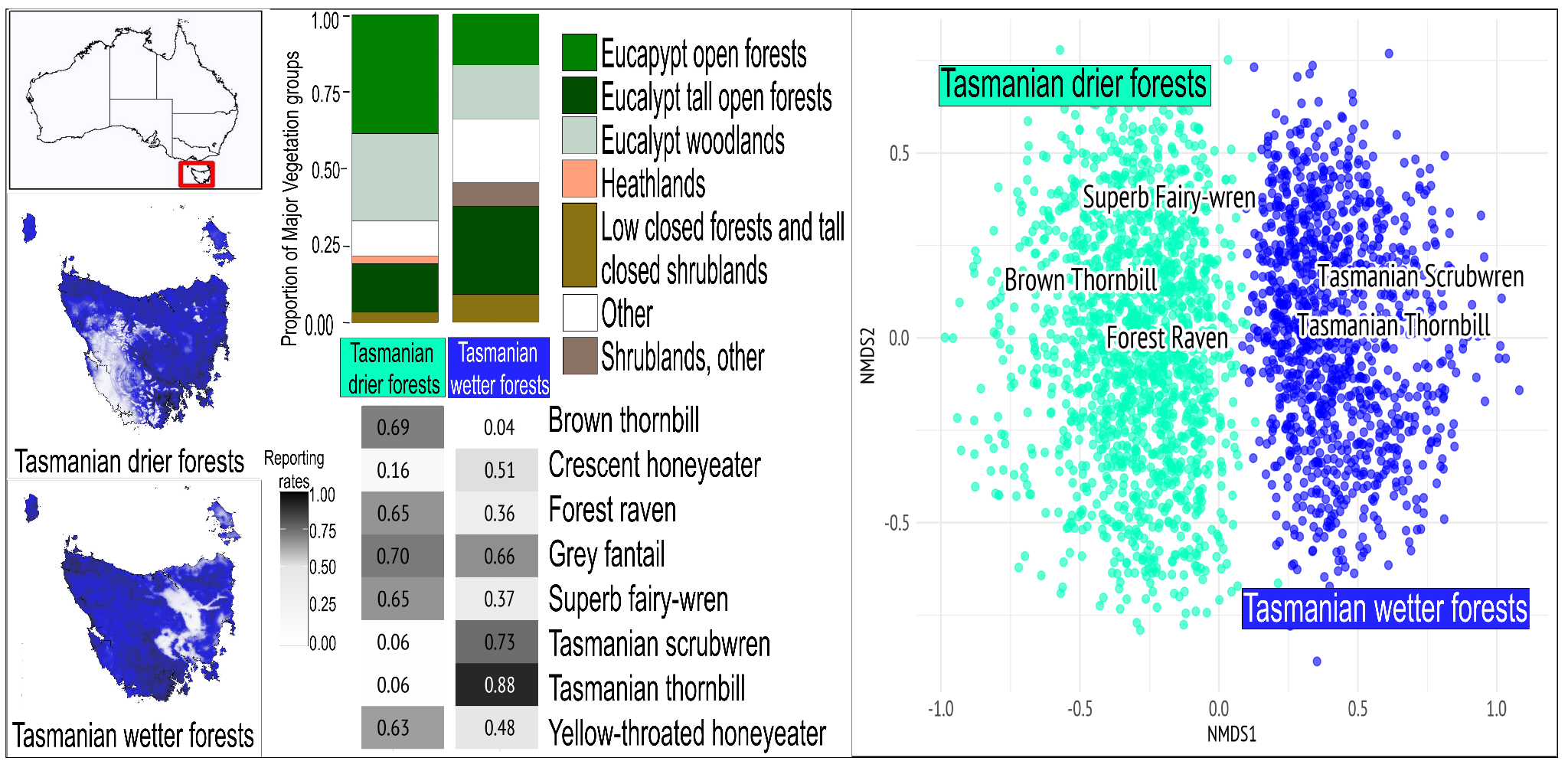

**Figure S1.1** From left to right: far left) the top map indicates location of region, the bottom maps show distribution of communities, darker shading represents higher likelihood of occurrence. Middle, top is the comparison of proportion of Major Vegetation Groups (MVG) and bottom is the comparison of Reporting Rates (RR) of the top 5 most commonly occurring species from each community. Far right, a non-metric multidimensional scaling (NMDS) plot depicting the different important species distinguishing communities.

#### **Tasmanian Drier Forest**

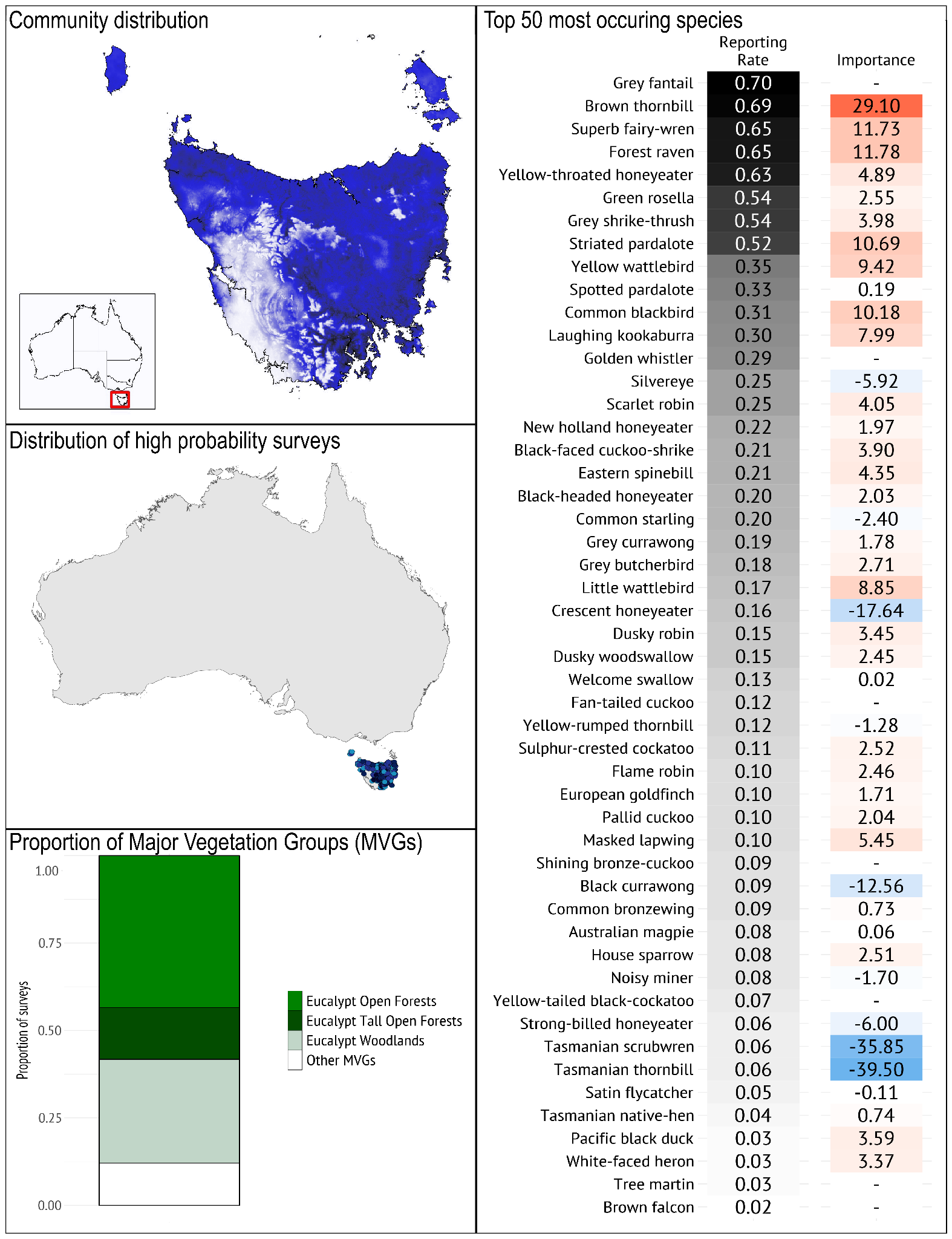

***Geographical occurrence***

The *Tasmanian Drier Forest* community occurs in eucalypt forest and woodland across Tasmania and the larger Bass Strait islands, except for the south-west of Tasmania, where the Tasmanian Wetter Forest bird community dominates.

***Species composition***

*Typical species*: The *Tasmanian Drier Forest* bird community contains many species typical of woodlands and open forests across eastern Australia, including brown thornbill, grey fantail, superb fairy-wren and grey shrike-thrush, but also has high prevalence of Yellow-throated Honeyeater, a Tasmanian endemic, and the Forest Raven. The community includes 8 Tasmanian endemic species. It is much less likely than the *Tasmanian Wetter Forest* bird community to support Tasmanian Thornbill, Tasmanian Scrubwren, and Pink Robin.

*Guild structure*: This community is dominated by small- to medium-sized insectivores and nectarivores, most of which forage in the canopy and subcanopy, and fewer ground-foragers than the Tasmanian Wetter Forest community.

*Temporal dynamics*: There is a large group of summer migrants, arriving in spring leaving Tasmania in Autumn to overwinter on the mainland. For example, Grey Fantail, cuckoos, Flame Robin, Satin Flycatcher, Black-faced Cuckoo-shrike, Dusky Woodswallow, Tree Martin, Striated Pardalote, Eastern Spinebill (Abbott 1972; Brereton & Taylor 2000).

*Geographical variation*: As this community is restricted to Tasmania and Bass Strait, there is limited compositional variation across its range; all typical species are distributed throughout its extent.

***Vegetation/habitat associations***

The *Tasmanian Drier Forest* bird community occurs mainly in Eucalypt open forest and Eucalypt woodlands, and although it is recorded in Eucalypt Tall Open Forests, it is less likely to occur there than is the *Tasmanian Wet Forest* bird community.

#### **Tasmanian Wetter Forest**

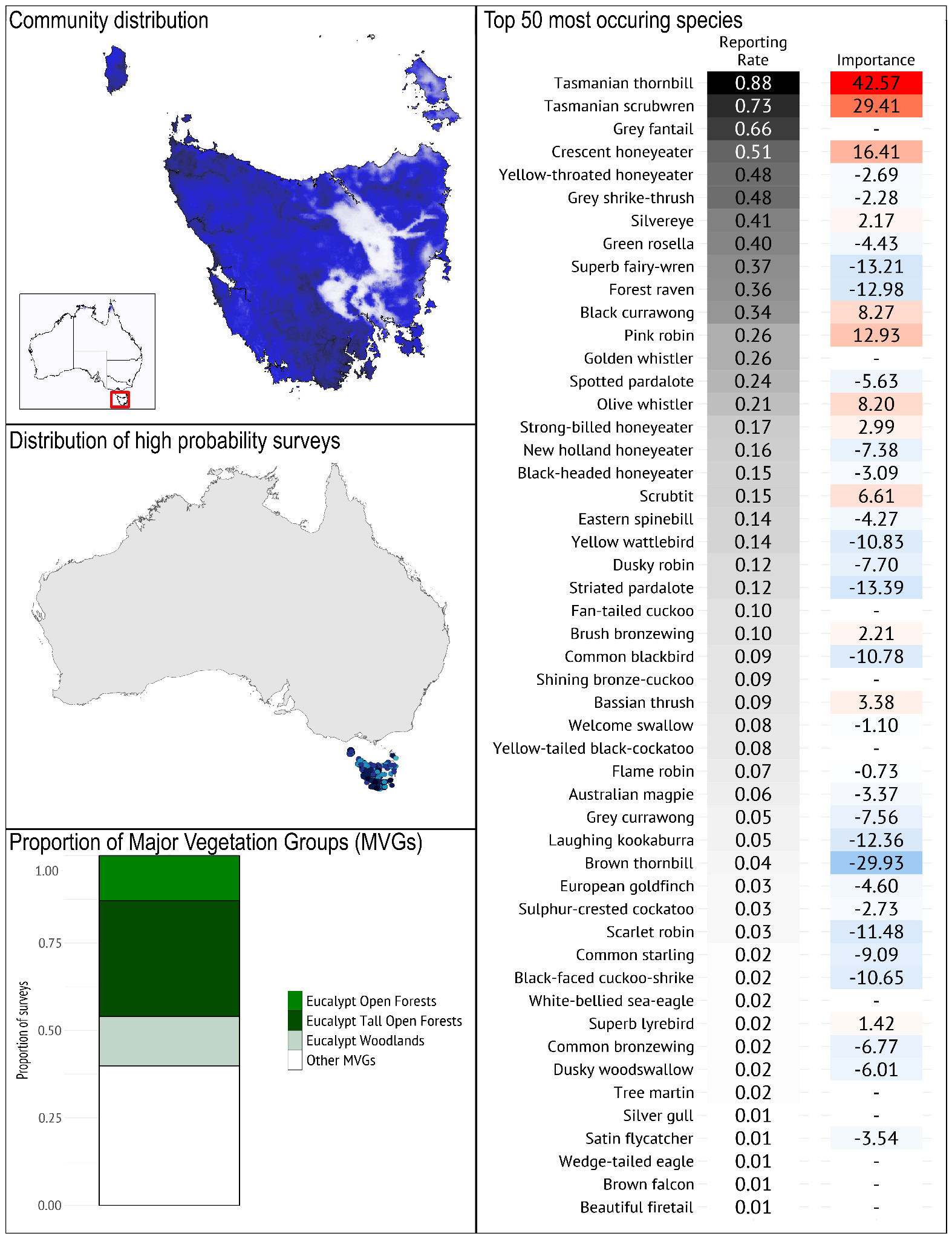

***Geographical occurrence***

This community occurs in more mesic forests across Tasmania with the exception of some parts of the central inland.

***Species composition***

*Typical species*: It is typified by the prevalence of Tasmanian Thornbill, Tasmanian Scrubwren, Grey Fantail, Crescent Honeyeater, and Grey Shrike-thrush. The presence of Tasmanian Thornbill and Tasmanian Scrubwren is a strong indicator of the bird community belonging to the *Tasmanian Wetter Forest* community rather than to the *Tasmanian Drier Forest* community.

*Guild structure*: The community is dominated by small to medium sized insectivores, as well as species that forage on the ground or in dense understory, especially gullies (Tasmanian Scrubwren, Pink Robin, Scrubtit).

*Temporal dynamics*: Some species in this community show local altitudinal movements presumably to avoid the harshest weather in winter (Ridpath & Moreau 1966). As with drier forests, there is also a large group of summer migrants such as Crescent Honeyeater, Slivereye, Grey Fantail, cuckoos, and Tree Martin (Abbott 1972; Brereton & Taylor 2000).

*Geographical variation*: Some species, such as Black Currawong, Tasmanian Thornbill, Pink Robin, Olive Whistler and Crescent Honeyeater, are more likely to occur in the cooler higher altitude tall eucalypt and cool temperate rainforests (Thomas 1980).

***Vegetation/habitat associations***

The *Tasmanian Wetter Forest* bird community is more likely to be found in areas with the Eucalypt tall open forest Major Vegetation Group (MVG) than the *Tasmanian Drier Forest* bird community.

### **Region A2: Eastern wet forests and coastal heath**

This region consists of three communities, *Eastern Coastal Woodland*, *Eastern Wet Forest*, and *High Elevation Tropical Rainforest*. The NVIS Major Vegetation Groups (MVGs) that these bird communities occur in are much more distinctive than other regions with few overlapping MVGs between communities. Important aspects of the vegetation that are likely to be driving differences in the bird assemblages between communities include the relative proportion of Eucalypts versus non-Eucalypts in the overstorey and the relative dominance and density of the understorey vegetation.

This community comprises a mixture of resident and non-resident nectarivorous and insectivorous birds that show large movement patterns depending on the availability of nectar and insects, that have diverse and complicated patterns of seasonal movements driven by a combination of seasonal flowering patterns of plant species and rainfall events. The plant communities are strongly influenced by fire regimes, and in particular wildfires, which can influence the composition and abundance of the bird community (and movements of birds up and down the coast). The majority of the bird species in these communities are not hollow-dependent, so their distribution is likely driven more by the availability of food resources and environmental gradients.

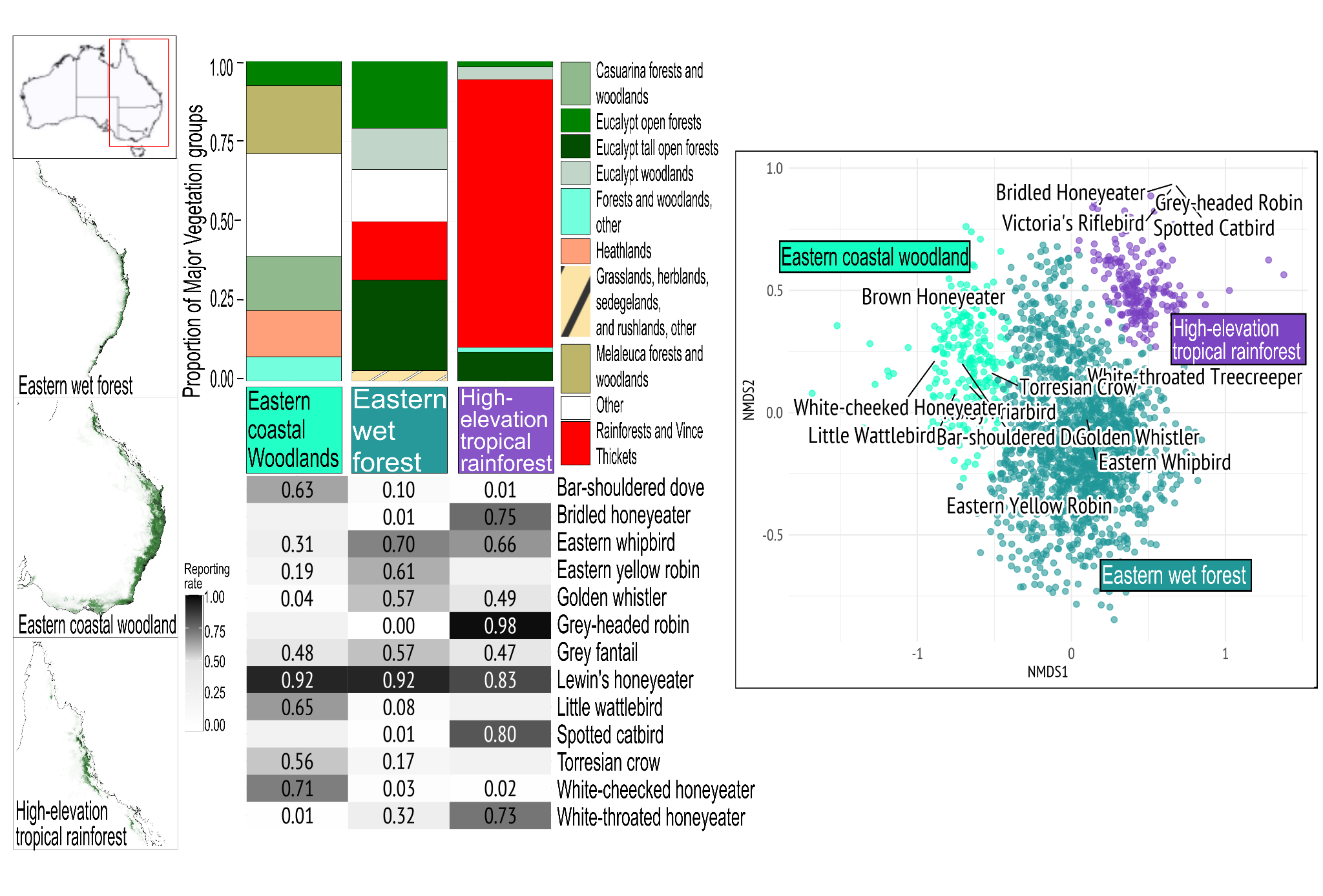

**Figure S1.2** From left to right: far left) the top map indicates location of region, the bottom maps show distribution of communities, darker shading represents higher likelihood of occurrence. Middle, top is the comparison of proportion of Major Vegetation Groups (MVG) and bottom is the comparison of Reporting Rates (RR) of the top 5 most commonly occurring species from each community. Far right, a non-metric multidimensional scaling (NMDS) plot depicting the different important species distinguishing communities.

#### **Eastern Coastal Woodland**

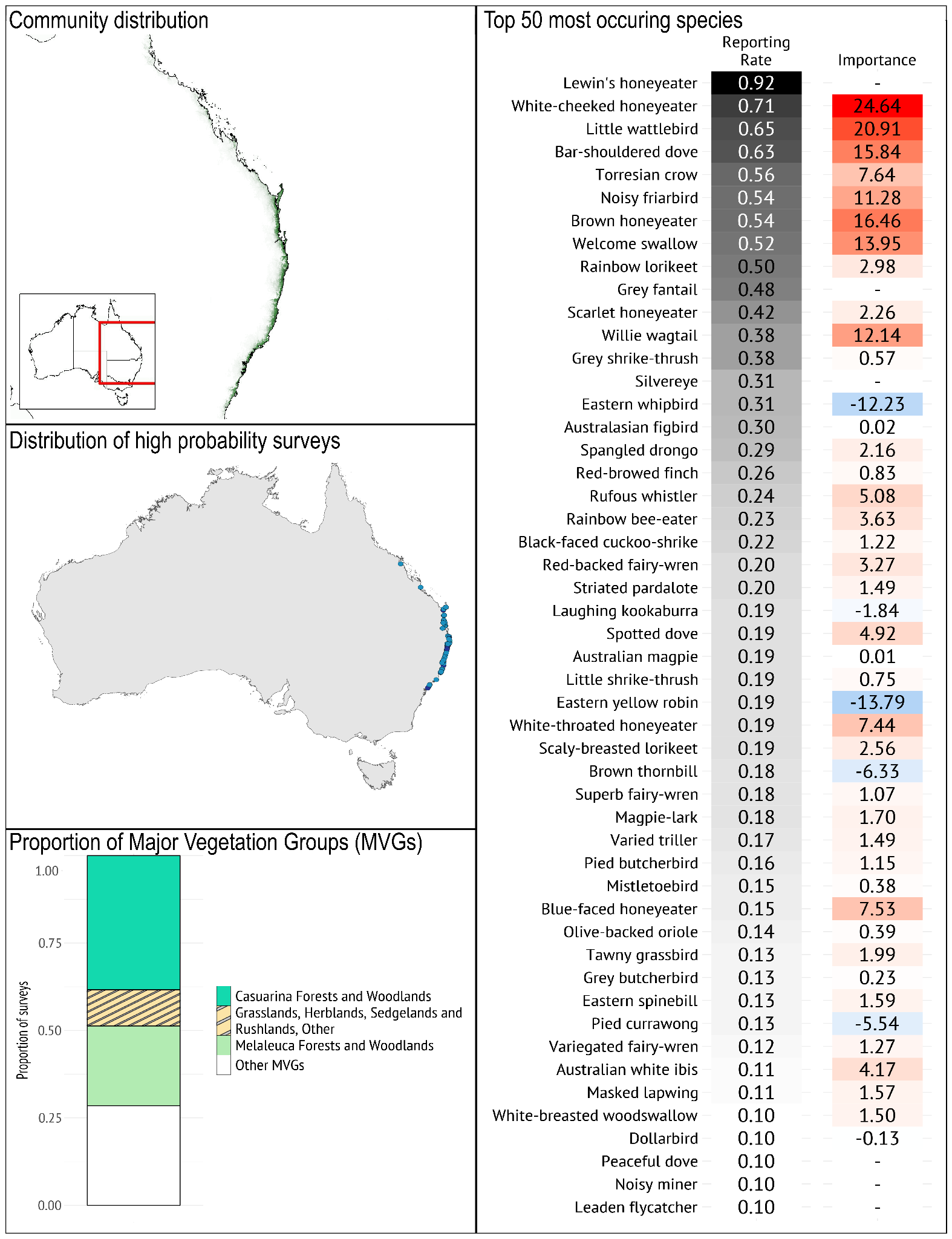

***Geographical occurrence***

The *Eastern Coastal Woodland* bird community occurs in a thin strip down Australia’s east coast, mainly in fine-scale mosaics of non-eucalypt dominated coastal woodlands, heath and heathy woodlands, most commonly between southern Queensland to the NSW-Vic border. It shows a species’ richness gradient from north to south, with some species absent in the southern part of the community - for example, Bar-shouldered Dove and Torresian Crow. The distribution of this community coincides strongly with the distribution of *Banksia* spp along the eastern coast of Australia (especially Banksia integrifolia [[Atlas Of Living Australia 2024])](https://www.zotero.org/google-docs/?YhPJs3).

***Species composition***

*Typical species*: The *Eastern Coastal Woodland* bird community is typified by the prevalence of Lewin’s Honeyeater, White-cheeked Honeyeater, Little Wattlebird, Bar-shouldered Dove and Torresian Crow. The prevalence of the White-cheeked Honeyeater, Little Wattlebird, Brown Honeyeater and Bar-shouldered Dove, and the lower occurrence of the Golden Whistler and White-throated Treecreeper, distinguish it from other communities in this group.

*Guild structure*: A diversity of honeyeaters, including several medium-sized to large species, dominates this group.

*Temporal dynamics*: This community is only moderately temporally variable with seasonal and locally nomadic movements by some honeyeaters e.g. Scarlet Honeyeater and relatively low prevalence of summer migrants e.g. Dollarbird.

*Geographical variation*: Although narrowly distributed close to the coast, this community has a broad north-south distribution, with generally higher species diversity in the north, and a handful of species only occur, or are more common, in northern examples of the community; e.g., Dusky Honeyeater, Bar-shouldered Dove and Torresian Crow.

***Vegetation/habitat associations***

This community is most prevalent in Casuarina forests and woodlands, and Melaleuca forests and woodlands Major Vegetation Groups (MVGs) with substantial occurrence in Heathlands and Other Forest and Woodlands MVGs. It is much less likely to occur in Eucalypt tall open forests, Eucalypt woodlands, or rainforest and vine thickets which support the other communities in this region. The community occurs in a fine-scale mosaic of different vegetation types, including heathlands and Banksia woodlands. These plant communities typically comprise a diversity of proteaceous plants that provide year-round availability of floral resources (nectar and pollen), including largely pollinator-dependent plant taxa such as Banksia, Persoonia, Hakea, Lambertia, and Grevillea (Armstrong 1979; Cardillo & Pratt 2013; Davis et al. 2014, 2022).

#### **Eastern Wet Forest**

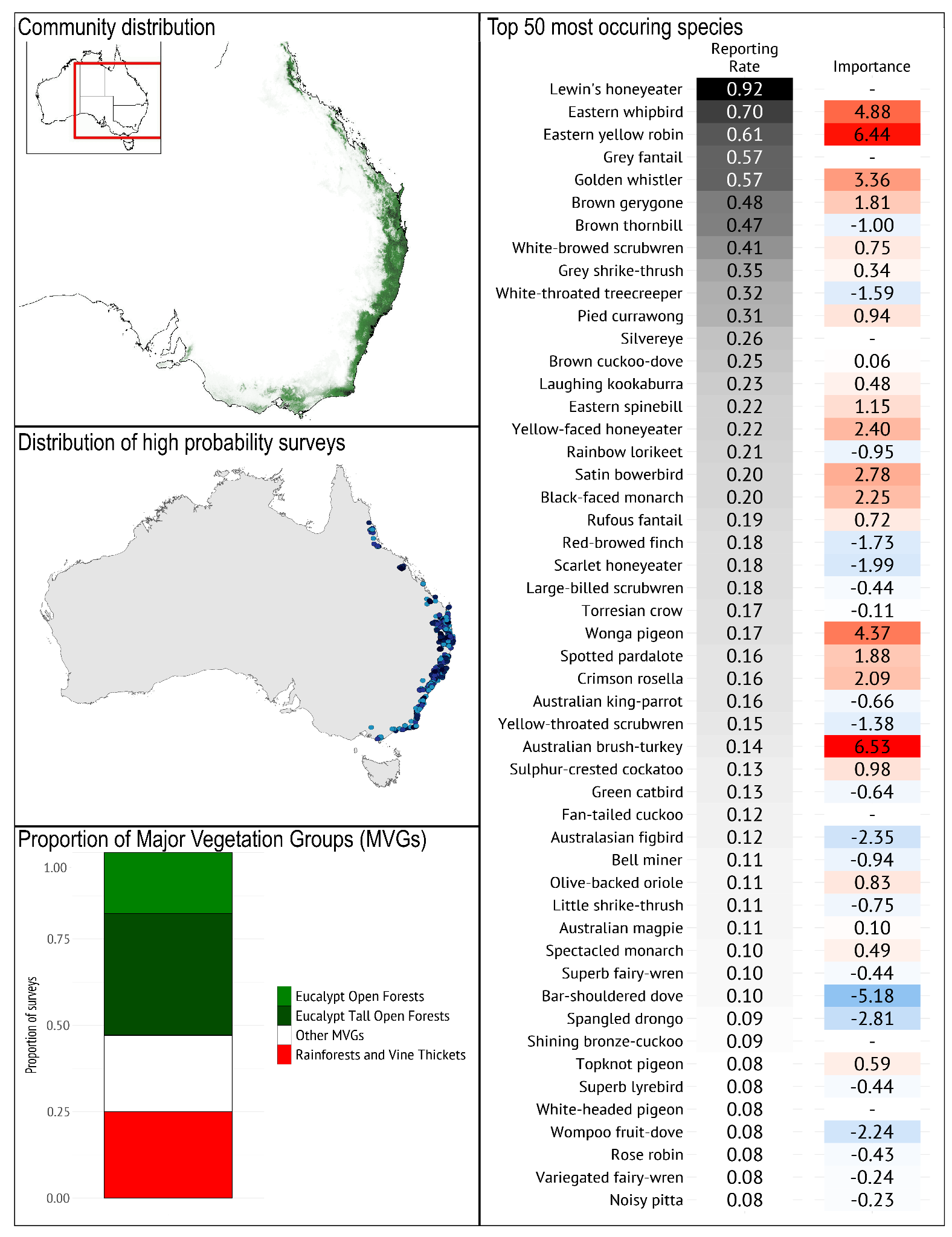

***Geographical occurrence***

The *Eastern Wet Forest* bird community typically occurs in the more mesic, taller eucalypt-dominated forests close to the east coast, along the eastern/southern slopes of the Great Dividing Range, typically in cool to warm temperate areas. It occurs on mainland Australia and is distinct from those in similar vegetation in Tasmania.

***Species composition***

*Typical species*: It is typified by high prevalence of Lewin’s Honeyeater, Eastern Whipbird, Eastern Yellow Robin, Golden Whistler, White-browed Scrubwren, and Grey Fantail. The species which are more likely to occur in this community than the other communities in this region include Eastern Whipbird, Eastern Yellow Robin, Wonga Pigeon, Satin Bowerbird; it largely lacks the High-elevation Tropical Rainforest endemics (though some typical rainforest species intrude along the ecotone between *Eastern Wet Forest* and *High Elevation Tropical Rainforest* communities (Chapman & Kofron 2010) and coastal woodland species such as White-cheeked Honeyeater.

*Guild structure:* The bird community is dominated by insectivores, small to medium-sized nectarivores and frugivores with a relatively high proportion of ground- and shrub-foraging insectivores (Milledge & Recher 1985; Recher et al. 1985; Chapman & Kofron 2010). Large forest predators including those reliant on tree hollows for nesting occur (e.g. Powerful Owl, Sooty Owl) but are not captured well by standard 2-ha 20-min surveys so aren’t reflected well in the reporting rates and importance scores presented here.

*Temporal dynamics*: The composition of the *Eastern Wet Forest* community includes a group of long-distance migratory species (e.g. Rufous Fantail, Olive-backed Oriole, Satin Flycatcher, Yellow-faced Honeyeater) that typically move south in spring and return north in late summer (Kikkawa 1968; Griffioen & Clarke 2002). The seasonal composition of the community is also influenced by local altitudinal migrants such as Brown Gerygone and Black-faced Monarch (Robinson 1993). Together these species are an important seasonal (spring-summer) component of the bird communities (Recher et al. 1983). Moderately high inter-annual variation in climatic conditions and food availability drive interannual dynamism of several honeyeater species (Mac Nally 1995; Mac Nally & Timewell 2005).

*Geographical variation*: In the far north, species that occur in the ecotone between the *High Elevation Tropical Rainforest* and *Eastern Wet Forest* communities appear in this community e.g. Mountain Thornbill, Bridled Honeyeater (Chapman & Kofron 2010).

***Vegetation/habitat associations***

The *Eastern Wet Forest* bird community is most prevalent in Eucalypt open forests, Eucalypt tall open forests and rainforest and vine thicket Major Vegetation Groups (MVGs). There was no separate bird community identified for the lowland rainforests which is consistent with the literature (Recher et al. 1991). Common dominant canopy species include Mountain Ash (*Eucalyptus regnans)* in Victoria’s central Highlands, Flooded Gum *Eucalyptus grandis* and a mix of sclerophyllous tree species - e.g., Red Stringybark, (*Eucalyptus resinifera*), Turpentine (*Syncarpia glomulifera*) and Swamp Box (*Lophostemon suaveolens*) in wet sclerophyll forests of tropical north-east Queensland. The community can also occur in the drier foothill forests adjacent to wetter areas. A high degree of structural complexity of vegetation typifies this community, and the typical species of this community require dense, low shrub cover (e.g. Eastern Whipbirds and White-browed Scrubwrens), trees with rough or decorticating bark for White-throated Treecreepers, and diverse forest strata for many species, for example Grey Fantail, Lewin’s Honeyeater, Golden Whistler, Brown Thornbill, that glean, sally or probe amongst the subcanopy. Timber harvesting and fire influence the structural and compositional characteristics of this community and succession following these disturbances may result in major variation in bird composition (Serong & Lill 2012).

#### **High Elevation Tropical Rainforest**

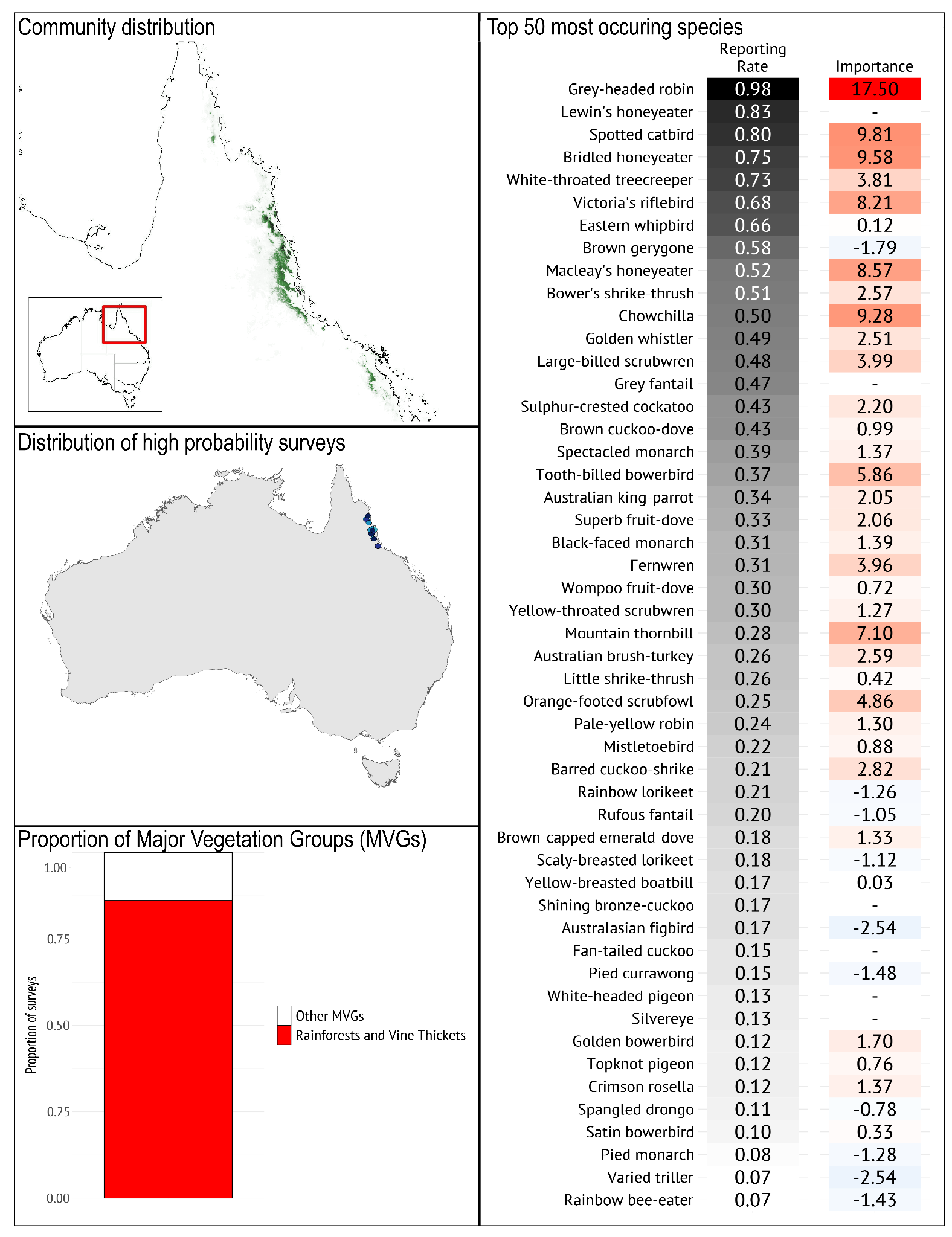

***Geographical occurrence***

The *High Elevation Tropical Rainforest* bird community highly restricted community occurs only in the high elevation rainforest areas of the wet tropics bioregion, far north Queensland.

***Species composition***

*Typical species*: The community is typified by the prevalence of Grey-headed Robin, Lewin’s Honeyeater, Spotted Catbird, Chowchilla, Bridled Honeyeater and White-throated Treecreeper. The species whose presence makes it more likely for a bird list to be deemed part of the *High Elevation Tropical Rainforest* bird community than the other communities in the region include Grey-headed Robin, Chowchilla, Spotted Catbird, Bridled Honeyeater, Victoria’s Riflebird, and Macleay’s Honeyeater. This community includes a large number of range-restricted species that only occur at high elevations and are increasingly threatened by the warming climate (De La Fuente et al. 2023).

*Guild structure*: This community has a particularly high proportion of frugivores and ground-foraging insectivores.

*Temporal dynamics*: The composition of this community is relatively stable seasonally, dominated by residents, but it is changing with climate and its boundaries are shifting to higher elevations; lowland species are also becoming more common (Williams & De La Fuente 2021).

*Geographical variation*: As this community occurs over a very small area primarily above ~800 m elevation there is little geographical variation; elevation is the primary determinant of composition (Williams et al. 2010).

***Vegetation/habitat associations***

The *High Elevation Tropical Rainforest* bird community is most prevalent in the rainforest and vine thicket Major Vegetation Group, and occasionally occurs in ecotonal areas with Eucalypt Tall Open Forest (Kutt & Vanderduys 2017).

### **Region A3: South-eastern woodland**

The South-eastern woodland region consists of four communities, *Eastern Heathy Woodland*, three *South-eastern Woodland* communities (*Coastal and Mid-elevation, Cool Temperate Lowland, Dry Lowland*). The climate envelope of the three different *South-eastern Woodlands* communities differs by elevation, rainfall and temperature, they are similar in species composition and also in the break-down of occurrence in different Major Vegetation Groups (MVGs). The *Eastern Heathy Woodland* bird community had lower prevalence in the Eucalypt open and tall open forests MVGs than the three *South-eastern Woodland* communities in this region, and correspondingly higher prevalence in Eucalypt open woodland, heathlands, and ‘other shrublands’ MVGs. All of the communities in this region had high prevalence in Eucalypt woodlands. The composition of the bird communities comprising this region and Region A2 (eastern wet forests and coastal heath) includes a group of migratory species (e.g. Yellow-faced Honeyeater, White-naped Honeyeater, Fan-tailed Cuckoo, Grey Fantail) that fly south along the eastern coastal regions to breed in the spring and then north to escape the southern winter. These species become an important seasonal (spring-summer) component of the region.

Landscape-scale fires can remove key resources needed by the communities making up the South-eastern Woodlands region. Some species (e.g. White-throated Treecreeper, Golden Whistler) may be reliant upon unburnt refuges in adjacent more mesic MGVs (e.g. Open Tall Forests, Rainforests) during the decades following a fire, from which these species will eventually recolonise the less mesic, burnt woodland habitats as they recover (Robinson et al. 2014, 2016).

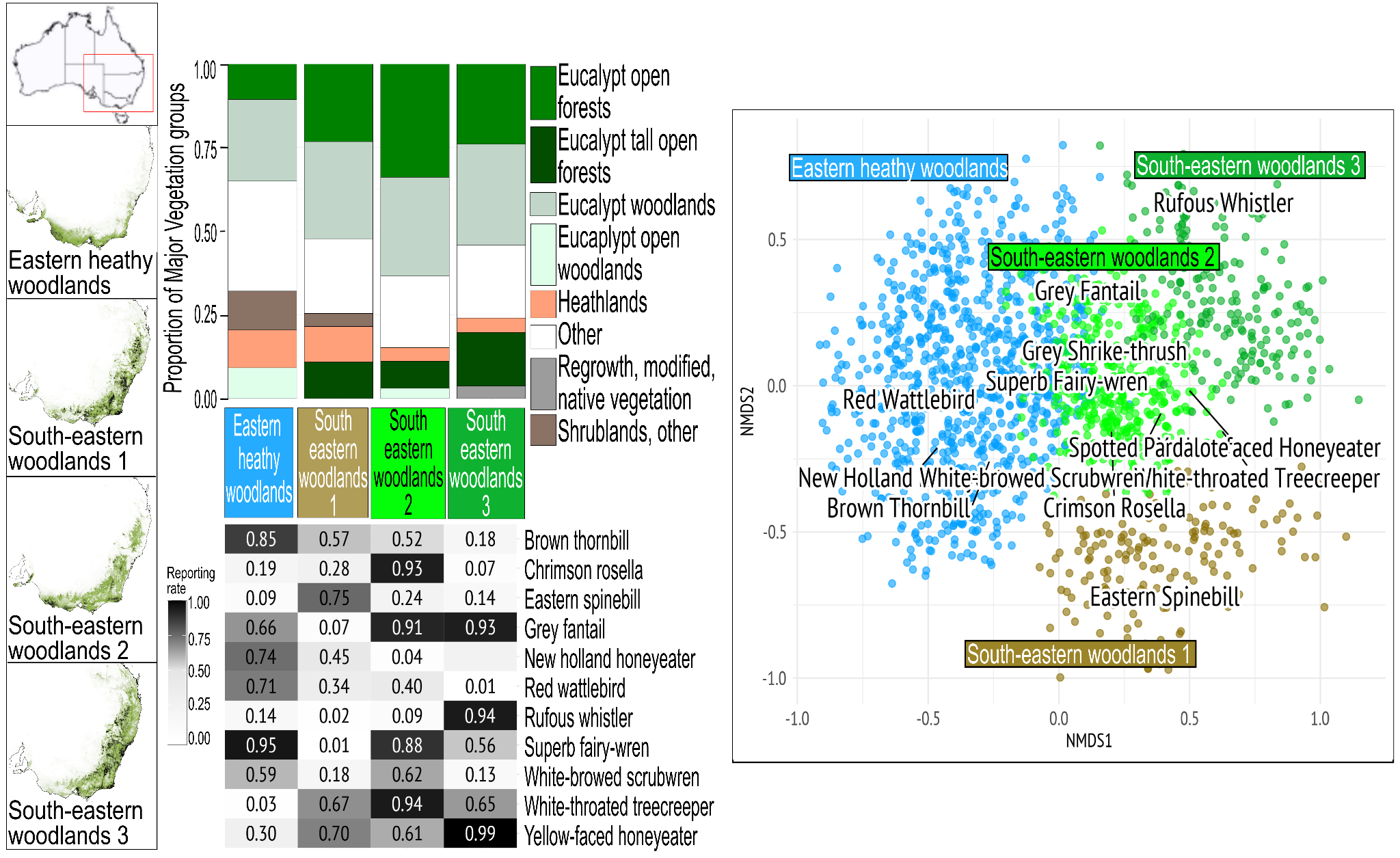

**Figure S1.3** From left to right: far left) the top map indicates location of region, the bottom maps show distribution of communities, darker shading represents higher likelihood of occurrence. Middle, top is the comparison of proportion of Major Vegetation Groups (MVG) and bottom is the comparison of Reporting Rates (RR) of the top 5 most commonly occurring species from each community. Far right, a non-metric multidimensional scaling (NMDS) plot depicting the different important species distinguishing communities.

#### **Eastern Heathy Woodland**

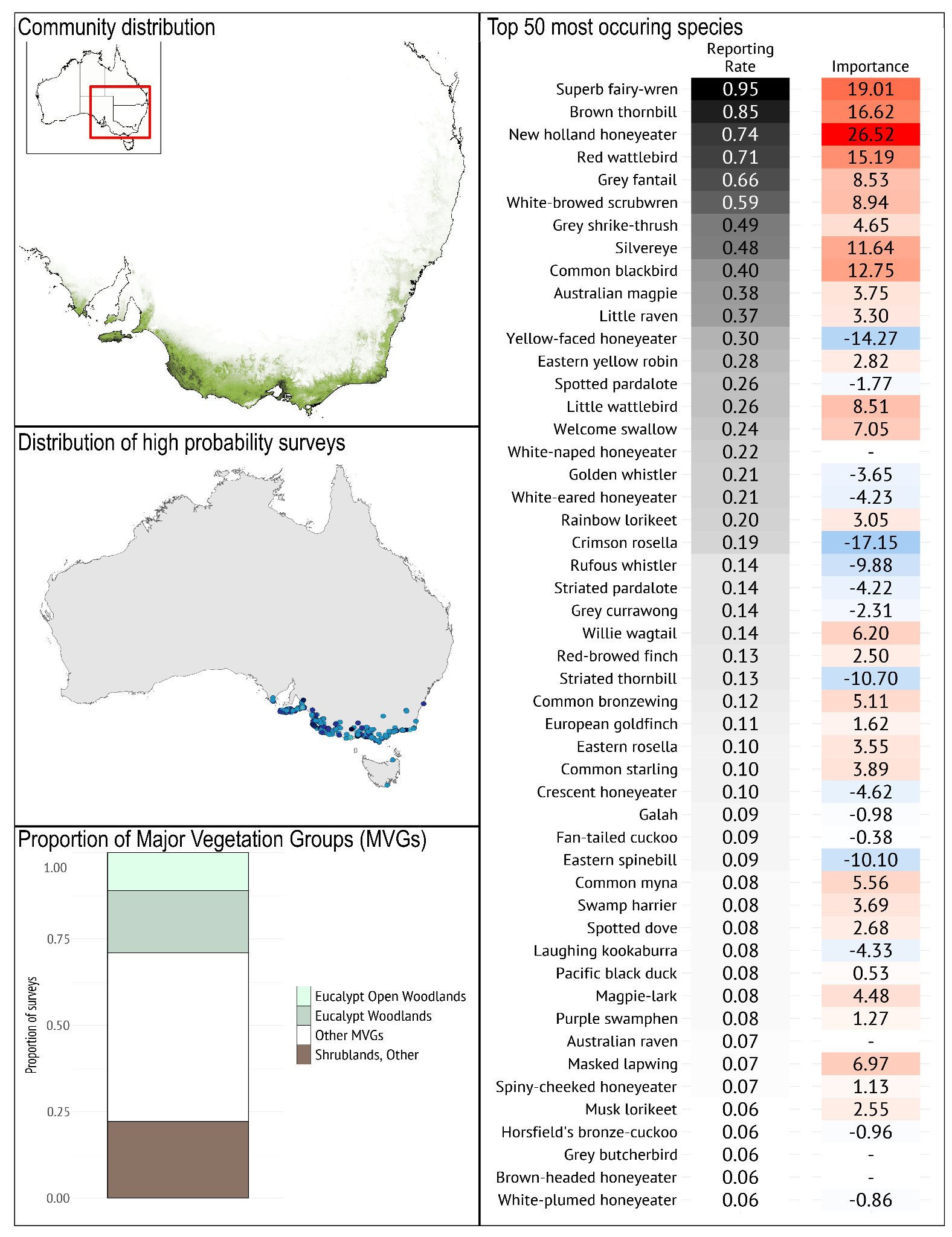

***Geographical occurrence***

The *Eastern Heathy Woodland* bird community occurs in heathlands and heathy forests and woodlands in cool areas (with annual temperatures near 15-16 degrees) along the south east fringe of Australia. The community runs from the Eyre peninsula, around the bottom of Victoria and up the east coast into New South Wales.

***Species composition***

*Typical species:* It is typified by the prevalence of Superb Fairy-wren, Brown Thornbill, New Holland Honeyeater, Red Wattlebird and Grey Fantail. It distinguishes itself from other communities in this region based on the higher likelihood of Brown Thornbill, New Holland Honeyeater, Red Wattlebird, Superb Fairy-wren occurring. The White-browed Scrubwren is also more likely to occur in this community, though also in the *Cool Temperate Lowland South-eastern Woodland* bird community. Equally important in distinguishing the community is the low likelihood of White-throated Treecreeper and Weebill occurring.

*Guild structure:* The community is distinguished from the other communities in this region by the abundance of the nectarivores Red Wattlebird and New Holland Honeyeater, and by the prevance of small understorey-dependent insectivores, nsectivores that forage at a range of heights and substrates but especially in the shrub layer.

*Temporal dynamics:* Eastern heathy woodlands experience significant fires, after which the Eucalypts and grass trees resprout, triggering mass flowering events (e.g. of Xanthorrhoea spp.) that attract a suite of nectarivore species, often at high abundance (Rainsford et al. 2022).

*Geographic variation:* Most species that typify this community occur throughout its extent, though with local variation e.g. Crescent Honeyeaters tend to occur in more mesic examples of this community type.

***Vegetation/habitat associations***

The *Eastern Coastal Woodland* bird community is most prevalent in the ‘other shrublands’ and Eucalypt woodlands Major Vegetation Groups (MVGs). *Eastern Heathy Woodland* communities overlap in bird species composition with other woodland communities depending upon the level of cover provided by emergent trees (e.g. Eucalypts or Allocasuarinas); increasing the likelihood of species associated with trees being present (e.g. Rufous Whistler, White-troated Treecreeper), and diminishing likelihood of species more associated with treeless heaths (e.g. Striated Fieldwren, Southern Emu-wren).

#### **Coastal and Mid-Elevation South-eastern Woodland**

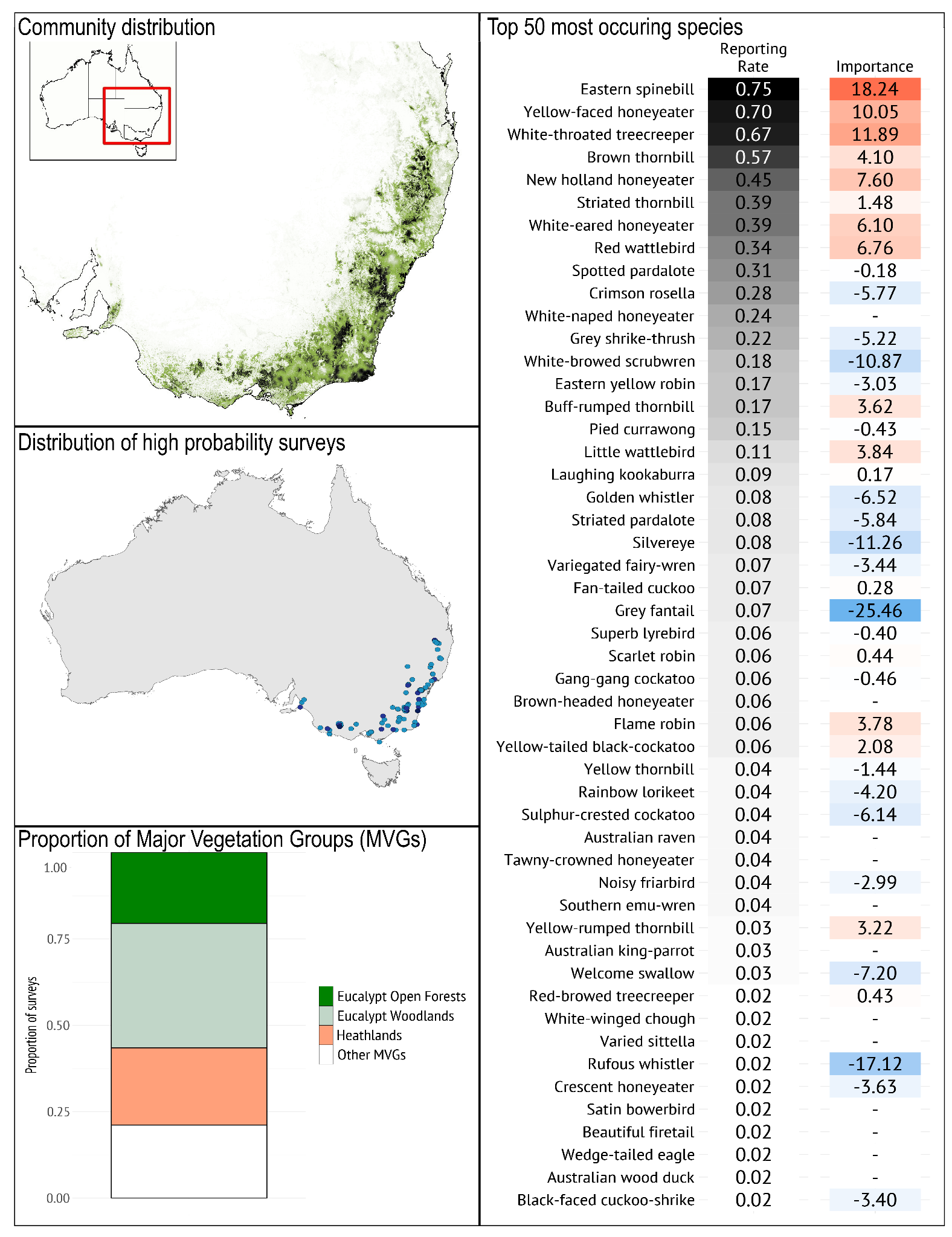

***Geographical occurrence***

The *South-eastern Woodland 1* bird community occurs in the woodlands of south-eastern Australia, primarily associated with the ridges and peaks (though not alpine or subalpine areas) along the great dividing range, occurring in areas which have higher rainfall than the other communities in this region. It stretches from Kangaroo island and Flinders Ranges in South Australia, across through the Grampians, Kara Kara, Lerderderg parks and across Victoria’s inland slopes before joining the main extent of the great dividing range and continuing up to the northern New England Tableland bioregion.

***Species composition***

*Typical species*: The *Coastal and Mid-Elevation South-eastern Woodland* bird community is typified by the prevalence of Eastern Spinebill, Yellow-faced Honeyeater, White-throated Treecreeper, Brown Thornbill, and New Holland Honeyeater. The community is distinguished from the other south-eastern woodland communities by the high likelihood of finding Eastern Spinebill and lower likelihood of finding Grey Fantail, Rufous Whistler, and Superb Fairy-wren. This community contains more of the typical wetter forest species like spinebills and brown thornbills than the other communities in this region.

*Guild structure*: The community is dominated by small to medium insectivores and nectarivores of all sizes. Lower in reporting rate but still important are forest cockatoos (Gang-gang Cockatoo and Yellow-tailed Black-Cockatoos, with the latter often occurring near pine plantations which they exploit as a food source, or in heathy woodland where they take woody seeds of native plants such as Hakea and Banksia spp.).

*Temporal dynamics*: The community is characterised by high seasonal turnover, with approximately 50% of breeding residents absent during the winter (Recher et al. 1983). Reporting rates of nectarivores (Eastern Spinebill, New Holland Honeyeater, Red Wattlebird, Yellow-faced Honeyeater) also vary substantially as they track floral resources.

*Geographical variation*: Most species typical of this community occur throughout its extent. Exceptions include Variegated Fairy-wren (absent from southern parts), Little Wattlebird (confined to heathy woodland, especially near coast) and New Holland Honeyeater (very common in gardens and heathy woodland, but scarce in forests elsewhere).

***Vegetation/habitat associations***

The *South-eastern Woodland 1* bird community is most prevalent in eucalypt woodlands with sparse mixed proteaceous understory, heathlands and eucalypt open forests Major Vegetation Groups (MVGs).

#### **Cool Temperate Lowland South-eastern Woodland**

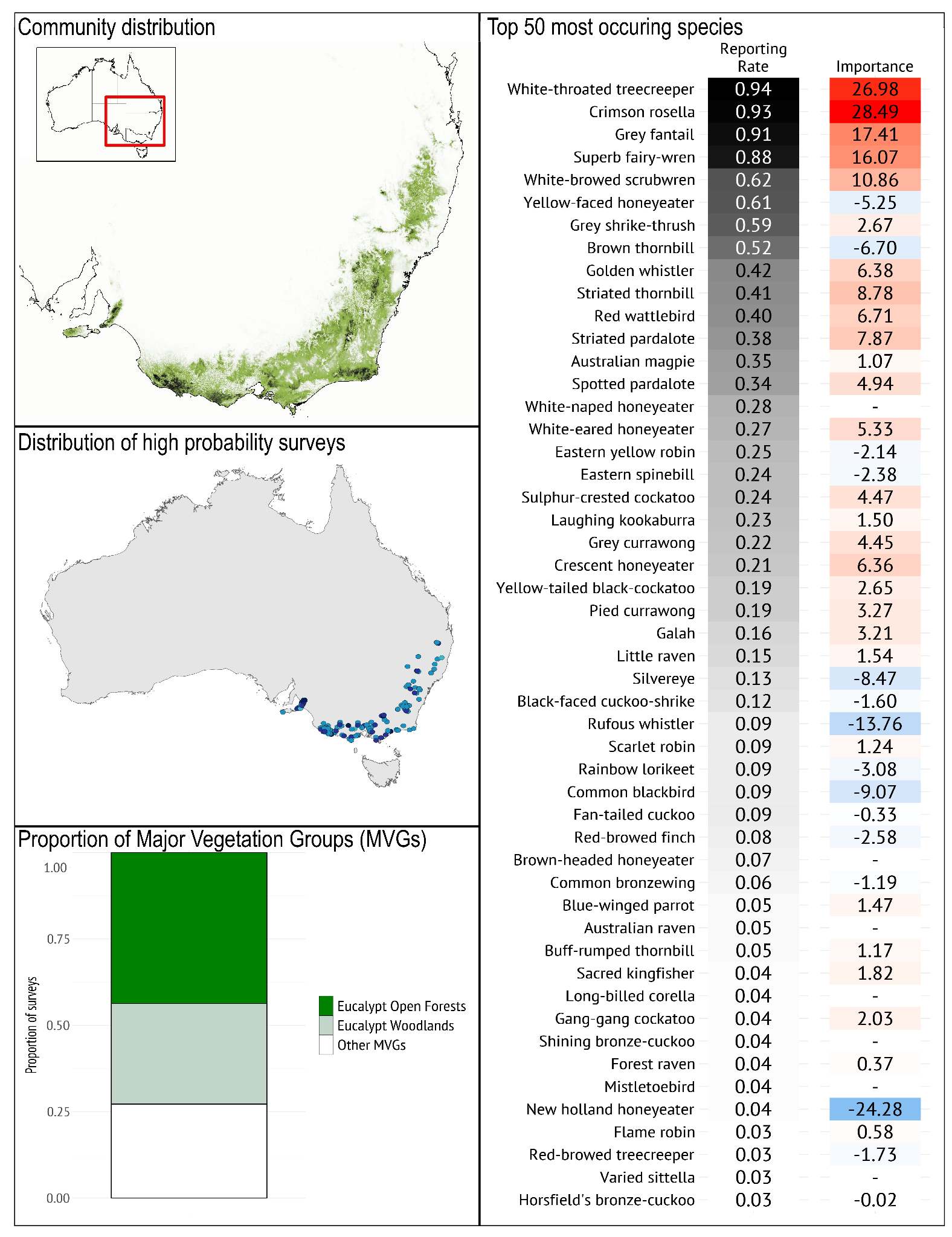

***Geographical occurrence***

The *Cool Temperate Lowland South-eastern Woodland* bird community occurs in more fertile low lying hills, in cooler areas of south-eastern Australia. The western extent of the community is in the Flinders Ranges and Kangaroo Island, then there is a gap in the distribution which resumes in the Naracoorte area before sweeping in a thick band around south-eastern Australia, through the Strzelecki Ranges (Radford & Bennett 2005), east Gippsland, eastern slopes of the Great Dividing Range, and as far north as Brisbane.

***Species composition***

*Typical species*: The *Cool Temperate Lowland South-eastern Woodland* bird community is typified by the prevalence of White-throated Treecreeper, Crimson Rosella, Grey Fantail, Superb Fairy-wren, and White-browed Scrubwren. The community is best distinguished from the other communities in this region by the prevalence of White-throated Treecreeper and Crimson Rosella, and the low probability of finding the New Holland Honeyeater or Rufous Whistler.

*Guild structure*: The community is dominated by small and medium insectivores, including both pardalotes which tend to be associated with Symphomyrtus eucalypts (Woinarski 1985 p. 19). The community also includes many hollow-nesting species, with a substantive representation of mesopredators such as Australian Magpies, Laughing Kookaburra, currawongs and corvids.

*Temporal dynamics*: The *Cool Temperate Lowland South-eastern Woodland* community is dominated by a suite of largely sedentary species, presumably as a function of relatively consistent availability of arthropod food resources. It supports a smaller number of local blossom nomads than *Coastal and Mid-elevation South-eastern Woodland*.

*Geographical variation*: The species of mesopredators typifying the community, varies across its range with Little, Australian or Forest Raven occurring and Pied and Grey Currawong occurring depending on location.

***Vegetation/habitat associations***

The *Cool Temperate Lowland South-eastern Woodland* bird community is most prevalent in Eucalypt open forests and Eucalypt woodlands Major Vegetation Groups (MVGs). Some of the small insectivores are tied to the presence of some understory particularly along gullies and drainage lines (scrubwrens, thornbills, yellow robins, fairy-wrens). The presence of sufficient tree cover of varying ages would support the treecreepers and rosellas. This community is also representative of woodlands with a sparse understory and where there might be some bigger trees.

#### **Dry Lowland, South-eastern Woodland**

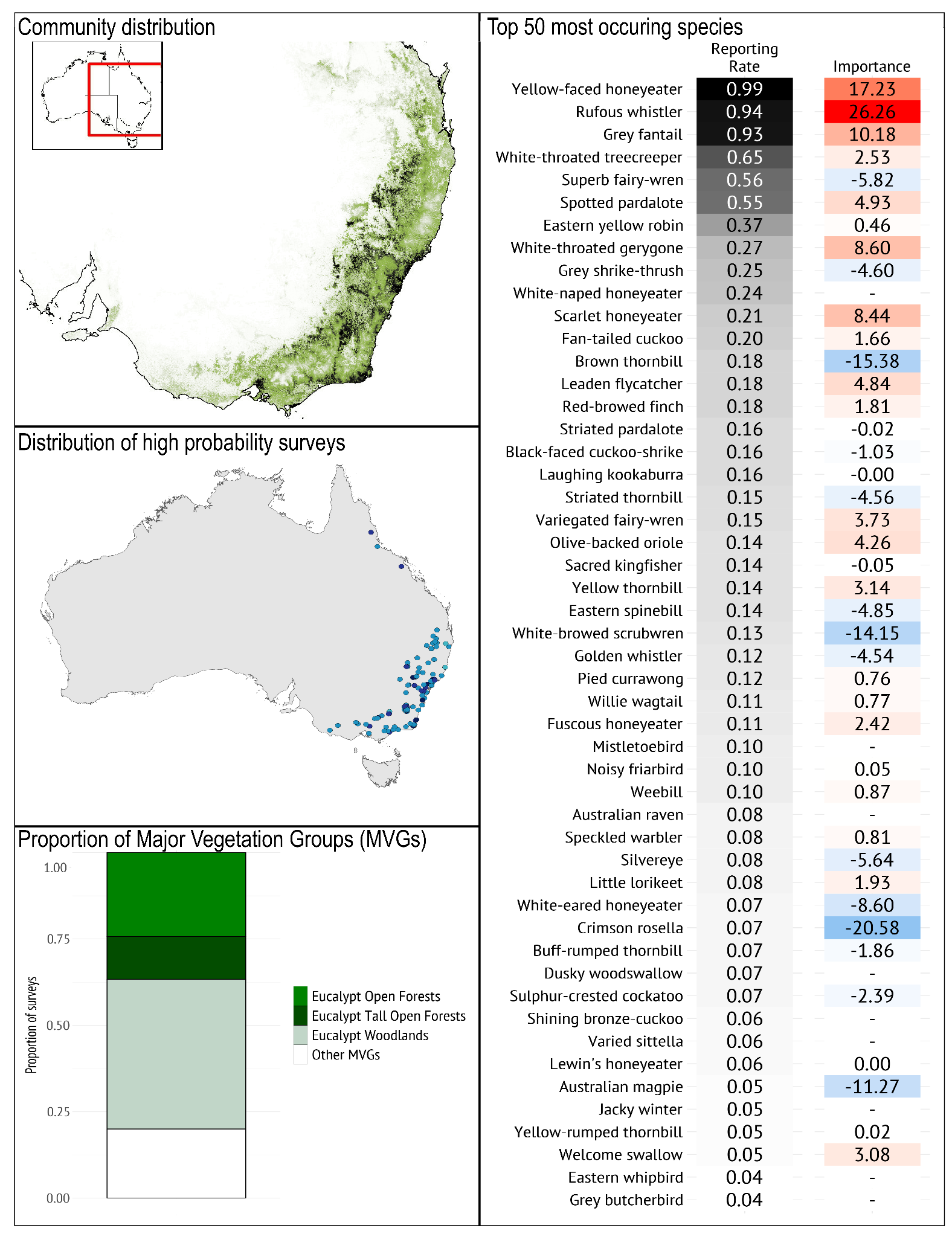

***Geographical occurrence***

The *Dry Lowland South-eastern Woodland* bird community is most likely to occur in warmer and drier, low lying areas than the other south-eastern woodland bird communities. There is a slight chance of this community occurring as far to the west as the Flinders Ranges but the areas with the highest probability of occurrence are south-east suburbs of Melbourne, Gippsland Plains, the Nooramunga Marine and Coastal Park, the ACT, Croajingalong National Park, greater Sydney area, and NSW central tablelands. The composition of this community is similar to that of the *South-eastern Open Box-Gum Woodland* bird community, the primary differences being in the difference in occupancy of Grey Fantails and Rufous Whistler which are much more prevalent in the *Dry Lowland South-eastern Woodland*.

***Species composition***

*Typical species*: The *Dry Lowland South-eastern Woodland* bird community is typified by the prevalence of Yellow-faced Honeyeater, Rufous Whistler, Grey Fantail, White-throated Treecreeper, and Superb Fairy-wren (Radford & Bennett 2005).

*Guild structure*: The community is dominated by small to medium insectivores, with a relatively low diversity of nectarivores compared with the other south-east woodland communities. Nest parasites were present at higher reporting rates than the other two south-eastern woodland bird communities.

*Temporal dynamics*: This community is characterised by the abundance of summer migratory species responding to the relatively high seasonal abundance of arthropods. These include Sacred Kingfisher, Olive-backed Oriole, Leaden Flycatcher, White-throated Gerygone and cuckoos. There may also be irruptions of migratory or nomadic nectarivores including Scarlet and Yellow-faced Honeyeater, and typical summer migrants including Sacred Kingfisher, Olive-backed Oriole, and cuckoos.

*Geographical variation*: The species that typify this community occur throughout its extent.

***Vegetation/habitat associations***

Like the *Cool Temperate Lowland South-eastern Woodland* bird community, The *Dry Lowland South-eastern Woodland* bird community is most prevalent in Eucalypt open forests and Eucalypt woodlands Major Vegetation Groups (MVGs).

### **Region B: Northern Australia**

The Northern Australia region consists of five bird communities: *Eastern Tropical Forest and Monsoon Thicket*, *Eastern Tropical and Subtropical Woodland, Northern Tropical Forest and Monsoon Thicket*, *Northern Savanna,* and *Tiwi Islands and Cape York*. Eucalypt woodlands and open forests dominate this region, with patches of Acacia, Melalauca and Casuarina Major Vegetation Groups (MVGs) throughout. The *Eastern* and *Northern Tropical Forest and Monsoon Thicket* bird communities differ from the others in that they occur in rainforests and vine thickets. The vast majority of the *Tiwi Islands and Cape York* bird community comprises a single MVG, Eucalypt open forests, while the other bird communities are spread more evenly across multiple MVGs.

Northern Australia is characterised by highly seasonal rainfall patterns, whereby rain predominantly falls between November and April (wet season), while the months of May to October receive little or no rain (dry season). There are rainfall gradients apparent across the five sub-communities and broad habitat types across the region, with the greatest rainfall occurring in coastal areas and rainfall declining north to south (Spessa et al. 2005; Hutley et al. 2011). Average annual rainfall totals range from >2000 mm in coastal areas (e.g. areas occupied by the *Tiwi Island and Cape York* bird community) to <600 mm in the more xeric, southern parts, inhabited by the *Northern Savanna* bird community, in which interannual rainfall can vary greatly.

Fire plays a major role in maintaining vegetation patterns across the region, with fire most frequent in open savannas, somewhat less frequent in other dry regions and very infrequent in vine thicket areas (Spessa et al. 2005). Though often characterised as a singular intact community (e.g., northern Australian tropical savannas) there is remarkable variation in topography, land use and land types, which is more apparent in Queensland, compared to the more uniform Top End savannas. In general, differences in the structural complexity, dominant tree species and amount of canopy cover present across Northern Australia influence the bird community assemblages found in each. What is often overlooked is the presence of Noisy Miners in many of the east coastal and upland savanna communities in Queensland, which gives way to Yellow-throated Miners across much of the remainder of northern Australia (Eyre et al. 2009; Mac Nally et al. 2014; Kutt et al. 2016). This, coupled with more intensive land use in Queensland, has had a transformative effect on some bird communities (Hernandez et al. 2024). The patterns of birds in this region and then each community, though in some respects fairly uniform, reflect this subtle variation, and the typical determinants of bird occurrence and composition. For example, in open savannas with lower rainfall there are more generalists (e.g., Peaceful Dove, Willie Wagtail, Magpie-lark), while where there is more structurally complex vegetation with higher canopy cover on the east coast, more perching and gleaning species are apparent (e.g., Little Shrike-thrush, Varied Triller).

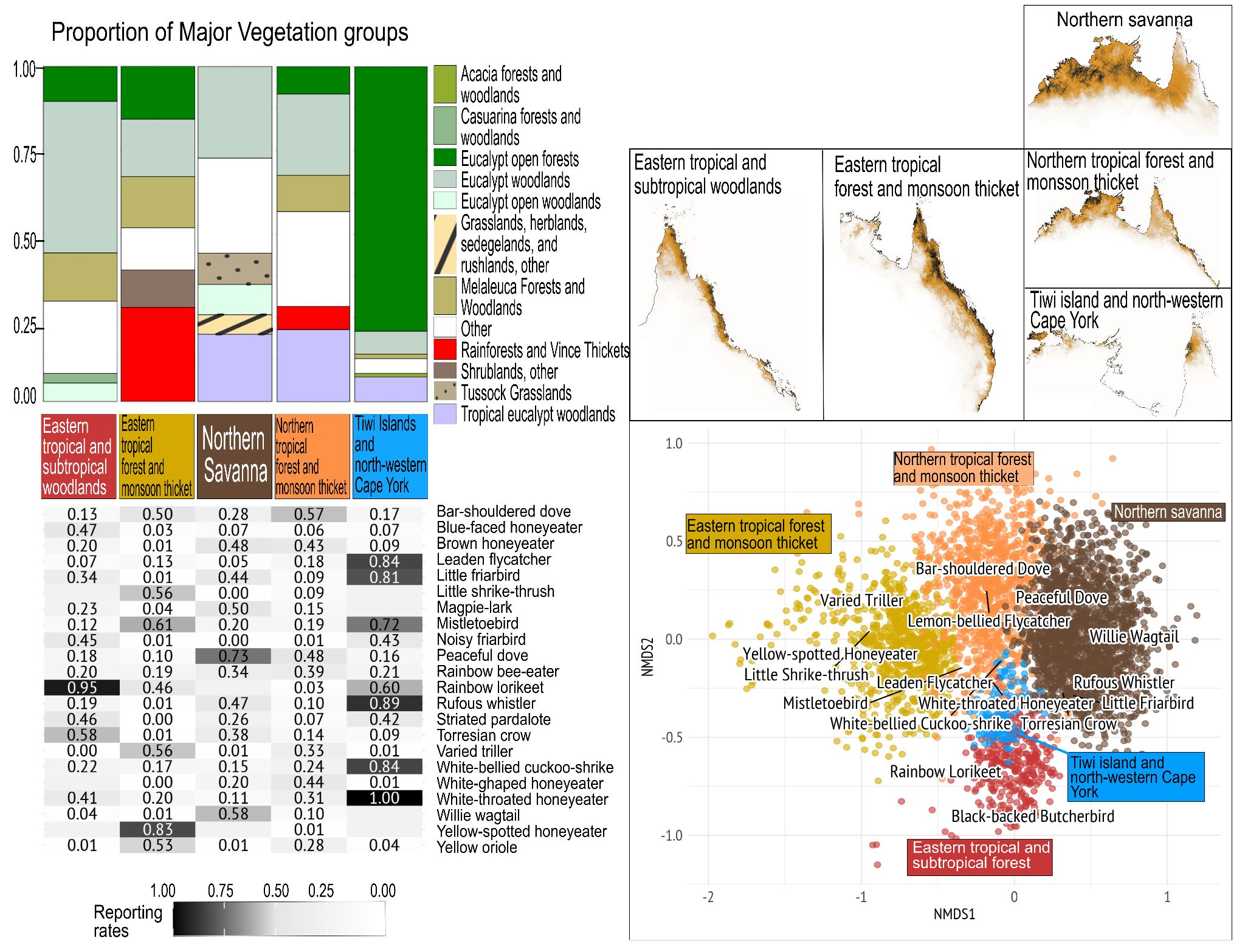

**Figure S1.4** Top left, the comparison of proportion of Major Vegetation Groups (MVG), bottom left is the comparison of Reporting Rates (RR) of the top 5 most commonly occurring species from each community. The right maps show distribution of communities, darker shading represents higher likelihood of occurrence, and the bottom right shows a non-metric multidimensional scaling (NMDS) plot depicting the different important species distinguishing communities.

#### **Eastern Tropical Forest and Monsoon Thicket**

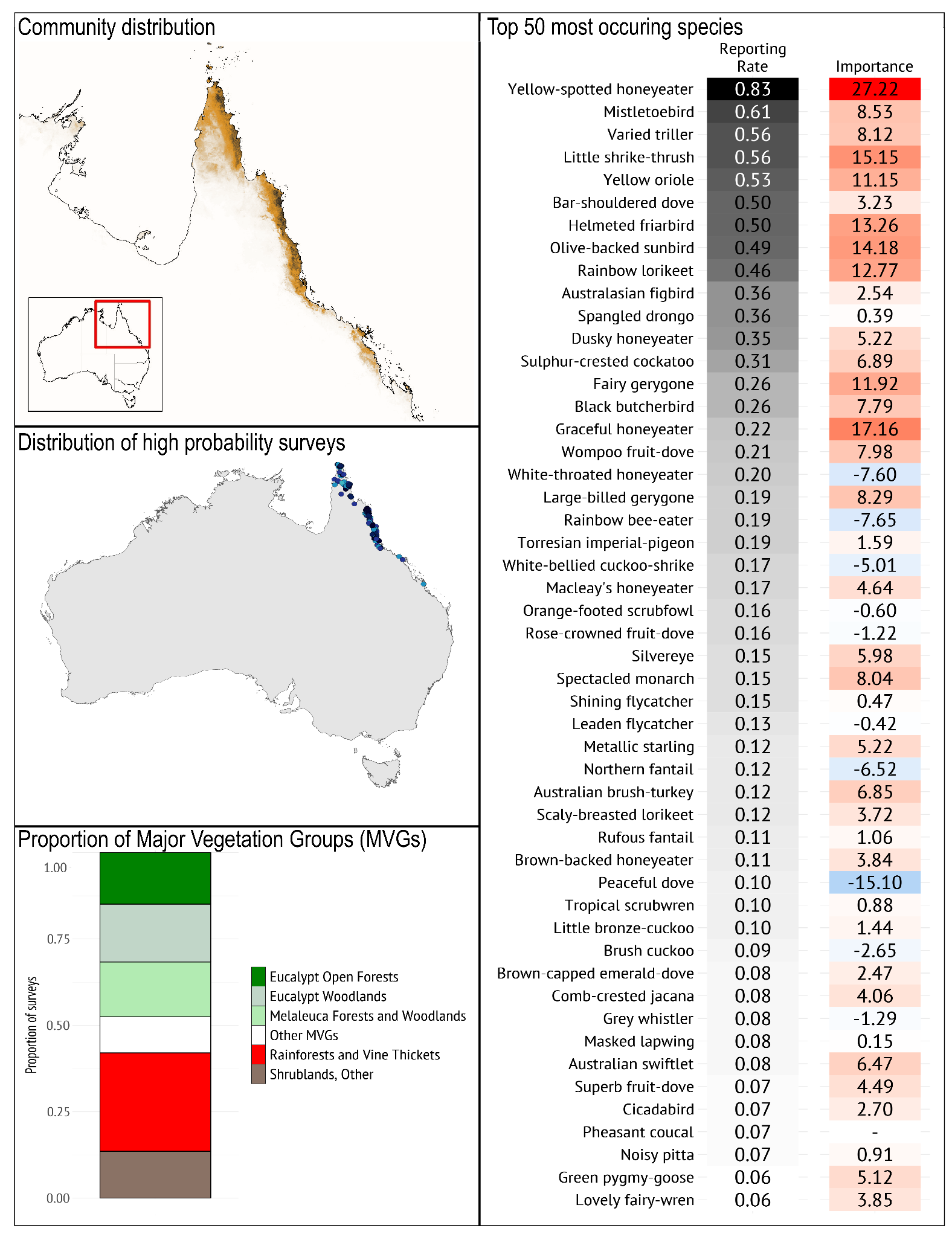

***Geographical occurrence***

The *Eastern Tropical Forest and Monsoon Thicket* bird community occurs on the Cape York Peninsula across the northern third of the peninsula north of Coen and then running down the east coast to Clairview. The extent covers numerous national parks including Girringun, Wooroonooran, Gadgarra, Malbon Mount Lewis, Daintree, and Jardine River.

***Species composition***

*Typical species*: It is typified by the high prevalence of Yellow-spotted Honeyeater, Mistletoebird, Varied Triller, Little Shrike-thrush, and Yellow Oriole. The *Eastern Tropical Forest and Monsoon Thicket* bird community is best distinguished from the other communities in the region by the prevalence of Yellow-spotted, Graceful Honeyeater, Olive-backed Sunbird and Little Shrike-thrush (which distinguishes it from the *Northern Tropical Forest and Monsoon Thicket* community largely as these species do not occur in the Northern Territory), Yellow Oriole, and Varied Triller. Notably, it also has lower reporting rates than other communities in this region for Brown Honeyeater, Little Friarbird, Magpie Lark, Rufous Whistler, Striated Pardalote, Torresian Crow, White-gaped Honeyeater and Willie Wagtail, as these are species that occur across the rainfall gradient.

*Guild structure*: As this community occurs predominantly in rainforest and vine thicket MVGs, the *Eastern Tropical Forest and Monsoon Thicket* bird community is characterised by relatively high reporting rates of frugivores (e.g., Yellow Oriole, Mistletoebird) and nectarivores (e.g., Yellow-spotted Honeyeater, Olive-backed Sunbird, fruit pigeons).

*Temporal dynamics*: A relatively temporally stable community but augmented by summer breeding migrants such as Metallic Starling.

*Geographical variation*: Most species that typify this community occur throughout its range but some are more likely to occur in denser or more coastal locations, e.g., Black Butcherbird, Large-billed Gerygone.

***Vegetation/habitat associations***

The *Eastern Tropical Forest and Monsoon Thicket* bird community occurs near this coast, most commonly in rainforests and vine thickets, but also in Eucalypt woodlands, Melaleuca forests and woodlands, and Eucalypt open forests Major Vegetation Groups (MVGs).

#### **Eastern Tropical and Subtropical Woodland**

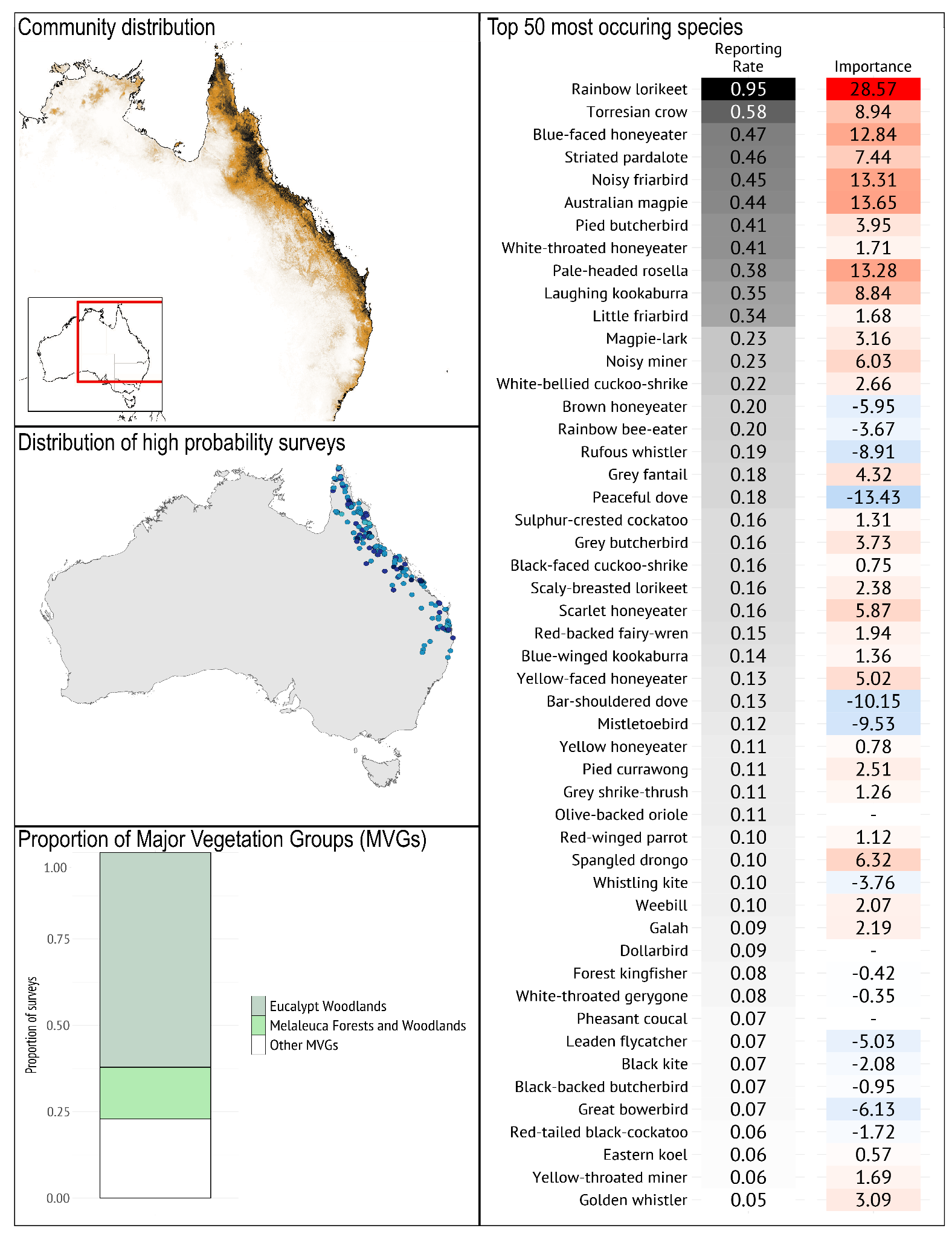

***Geographical occurrence***

The *Eastern Tropical and Subtropical Woodland* bird community extends further inland than does the *Eastern Tropical Forest and Monsoon Thicket* community, including on the inland slopes of the Great Dividing Range, throughout the tropical and subtropical zones. The northern-most occurrences of the community include the drier woodlands of Cape York Peninsula, and south to approximately central New South Wales. Its modelled distribution occurs as far west as Katherine in the Northern Territory.

***Species composition***

*Typical species*: *Eastern Tropical and Subtropical Woodland* is typified by the prevalence of Rainbow Lorikeet, Torresian Crow, Blue-faced Honeyeater, Striated Pardalote, and Noisy Friarbird. The community is best distinguished from other communities in the region by the high reporting rate of Rainbow Lorikeet (rather than Red-Collared Lorikeet), Torresian Crow, and Blue-faced Honeyeater, Australian Magpie and Pale-headed Rosella, and is less likely to include Yellow-spotted and Yellow-tinted Honeyeater, Olive-backed Sunbird, and Yellow Oriole.

*Guild structure*: The community is dominated by larger-bodied insectivores/carnivores and nectarivores, with relatively few granivores, frugivores and smaller-bodied insectivores. Aggressive and territorial Noisy or Yellow-throated Miners can dominate, which may reflect why the dominant birds are large-bodied rather than small-bodied. The dominance of Noisy Miners (in the south of this community) and Yellow-throated Miners (in the north and west of this community is exacerbated by broadscale clearing of habitat and increased timber harvesting activity (Eyre et al. 2009; Mac Nally et al. 2014; Kutt et al. 2016).

*Temporal dynamics*: Many of the honeyeaters, including friarbirds and smaller species, are migrants/nomads, contributing to the seasonal variation in this community. They will also occur in the structurally more complex components of this community.

*Geographical variation*: Some species only occur toward the northern part of the community’s extent, such as Yellow Honeyeater, Blue-winged Kookaburra and Great Bowerbird.

***Vegetation/habitat associations***

The *Eastern Tropical and Subtropical Woodland* bird community occurs primarily in Eucalypt woodlands and Melaleuca forests and woodlands Major Vegetation Groups (MVGs).

#### **Northern Tropical Forest and Monsoon Thicket**

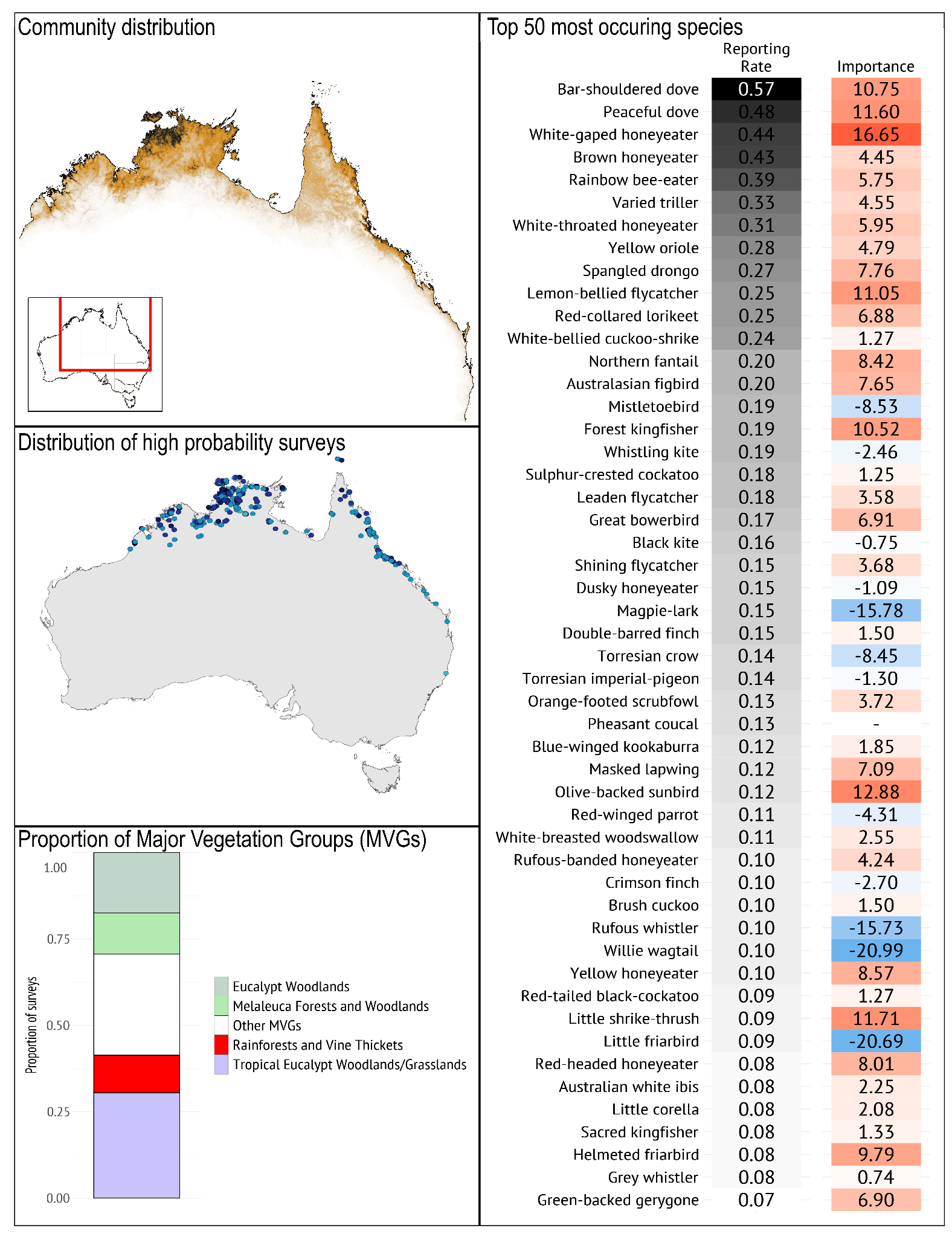

***Geographical occurrence***

The *Northern Tropical Forest and Monsoon Thicket* bird community occurs from Broome in Western Australia, across the top of the Northern Territory across to Cape York Peninsula, with the likelihood of occurrence tailing off down the east coast of Queensland towards Maryborough; the Queensland part of its distribution may stem from its similarity with Eastern Tropical Forest and Monsoon Thicket, but several key species distinguish the two. The highest probability of occurrence is around the Tiwi Islands, Kakadu, and Darwin in the Northern Territory.

***Species composition***

*Typical species*: The *Northern Tropical Forest and Monsoon Thicket* bird community is typified by the prevalence of Bar-shouldered Dove, Peaceful Dove, White-gaped Honeyeater, Brown Honeyeater, and Rainbow Bee-eater, species that occur across the entire distribution of the community. The community has intermediate reporting rates of many of the species common to the region. It is best distinguished from other communities based on the higher reporting rate of Forest Kingfisher, White-gaped Honeyeater, Olive-backed Sunbird, Little Shrike-thrush, and Lemon-bellied Flycatcher.

*Guild structure*: Granivorous doves are common in the *Northern Tropical Forest and Monsoon Thicket* bird community (e.g. Bar-shouldered, Peaceful Dove), frugivores are also well represented (e.g. Yellow Oriole, Australisian Figbird), the rest of the community comprises a mix of insectivores, carnivores, and honeyeaters of different sizes.

*Temporal dynamics*: Most of the species in this community are resident, but as with most of the Northern Australian communities, the rainfall seasonality, coupled with fire history and the high number of honeyeaters, means that the relative abundance of each species will reflect the resources present at a site. Species such as the Rainbow Bee-eater seasonally migrate.

*Geographical variation*: The commonest parrot in this community is the Red-collared Lorikeet, replaced by the Rainbow Lorikeet in the east; Yellow Honeyeater and Olive-backed Sunbird occurs only in the east while Green-backed Gerygone occurs only in the Top End and Kimberley. Most other typical species occur throughout the community’s extent. Little Shrike-thrush have now been split into Arafura (west) and Rufous (east).

***Vegetation/habitat associations***

The *Northern Tropical and Subtropical Woodland* bird community occurs primarily in tropical Eucalypt woodlands/grasslands, and Eucalypt woodlands Major Vegetation Groups (MVGs).

#### **Northern Savanna**

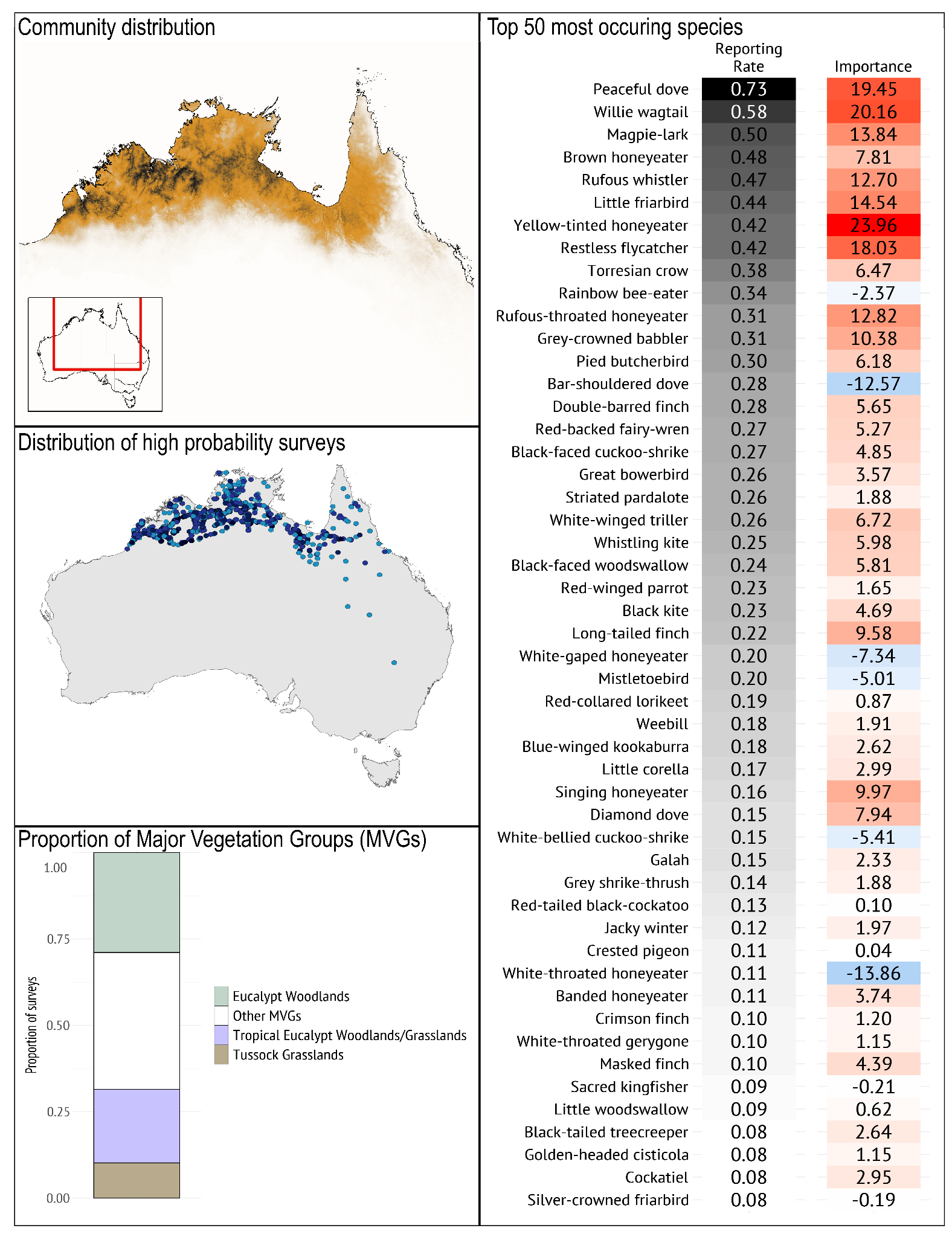

***Geographical occurrence***

The *Northern Savanna* bird community occurs throughout the northern savanna woodland zone across northern Australia; from Eighty Mile Beach in Western Australia to the Einasleigh Uplands in Queensland, extending north to the coast and into the Cape York Peninsula. The highest likelihood of occurrence is in the Wunaamin Miliwundi Ranges and the ranges around Lake Argyle in Western Australia.

***Species composition***

*Typical species*: The *Northern Savanna* bird community is typified by the prevalence of Peaceful Dove, Willie Wagtail, Magpie-lark, Brown Honeyeater, and Rufous Whistler. It is best distinguished from the other communities in the region by the higher prevalence of Yellow-tinted Honeyeater, Magpie-lark, Restless Flycatcher, and Willie Wagtail, and the lower reporting rate of Yellow Oriole, Varied Triller, Spangled Drongo, and Lemon-bellied Flycatcher.

*Guild structure*: The *Northern Savanna* bird community is dominated by species that prefer more open woodlands. This includes both small and large granivores (e.g. Peaceful Dove, Bar-shouldered Dove, Double-barred and Long-tailed Finch, Red-winged Parrot), as well as insectivores that typically forage on the ground, or from low perches (e.g. Willie Wagtail, Magpie-lark, Restless Flycatcher, Grey-crowned Babbler). There are also birds of prey (e.g. Whistling Kite, Black Kite), and a mix of resource-tracking nectarivores (Brown, Yellow-tinted, and Rufous-throated Honeyeater and Little Friarbird).

*Temporal dynamics*: The presence of honeyeaters is driven by flowering of nectar-bearing plants, and granivore dynamics are affected by fire which is a common disturbance in this community.

*Geographical variation*: As this community spans a large extent there are some species which only enter this community in the east such as Brown Treecreeper and Black-throated Finch which are replaced in the north and west by Black-tailed Treecreeper and Long-tailed Finch; others only occur on the inland fringes (e.g. Grey-fronted Honeyeater).

***Vegetation/habitat associations***

The *Northern Savanna* bird community occurs primarily in Eucalypt woodlands, and tropical Eucalypt woodlands/grasslands Major Vegetation Groups (MVGs).

#### **Tiwi Islands and Cape York**

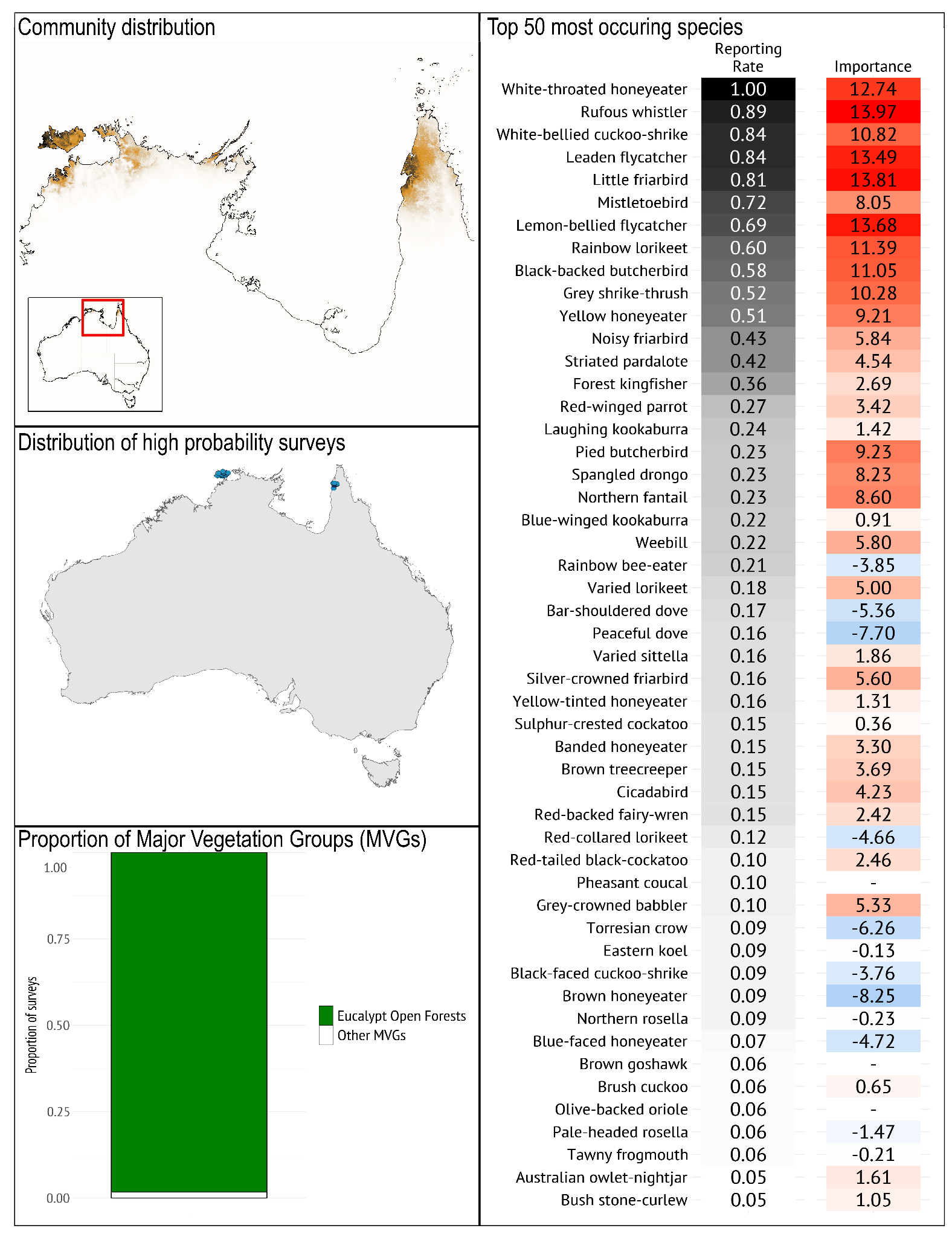

***Geographical occurrence***

The unusual distribution of the *Tiwi Islands and Cape York* bird community occurs in the Tiwi Islands and the north west portion of the Cape York Peninsula, as well as just south of the Tiwi Islands on the Cox Peninsula.

***Species composition***

*Typical species*: The *Tiwi Islands and Cape York* bird community is typified by the prevalence of White-throated Honeyeater, Rufous Whistler, White-bellied Cuckoo-shrike, Leaden Flycatcher, and Little Friarbird. It is best distinguished from other communities in the region by the high reporting rate of White-throated Honeyeater, White-bellied Cuckoo Shrike, Rufous Whistler, Little Friarbird and Leaden Flycatcher, typical of tall open forest vegetation. Also important in distinguishing the community is the virtual absence of Little Shrike-thrush, Magpie-lark, Noisy Friarbird, Varied Triller, White-gaped Honeyeater, Willie Wagtail, and Yellow-spotted Honeyeater.

*Guild structure*: Typical of the Northern Australia region, granivores are strongly represented (e.g. Peaceful Dove, Bar-shouldered Dove) in the *Tiwi Islands and Cape York* bird community. It is a distinctly open-forest community with a dominance of large predatory species (e.g. Laughing Kookaburra (Cape York only), Blue-winged Kookaburra, Pied Butcherbird, Black-backed Butcherbird (Cape York only), and flycatchers (e.g. Leaden Flycatcher, Lemon-bellied Flycatcher, Northern Fantail) along with a mix of nectarivores and insectivores.

*Temporal dynamics*: The species that distinguish this community are typically resident, with a standard mix of honeyeaters and granivores that will track resources within the region depending on season, flowering and land management.

*Geographical variation*: The main variation in this community is between its occurrence on the Tiwi Islands and that on western Cape York Peninsula, resulting in species substitutions like Red-collared/Rainbow Lorikeet and Northern/Pale-headed Rosella, and some species like Laughing Kookaburra, Black-backed Butcherbird, Yellow Honeyeater and Noisy Friarbird occurring only in the east.

***Vegetation/habitat associations***

The vast majority of the *Tiwi Islands and Cape York* bird community occurs in the Eucalypt open forests Major Vegetation Group (MVG).

### **Region C: Arid zone**

The Arid Zone region consists of six bird communities *Arid Open Grassland with Low Shrubs, Arid Open Grassland with Sparse Trees, Arid Shrubby Woodland, Arid Woodland, General Arid Zone,* and *Inland Nomad*. Acacia and Chenopod shrublands and hummock grasslands Major Vegetation Groups (MVGs) dominate this region, with forest/woodland MVGs more common in *Arid Open Grassland with Sparse Trees, Arid Woodland, Inland Nomad* and, to a lesser extent*, Arid Shrubby Woodland* bird communities. Community composition can be spatially and temporally dynamic across the arid zone, particularly in relation to rainfall (Burbidge & Fuller 2007; Tischler et al. 2013) and presence of large trees on watercourses can have over-riding effects on species occurrence (Pavey & Nano 2009; Smith 2015). Where water courses are heavily treed, the bird community is likely to be classified into Region D’s inland Treed Water Source community.

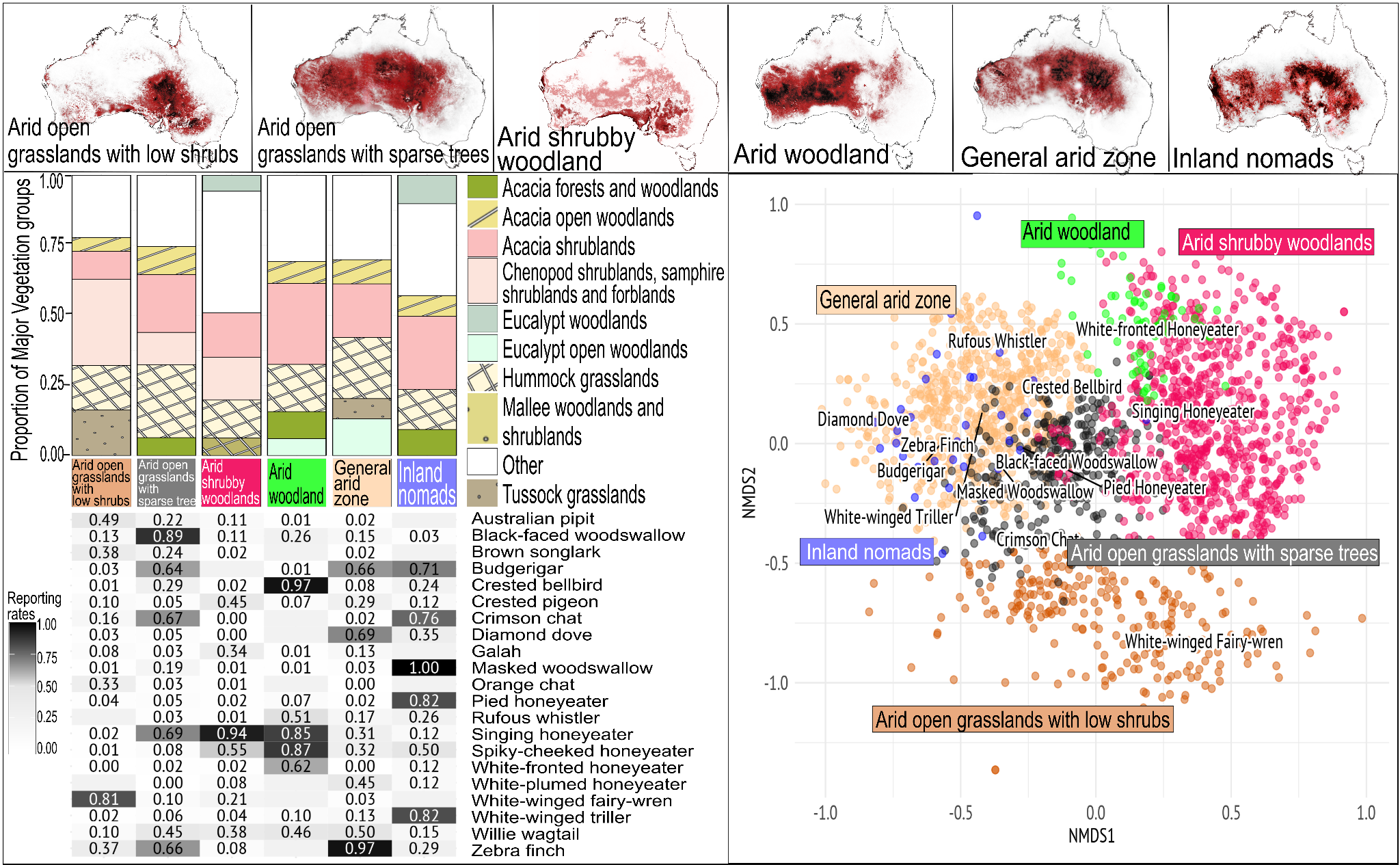

**Figure S1.5** Top across the figure, the distribution of communities, darker shading represents higher likelihood of occurrence. Top left, is the comparison of proportion of Major Vegetation Groups (MVG), the bottom left is the Comparison of Reporting Rates (RR) of the top 5 most commonly occurring species from each community, across all communities in the region. Bottom right shows a non-metric multidimensional scaling (NMDS) plot depicting the different important species distinguishing communities.

#### **Arid Open Grassland with Low Shrubs**

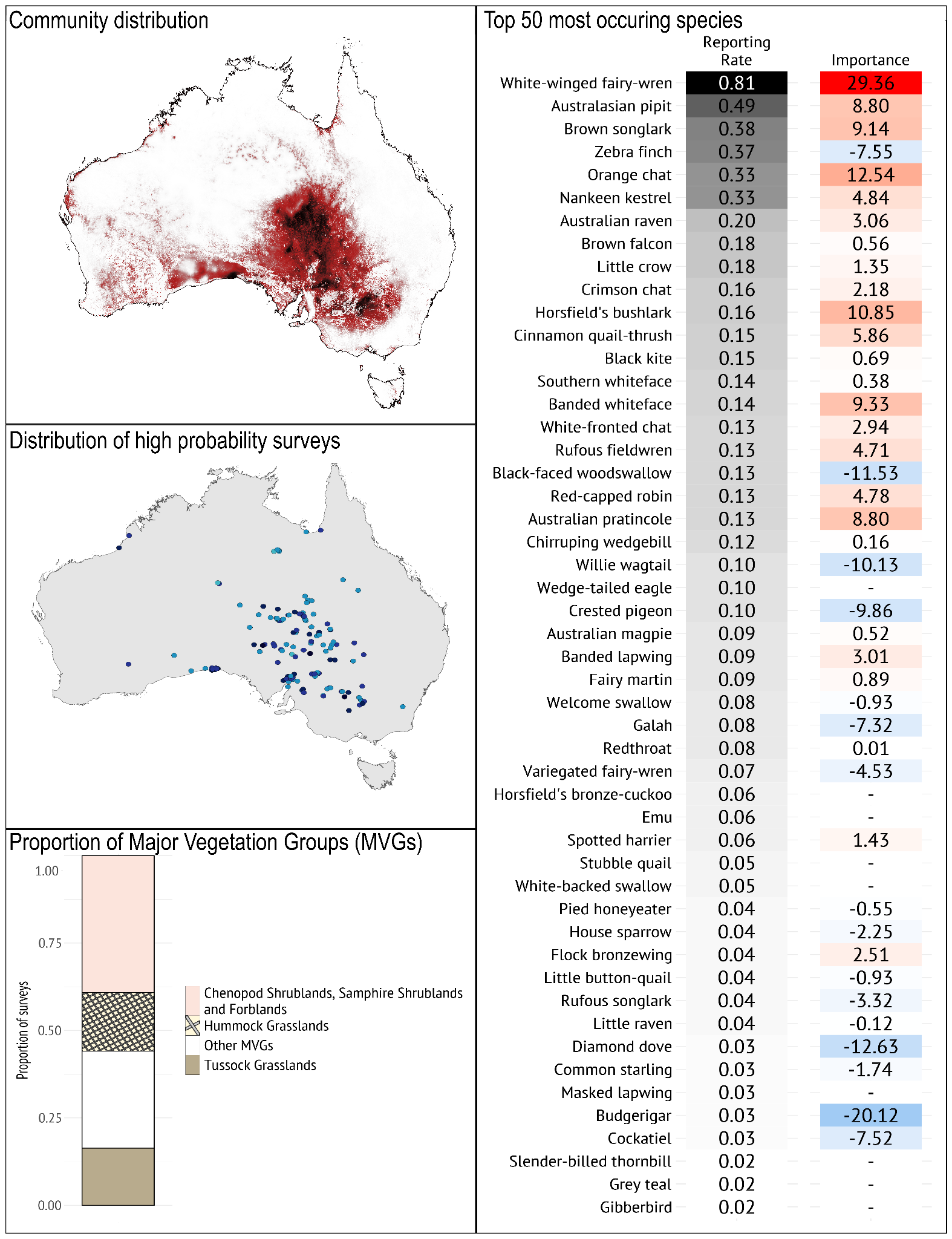

***Geographical occurrence***

The *Arid Open Grassland with Low Shrubs* bird community primarily occurs across arid southern and east-central Australia in the Central Lowlands. The areas where this community is most likely to occur are around the Nullabor Wilderness protected area on the Nullarbor Plain (SA), the gibber plains of Sturts Stony Desert, and across the Simpson and Strzelecki Deserts. It includes the species-poor assemblages in the Simpson–Strzelecki Dunefields bioregion, both in the sandridge deserts themselves and the drier, outer floodplains of the LEB rivers (Reid 1990), as well as the treeless steppes of the Nullarbor Plain (Brooker et al. 1979; McKenzie & Robinson 1987), but also extends to the drier western parts of the Murray-Darlin Basin.

***Species composition***

*Typical species*: The *Arid Open Grassland with Low Shrubs* bird community is typified by the prevalence of White-winged Fairy-wren, Australasian Pipit, Brown Songlark, Zebra Finch, and Orange Chat. It is best distinguished from other arid zone communities by the presence of White-winged Fairy-wren, Orange Chat, Horsfield’s Bushlark and Banded Whiteface. Tall shrubland- and woodland-dependent species are largely absent from this community.

*Guild structure*: This community is dominated by ground feeders: insectivores (e.g. Australasian Pipit, Brown Songlark, Rufous Fieldwren); raptors (e.g. Brown Falcon, Nankeen Kestrel, Black Kite); omnivores (corvids, wedgebills, whitefaces, Horsfield’s Bushlark, Crimson Chat, Cinnamon Quail-thrush); and the generalist arid-zone graminivore, Zebra Finch. Nectar-feeding honeyeaters are scarce except in good seasons.

*Temporal dynamics*: Given the paucity of larger shrubs and trees in these landscapes, bird communities are depauperate in truly sedentary bird species, and so community composition appears especially dynamic depending on seasonal conditions, i.e. the influx of nomadic species after large rainfall and flooding events can outnumber the few residents (Reid 1990). Sparse sedentary Red-capped Robin and Willie Wagtail populations are boosted in autumn-winter by migrants from southern Australia.

*Geographical variation*: The Cinnamon Quail-thrush in the Lake Eyre Basin is replaced by the Nullarbor Quailthrush on the Nullarbor Plain, while the Chiming Wedgebill replaces the Chirruping Wedgebill in western parts of the Simpson Desert and the stony deserts west of lakes Eyre and Torrens. The quail-thrushes, wedgebills, Inland Dotterel, Gibberbird and Banded Whiteface are confined to the arid portions of the mapped community extent.

***Vegetation/habitat associations***

The *Arid Open Grassland with Low Shrubs* bird community is spread relatively evenly across chenopod (including samphire) low shrublands and forblands; hummock grasslands (often dominated by Sandhill Canegrass *Zygochloa paradoxa*); and tussock grasslands Major Vegetation Groups (MVGs). Three plant families dominate numerically the flora of these MVGs, namely Asteraceae, Chenopodiaceae and Poaceae, and annual and short-lived members of the Brassicacea and Fabaceae are also prominent (Mollenmans et al. 1984).

#### **Arid Open Grassland with Sparse Trees/Tall Shrubs**

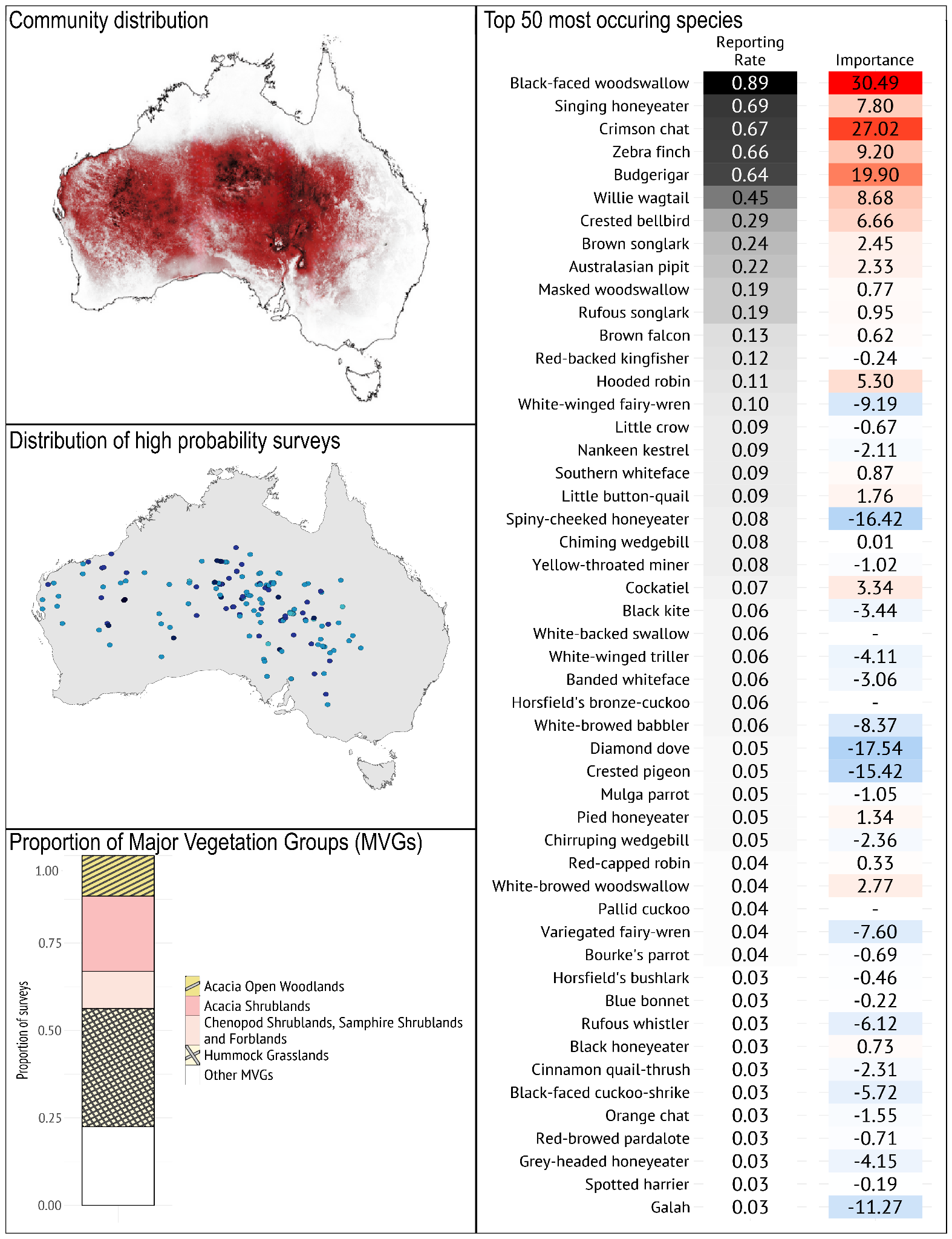

***Geographical occurrence***

The *Arid Open Grassland with Sparse Trees* bird community is widespread across arid Australia in a broad band, stretching from the Pilbara region to Jilbadji Nature Reserve in Western Australia and across central Australia to Dutton River (QLD) and Broken Hill (NSW). Its distribution is notably similar to that of the General Arid Zone bird community, though it covers a wider area, including a broader southerly extent. The areas with the highest probability of this community occurring are areas away from watering points and natural sources of fresh water, i.e. in true desert and salt lake regions, including Tjoritja / West MacDonnell National Park in the Northern Territory (and surrounding ranges), around Lakes Frome, Blanche, and Eyre in South Australia, and Lake Disappointment, Dora and other salt lakes spread across Western Australia.

***Species composition***

*Typical species*: The *Arid Open Grassland with Sparse Trees* bird community is typified by the prevalence of Black-faced Woodswallow, Singing Honeyeater, Crimson Chat, Zebra Finch, and Budgerigar. It is best distinguished from the other arid zone communities by the high reporting rate of Black-faced Woodswallow and Crimson Chat (also prevalent in the *Inland Nomad* bird community), and the relatively low reporting rate of larger water-dependent species (Crested Pigeon and Galah), and species preferring denser, shrubbier vegetation (Purple-backed Fairy-wren, Spiny-cheeked Honeyeater, Rufous Whistler and White-browed Babbler).

*Guild structure*: The majority of species in the *Arid Open Grassland with Sparse Trees* bird community prefer open habitats including ground-foraging insectivores (e.g. Crimson Chat, Willie Wagtail), granivores (e.g. Zebra Finch, Little Button-quail), and predators (e.g. Red-backed Kingfisher, Nankeen Kestrel). However, some tree and tall shrub using/dependent species occur as well (e.g. Singing Honeyeater, Crested Bellbird, and Hooded Robin) but most of these are also ground foragers.

*Temporal dynamics*: Within the extensive hummock grasslands Major Vegetation Group, this bird community is typical of early, post-fire successional states, when bunch grasses and herbs predominate (Reid et al. 1993). There are relatively few migratory species although some of the common species of this community are irruptive or nomadic, with most temporal dynamics likely driven by variation in rainfall among years, as for most communities in this region.

*Geographical variation*: Despite the extensive distribution of this community, most of its typical species occur throughout. Exceptions include the Chiming Wedgebill which is replaced by Chirruping Wedgebill in the east.

***Vegetation/habitat associations***

The *Arid Open Grassland with Sparse Trees* bird community occurs primarily in hummock grasslands, and Acacia shrublands Major Vegetation Groups (MVGs), especially in early successional states.

#### **Arid Western Woodland/Tall Shrubland**

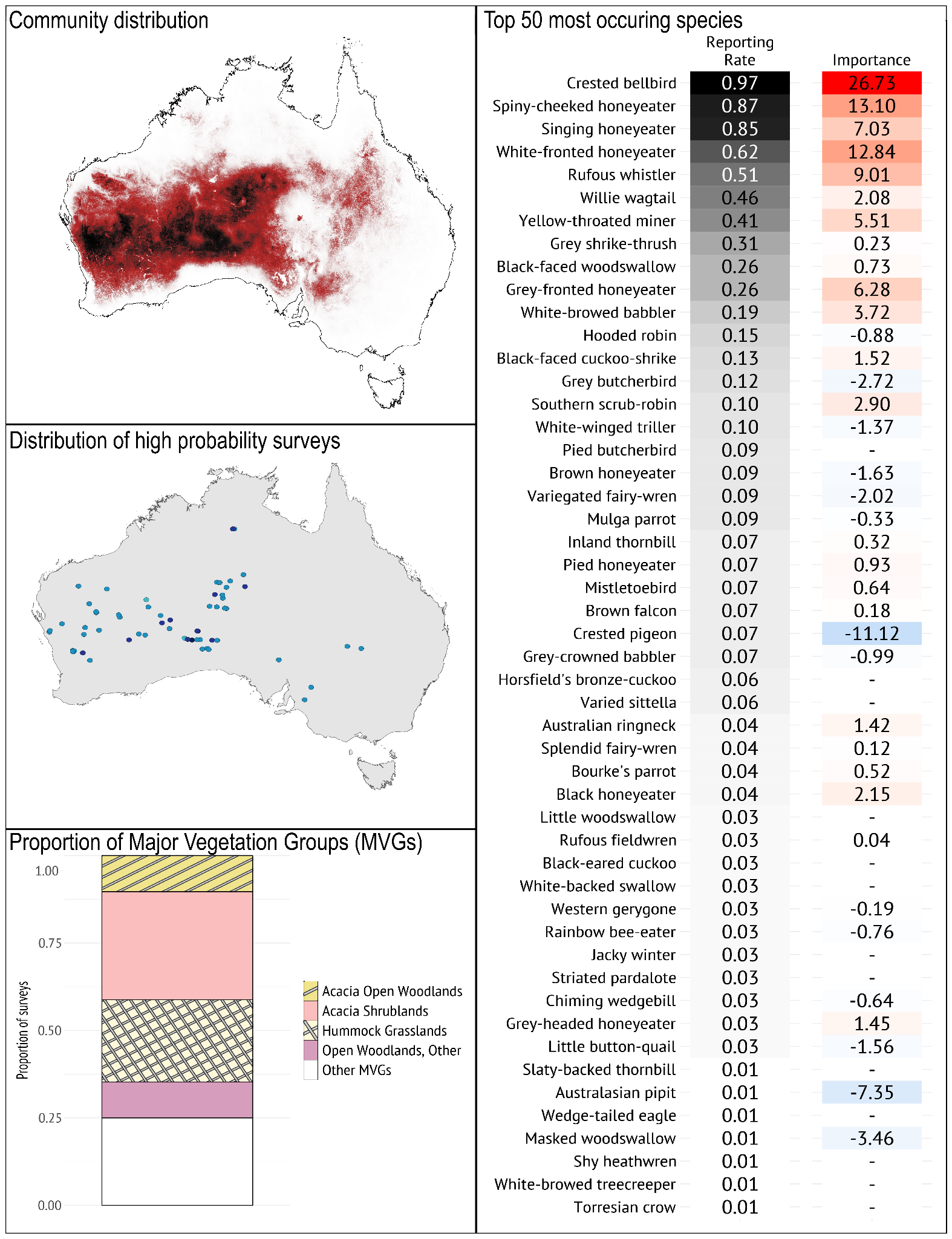

***Geographical occurrence***

The *Arid Woodland* bird community occurs mainly in the central and western parts of the arid zone of Australia from Mount Sheila and Bridgetown in Western Australia to the edge of the Simpson and Strezlecki deserts in the east, especially in areas with approximately 200-300 mm annual rainfall.

***Species composition***

*Typical species*: The *Arid Woodland* bird community is typified by the prevalence of Crested Bellbird, Spiny-cheeked Honeyeater, Singing Honeyeater, White-fronted Honeyeater, and Rufous Whistler. The community is best distinguished from others in the Arid Zone region by the greater probability of occurrence of Crested Bellbird, Rufous Whistler and White-fronted Honeyeater, and by the low probability of occurrence for Budgerigar, Crested Pigeon, and Zebra Finch.

*Guild structure*: The community primarily comprises medium sized insectivores (e.g. Crested Bellbird, Rufous Whistler, Willie Wagtail) and nectarivores (e.g. Spiny-cheeked, Singing, and White-fronted Honeyeaters). The most prevalent species are strongly associated with trees and shrubs, which is fitting given its vegetation associations.

*Temporal dynamics*: This community includes some seasonal migrants like the White-winged Triller and Horsfield’s bronze-cuckoo and Black-eared Cuckoo. These species all migrate to southern Australia in summer (August to February) to breed, and overwinter in the inland.

*Geographical variation*: Southern Scrub-robin, Rufous Fieldwren and Shy Heathwren only occur in the southern examples of this community, while Grey-headed Honeyeater occurs more often in the north.

***Vegetation/habitat associations***

The *Arid Woodland* bird community occurs primarily in Acacia shrublands, and hummock grasslands Major Vegetation Groups (MVGs), but its presence in the mapped hummock grasslands is largely confined to areas with a moderate to high cover of overtopping shrub and low tree canopies including mallee and mulga. It is associated with vegetation groups that have few or no large, hollow-bearing Eucalypts, and instead are dominated by Acacia trees (e.g. Gidyea) or shrubs (e.g. Mulga) that provide habitat for birds and insects that they forage from leaves, flowers and stems. These vegetation types are also typified by a low perennial shrubby understory dominated by species of chenopods and *Eremophila* that provide nesting and foraging habitat for insectivores and nectarivores (Tulloch et al. 2023).

#### **General Arid Zone**

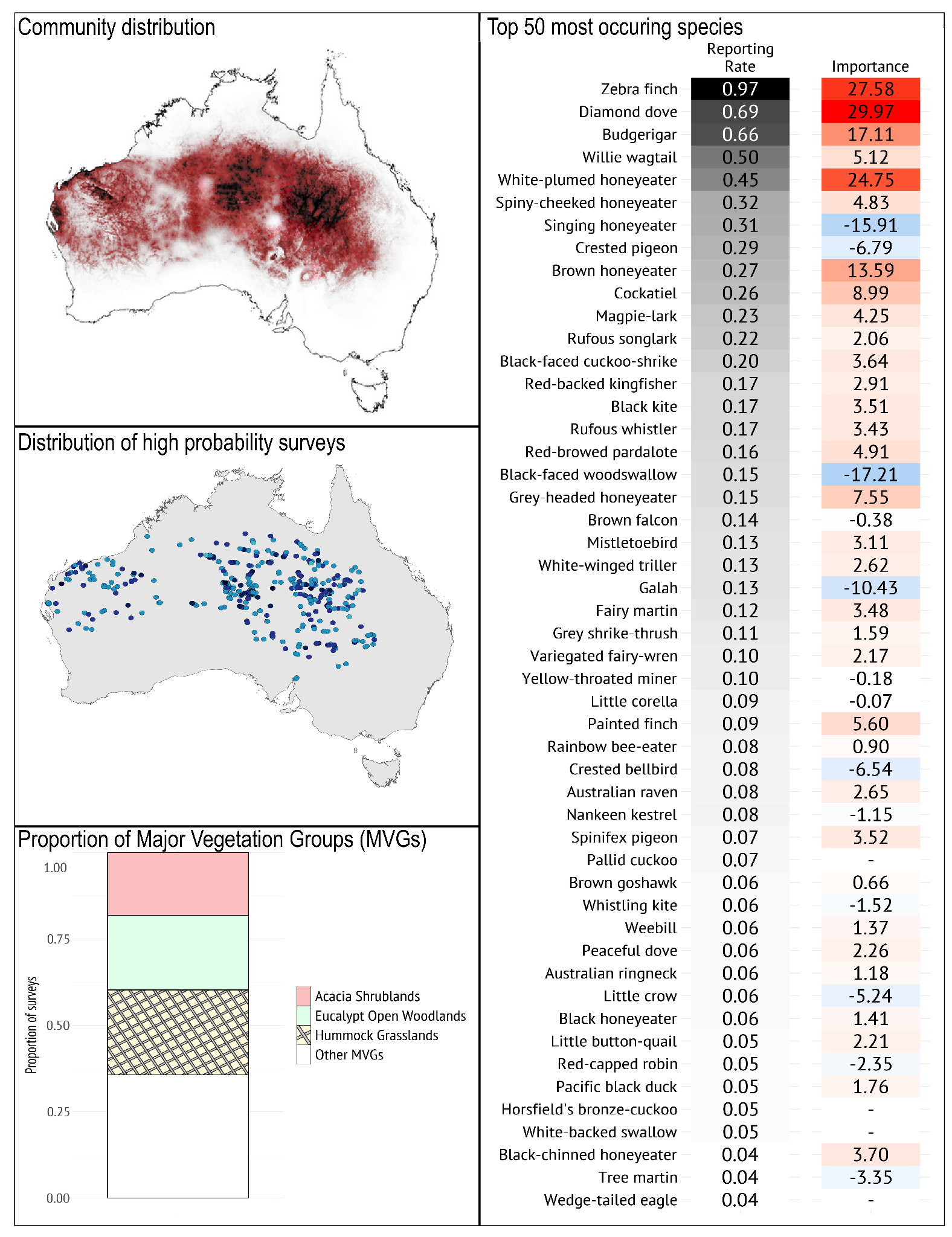

***Geographical occurrence***

The distribution of the *General Arid Zone* bird community is generally similar to that of the Arid Open Grassland with Sparse Trees bird community, but a little more restricted and somewhat further north. It ranges in a band from Eighty Mile Beach (WA) and Kalbarri (WA) in the west, to Gilberton (QLD) and Bourke (NSW) in the east. The areas with the highest probability of this community occurring in areas with higher water availability including around Tjoritja / West MacDonnell National Park in the Northern Territory (and surrounding ranges), in the north-eastern Lake Eyre Basin of south-west Queensland, and around Mount Sheila (WA) and Kennedy Range (WA). It frequently occurs in hilly arid areas.

***Species composition***

*Typical species*: The *General Arid Zone* bird community is typified by the prevalence of Zebra Finch, Diamond Dove, Budgerigar, Willie Wagtail, and White-plumed Honeyeater. It is best distinguished from other communities in the Arid Zone by the prevalence of Zebra Finch and White-plumed Honeyeater, and the low occurrence of Crimson Chat and Crested Bellbird. Although recorded at low frequencies, the arid-zone, rocky range specialists such as Spinifex Pigeon, Grey-headed Honeyeater and Painted Finch are a distinctive component of this broad community.

*Guild structure*: The community is dominated by small ground-foraging granivores (e.g. Zebra Finch, Diamond Dove, Budgerigar), with a substantial representation of nectarivores (e.g. White-plumed, Spiny-cheeked, Singing, and Brown Honeyeaters) and relatively few insectivores relative to other bird communities.

*Temporal dynamics*: With a high proportion of nomadic species in this community, temporal dynamics are likely linked to availability of resources across multi-season time spans. Many of the granivores breed opportunistically (e.g. Budgerigar, Painted Finch, Zebra Finch) and can be eruptive breeders when conditions are favourable (Jordan et al. 2017).

*Geographical variation*: Most species in this community occur throughout its extent, but the Spotted Bowerbird found in Queensland portions of the Lake Eyre Basin is replaced by the Western Bowerbird in central and Western Australia, while different subspecies of Spinifex Pigeon occur from east to west.

***Vegetation/habitat associations***

The *General Arid Zone* bird community occurs primarily in (Spinifex-dominated) hummock grasslands, Eucalypt open woodlands, and Acacia shrublands Major Vegetation Groups (MVGs). Understory in these vegetation types is likely to have a high prevalence of perennial and annual grasses that provide seed for foraging. Dominant trees are likely to provide seasonal flowering resources, and understory probably lacks a diversity of shrubs and flowering species that would provide habitat for insects and insect-foraging birds.

#### **Inland Nomads**

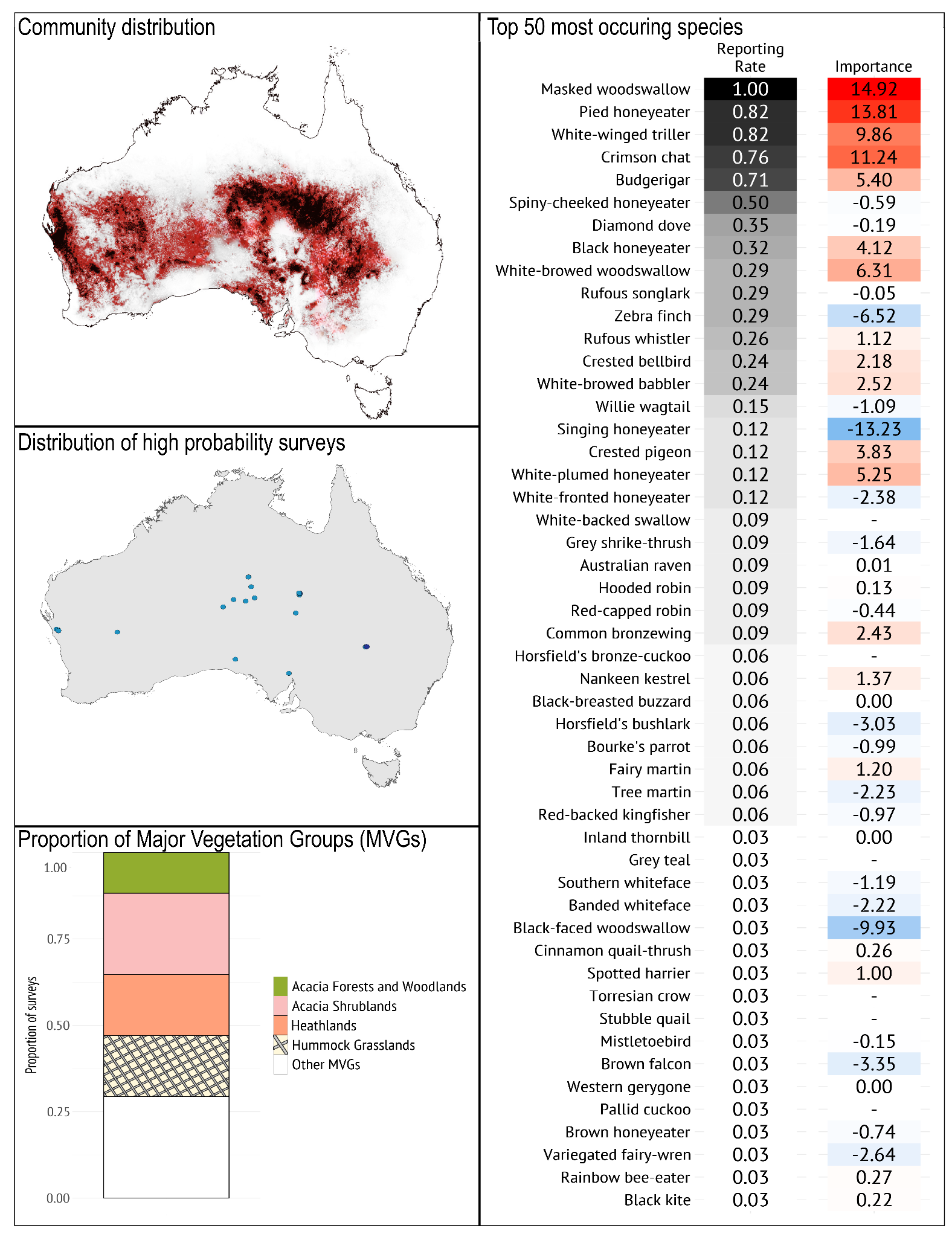

***Geographical occurrence***

The *Inland Nomads* bird community has spatially - and likely, temporally - patchy occurrence across arid regions of western, central and southern Australia. It ranges from Exmouth to Williams in Western Australia, skirting the top of the Nullarbor Plain and around the Simpson Desert to reach as far east as Bourke (NSW).

***Species composition***

*Typical species*: The *Inland Nomads* bird community is almost always associated with Masked (and often White-browed) Woodswallow, and includes several nomadic and/or irruptive species such as Pied Honeyeater, White-winged Triller, Crimson Chat, and Budgerigar. It is best distinguished from other communities in the Arid Zone region by the high prevalence of Masked Woodswallow, Pied Honeyeater, and White-winged Triller

*Guild structure*: This community includes many small- and medium-sized nectarivores, as well as small granivores and aerial and ground-foraging insectivores.

*Temporal dynamics*: This is a highly variable community, and its distribution is likely to change substantially by season and also in response to resource pulses caused by rainfall events.The most common species in this community are migratory (e.g. Masked Woodswallow, White-winged Triller, White-browed Woodswallow, Rufous Songlark) or nomadic and irruptive Budgerigar, Black Honeyeater, Crimson Chat, Diamond Dove, Pied Honeyeater).

*Geographical variation*: The wide-ranging nomads of inland Australia are characterised by their lack of geographical variation, although the complex, recent evolutionary divergence of White-browed and Masked Woodswallows (Joseph 2017) has its origins in west vs east centres of occurrence.

***Vegetation/habitat associations***

The *Inland Nomad* bird community occurs primarily in Acacia shrublands, heathlands, hummock grasslands Major Vegetation Groups (MVGs).

### **Region D: Inland eastern woodland**

This region consists of three bird communities, *Inland Treed Water Source, South-eastern Open Box-Gum Woodland*, and *South-eastern Woodland Inland Slopes*. These communities clustered most closely to many of the community types that were deemed anthropogenic or ‘degraded’. They share species that often occur in open modified environments, such as Willie Wagtail, Australian Magpie and Galah. The geographic location and the types of woodlands/forests in which these communities occur differ substantially and can be considered in two parts.

The *Inland Treed Water Source* community occurs in dry environments extending across much of arid and semi-arid Australia, but within this broad area the community is restricted mainly to open woodlands associated with creeks, drainage lines, flood plains and water points (both natural and anthropogenic). Suitable habitat often is limited in size, such as linear strips of River Red Gum woodland along creeklines, with adjacent vegetation being structurally distinct (e.g. chenopod shrublands). The woodland habitat supports characteristic species that forage in the tree canopy (e.g. White-plumed Honeyeater, Black-faced Cuckoo-Shrike, Yellow-throated Miner), nest in tree hollows (Galah, Australian Ringneck) or favour elevated perch sites (Pied Butcherbird, Whistling Kite, Black Kite). The core elements of this community are relatively stable throughout the year. Drought is a primary environmental disturbance factor, and the woodlands and shrublands that support this community may serve as a refuge for a wider range of species during severe drought.

The *South-eastern Woodland Inland Slopes* community and the *South-eastern Open Box-Gum Woodland* community both occur in temperate south-eastern Australia. The *South-eastern Woodland Inland Slopes* community occurs mainly on the inland slopes of the Great Dividing Range, on gentle slopes and hills, often on nutrient-poor soils. This bird community occupies dry forests and woodlands, commonly with a shrubby understorey. The *South-eastern Open Box-Gum* community commonly occurs on the flatter plains on more-fertile soils, including moister woodlands along creeklines and floodplains. These woodlands typically have a more-open, grassy ground layer and lower density of trees; and give way to open woodlands and native grasslands.

*South-eastern Open Box-Gum Woodland* and *South-eastern Inland Slopes Woodland* bird communities share species that are widespread in woodlands and dry forests of south-eastern Australia (e.g. Grey Shrike-thrush, Superb Fairy-wren, Rufous Whistler, Striated Pardalote, White-winged Chough, Black-faced Cuckoo-shrike), but each also has distinctive elements associated with differences in landform, vegetation types and the dominant eucalypt species. These communities change in composition through the year as a result of an influx of spring-summer migrants from the north (e.g. Sacred Kingfisher, White-browed Woodswallow, Pallid Cuckoo), winter migrants from the ranges (e.g. Yellow-faced Honeyeater, White-naped Honeyeater, Flame Robin), and local seasonal variation in nectar-feeders that track eucalypt flowering (e.g. Musk Lorikeet, Red Wattlebird, Noisy Friarbird). The *South-eastern Open Box-Gum Woodland* bird community is especially similar to the *Dry Lowland South-eastern Woodland* bird community but lacks Grey Fantail and has a much lower reporting rate of Rufous Whistler.

Clearing for agriculture has had a marked effect on the distribution and abundance of species in these two communities (Robinson & Trail 1996; Paton et al. 1999; Ford et al. 2001; Radford et al. 2005; Szabo et al. 2011), particularly the *South-Eastern Open Box-Gum Woodland* community which occupies what has become the Sheep-Wheat Belt of south-eastern Australia. These woodlands now occur largely as remnants, large and small, within a vast sea of grazing and cropping lands. Bird communities become depleted as the amount of wooded cover in the landscape declines (Bennett & Ford 1997; Radford et al. 2005; Cunningham et al. 2014), and as remnants become degraded by stock grazing and/or loss of structural complexity (Loyn 1987; Arnold 1988). Loss and fragmentation of woodlands has greatly advantaged the Noisy Miner, a native species but despotic competitor, which has profoundly altered bird communities over vast areas of these woodlands in south-eastern Australia (Loyn 1987; Grey et al. 1998, 1998; Mac Nally et al. 2012; Maron et al. 2013). Many of the ‘declining woodland birds’ in south-eastern Australia (Reid 1999; Ford et al. 2001; Watson 2011; Bennett et al. 2024) are members of these communities. Extended drought also has a detrimental effect on these communities (Mac Nally et al. 2009; Bennett et al. 2014; Selwood et al. 2015a) especially in association with habitat loss (Haslem et al. 2015).

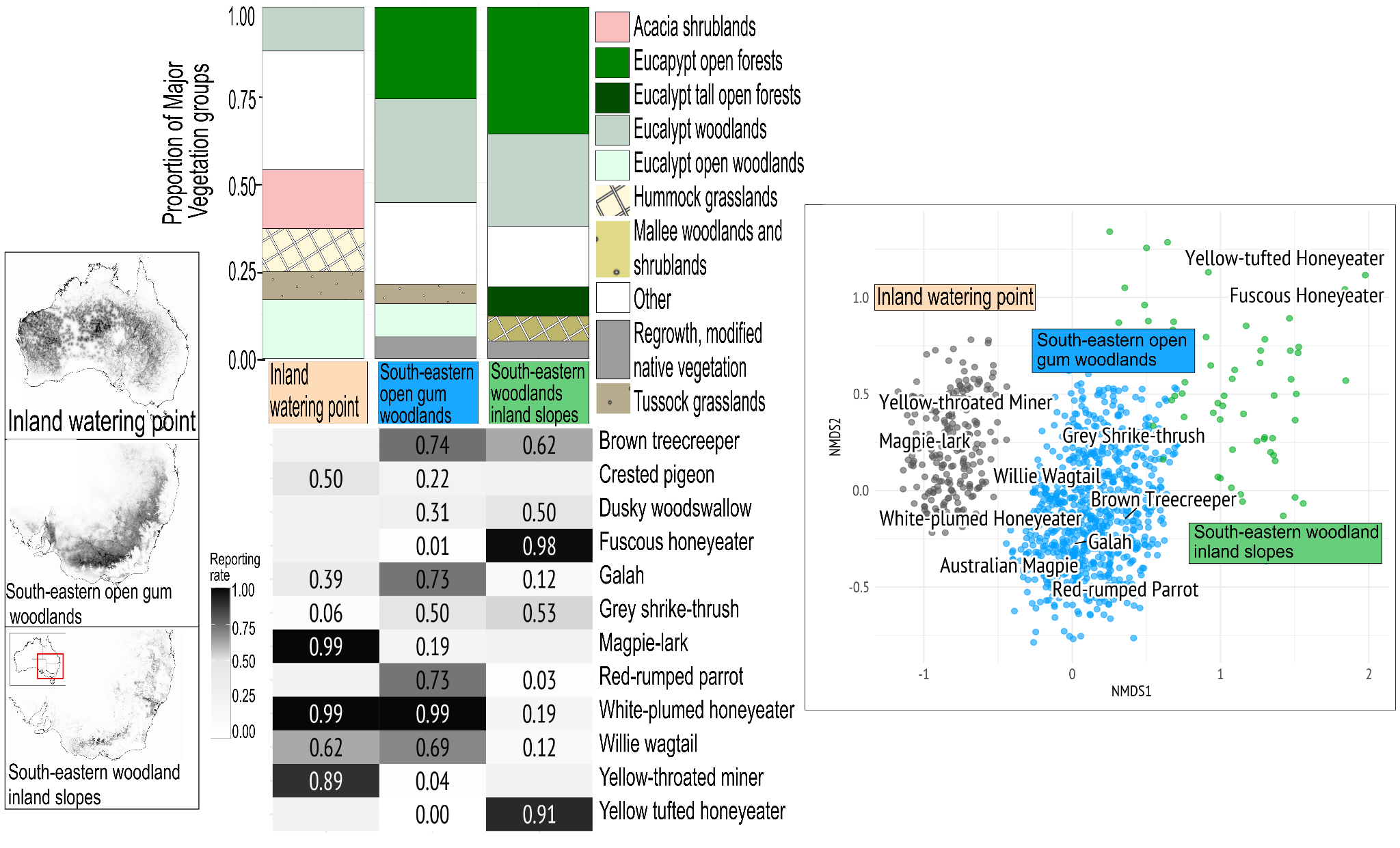

**Figure S1.6** From left to right: far left, maps show distribution of communities, darker shading represents higher likelihood of occurrence. Middle, top is the comparison of proportion of Major Vegetation Groups (MVG) and bottom is the comparison of Reporting Rates (RR) of the top 5 most commonly occurring species from each community. Far right, a non-metric multidimensional scaling (NMDS) plot depicting the different important species distinguishing communities.

#### **Inland Treed Water Source**

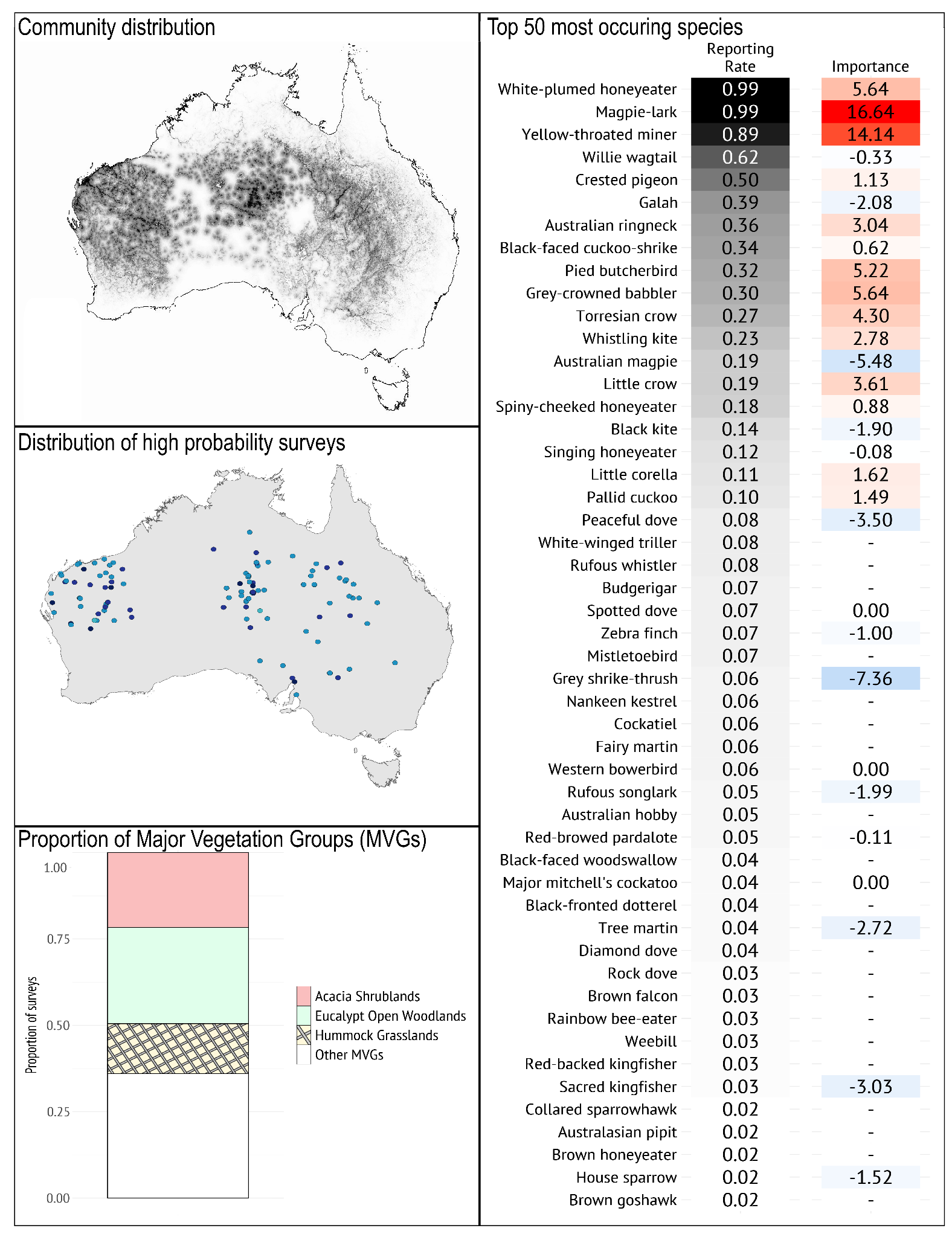

***Geographical occurrence***

The *Inland Treed Water Source* bird community is spread broadly across inland Australia, associated with creeklines, floodplains and permanent or ephemeral water. This community can be found in open arid and semi-arid rangelands where there are tree-lined watercourses or (natural or artificial) watering points.

***Species composition***

*Typical species*: The *Inland Treed Water Source* bird community is typified by the prevalence of the White-plumed Honeyeater, Magpie-lark, Yellow-throated Miner, Willie Wagtail, and Crested Pigeon. It is best distinguished from the other communities in this region by the high prevalence of White-plumed Honeyeater, Magpie Lark, and Yellow-throated Miner.

*Guild structure*: This community has a small number of key species: Magpie-lark and White-plumed Honeyeater occurring in 99% of lists, and Yellow-throated Miner in 89% of lists. The ubiquity of the White-plumed Honeyeater and Yellow-throated Miner means that this community is associated with treed water sources, which often will be adjacent to structurally different vegetation types (e.g. Acacia shrublands, chenopod shrublands, grasslands) with different bird communities that may ‘spill over’ into this one. The *Inland Treed Water Source* community includes a range of insectivores (e.g. Black-faced Cuckoo-Shrike, Red-browed Pardalote, Grey-crowned Babbler, Rufous Whistler, White-winged Triller) and granivores (e.g. Galah, Budgerigar, Crested Pigeon, Peaceful Dove, Australian Ringneck), with lower prevalence of nectarivores indicating ephemeral nectar resources, most likely dependent on flowering seasons of red gums and bloodwoods. Larger predators, such as Whistling Kite, Black Kite, Pied Butcherbird and Little Crow, regularly use trees in this vegetation for perching, but forage widely beyond it.

*Temporal dynamics*: The composition of this community is largely stable, being associated with water, with relatively few migratory species (e.g. White-winged Triller, Pallid Cuckoo) and a few nomadic/irruptive species (e.g. Budgerigar). Species that distinguish this community (Australian Ringneck, Magpie-lark, Pied Butcherbird) often use floodplain habitat as refuge during drought (Selwood et al. 2015a, 2015b).

*Geographical variation*: As this community occurs around disjunct watering points, the composition is likely influenced by the surrounding bird communities. The core list of species that are reliant on the watering points will be supplemented by species from the surrounding communities that also utilise this key resource.

***Vegetation/habitat associations***

The community occurs primarily in Eucalypt woodland and open woodland Major Vegetation Groups (MVGs), associated with drainage lines (e.g. River Red Gums, Coolabahs and Bloodwoods). Other vegetation types, such as Acacia shrublands and Hummock Grasslands may occur adjacent to the watercourses.

#### **South-eastern Open Box-Gum Woodland**

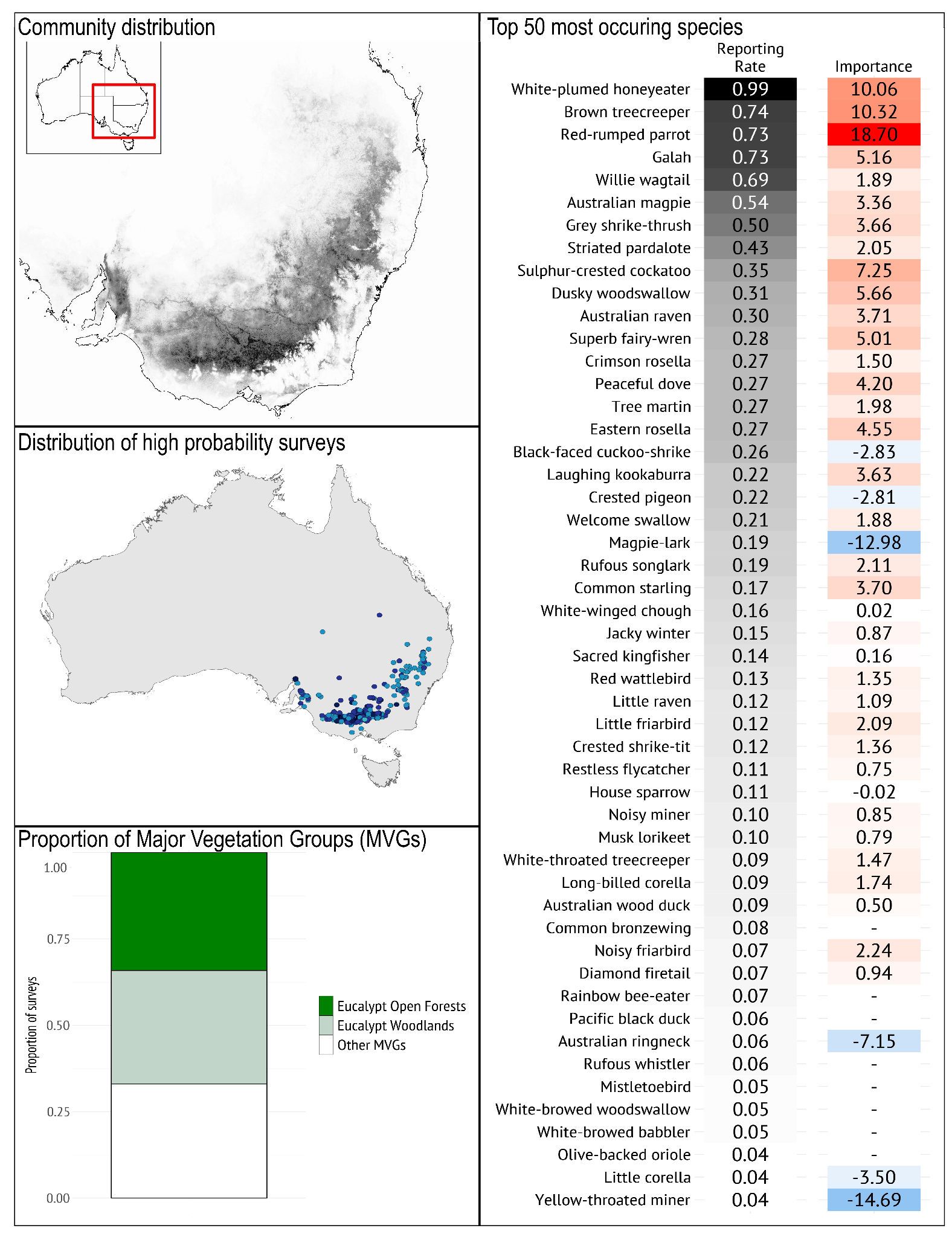

***Geographical occurrence***

The *South-eastern Open Box-Gum Woodland* bird community occurs from the ranges north of Adelaide (SA), through central and northern Victoria, to inland NSW and reaching as high as Brisbane (QLD). It occurs mainly inland of the Great Dividing Range, on plains with reasonably fertile soil, in areas with between 300 and 600 mm annual rainfall; a higher probability of occurrence is associated with higher rainfall areas within that range.

***Species composition***

*Typical species*: The *South-eastern Open Box-Gum Woodland* bird community is typified by the prevalence of White-plumed Honeyeater, Brown Treecreeper, Galah, Red-rumped Parrot, Willie Wagtail, Australian Magpie, Grey Shrike-thrush and Striated Pardalote. It is best distinguished from other communities in this region by the high prevalence of Galah and Red-rumped Parrot, and the absence of Fuscous and Yellow-tufted Honeyeaters.

*Guild structure*: This community occurs in woodland habitats of the plains (e.g. box woodlands) and associated stream systems (e.g. River Red Gum woodlands), typified by numerous hollow-bearing trees. Consequently, a distinctive feature of this community is the diversity of hollow-nesting species including parrots and cockatoos (e.g. Eastern Rosella, Sulphur-crested Cockatoo, Red-rumped Parrot, Galah) and other species (Brown Treecreeper, Sacred Kingfisher, Laughing Kookaburra, Striated Pardalote) (Loyn 1985; Lunt & Bennett 2000). The open ground layer is likely maintained in some areas through grazing and, though it allows a range of ground-foraging species, the most prevalent species are resilient to grazing (e.g. insectivores: Willie Wagtail, Australian Magpie, Restless Flycatcher, Jacky Winter, and granivores: Red-rumped Parrot, Peaceful Dove, Galah, Diamond Firetail) (Antos & Bennett 2005, 2006). This community includes a number of species of conservation concern, listed as ‘declining’ woodland birds (e.g. Diamond Firetail, Brown Treecreeper, Jacky Winter, Dusky Woodswallow) (Reid 1999; Watson 2011; Bennett et al. 2024). A characteristic species of this community, the White-plumed Honeyeater, is a canopy-feeding species that is particularly widespread and abundant in streamside River Red Gum vegetation. Where it is common, it may exclude small insectivores, such as thornbills and whistlers.

*Temporal dynamics*: Spring-summer visitors to this community include Rufous Songlark, Sacred Kingfisher, Rainbow Bee-eater, White-browed Woodswallow, Olive-backed Oriole. The local abundance of nectar-feeders (e.g. Red Wattlebird, Noisy Friarbird, Musk Lorikeet) can vary greatly depending on the flowering patterns of dominant eucalypt species.

*Geographical variation*: There is no substantial geographic variation in species composition throughout the extent of this community.

***Vegetation/habitat associations***

The *South-eastern Open Box-Gum Woodland* bird community occurs primarily in Eucalypt open forests, and Eucalypt woodlands Major Vegetation Groups (MVGs), on relatively fertile soils of plains and lower slopes. This includes forests and woodlands along streams, rivers and floodplains, characterised by River Red Gum; and box-dominated woodlands of the plains (e.g. Grey Box *E. microcarpa*, Yellow Box *E. melliodora*, White Box *E. albens* and Yellow Gum *E. leucoxylon*)*.* The ground layer typically is open and grassy with relatively few shrubs. These latter box-dominated woodlands are now largely cleared for agriculture: small remnants among farmland and along roadsides commonly are dominated by the Noisy Miner, which aggressively excludes small insectivores.

#### **South-eastern Woodland Inland Slopes**

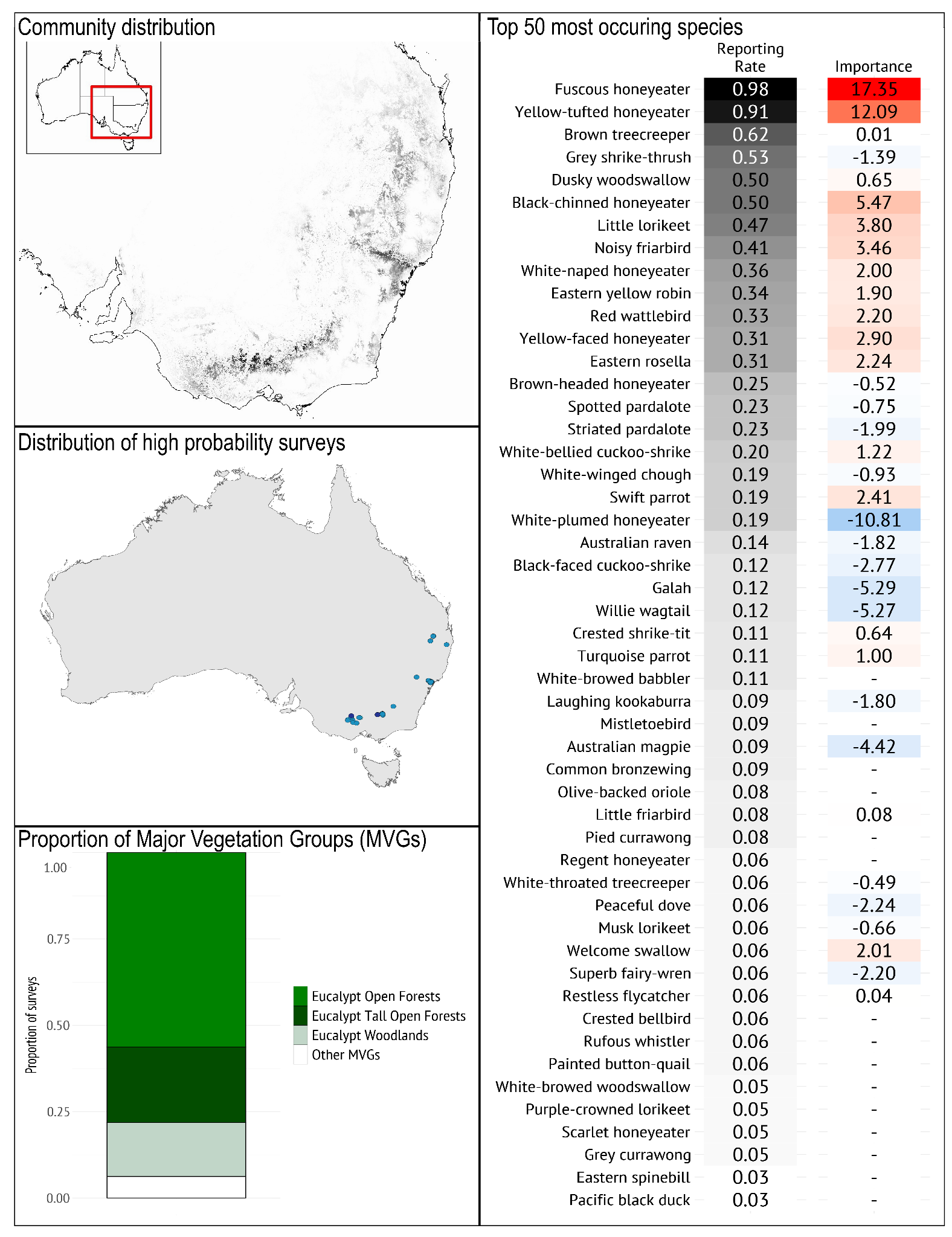

***Geographical occurrence***

The *South-eastern Woodland Inland Slopes* bird community is sparsely distributed along the Great Dividing Range and inland slopes in Victoria and New South Wales, with hotspots around Bendigo and Heathcote (Vic), Wangaratta (Vic) and Wodonga (Vic)/Albury (NSW), and the Hunter Valley (NSW).

***Species composition***

*Typical species*: The *South-eastern Woodland Inland Slopes* bird community is typified by the prevalence of the Fuscous Honeyeater, Yellow-tufted Honeyeater, Brown Treecreeper, Grey Shrike-thrush, Dusky Woodswallow, Brown-headed Honeyeater and Black-chinned Honeyeater. It is best distinguished from other bird communities in the region by the high prevalence of Fuscous Honeyeater and Yellow-tufted Honeyeater, and the relatively low prevalence of Crested Pigeon, Galah, White-plumed Honeyeater, and Willie Wagtail.

*Guild structure*: A characteristic feature of this bird community is the diversity and abundance of nectar-feeding species (honeyeaters, lorikeets, friarbirds), which have a strong influence on the spatio-temporal dynamics of the community (Mac Nally & McGoldrick 1997; McGoldrick & Mac Nally 1998; Radford & Bennett 2007).

The shrubby ground layer provides habitat for species that use low shrub cover (e.g. White-browed Babbler, Superb Fairy-wren, Speckled Warbler, Crested Bellbird); whereas there tends to be fewer ground-foraging insectivores (such as Buff-rumped Thornbill, Scarlet Robin) than in more open woodlands. Seed-eating birds are scarce in this community (Loyn 1985).

*Temporal dynamics*: Spring-summer migrants include White-browed Woodswallow, Olive-backed Oriole and Sacred Kingfisher, but do not make up a major component of the community. Resident nectarivores (e.g. Red Wattlebird, Musk Lorikeet, Little Lorikeet, Fuscous Honeyeater, Yellow-tufted Honeyeater) move within the forests, depending on flowering patterns, and are particularly abundant in years when winter-flowering eucalypts (e.g. Red Ironbark *E. tricarpa*, Mugga Ironbark *E. sideroxylon*, Yellow Gum *E. leucoxylon*) flower heavily. In such years, other nectarivores may migrate into the system from forests of the Great Dividing Range (e.g. White-naped Honeyeater, Yellow-faced Honeyeater, Eastern Spinebill) or from elsewhere (Noisy Friarbird). In years when these tree species flower sparsely, or not at all, there is a mass movement of nectarivores out of the ecosystem (Mac Nally et al. 2009).

The critically endangered Swift Parrot migrates annually from Tasmania, often occurring in this community over winter months, tracking variable nectar and lerp resources (Mac Nally & Horrocks 2000). The spatiotemporal breeding and foraging patterns of the critically endangered Regent Honeyeater are also influenced by flowering patterns in this community (Franklin et al. 1989; Crates et al. 2017). Periods of severe drought (impacting resource availability) in this community have caused wide scale declines across guilds, which can persist post-drought (Mac Nally et al. 2009, 2020).

*Geographical variation*: There is no substantial geographic variation in species composition throughout the extent of this community

***Vegetation/habitat associations***

The *South-eastern Woodland Inland Slopes* bird community occurs primarily in Eucalypt open forests, and Eucalypt woodlands Major Vegetation Groups (MVGs). These dry forests and woodlands occur on inland slopes and hills, typically on nutrient-poor soils with low water-holding capacity. Termed ‘box and ironbark’ forests, these vegetation types include a range of eucalypt species that vary in composition depending on topographic position. Ironbarks (e.g. Red Ironbark, Mugga Ironbark) together with species such as Red Stringybark and Red Box, typically occur on dry slopes; whereas Grey Box and Yellow Gum occur on lower slopes. The understorey typically is shrubby, including Acacia spp and a range of sclerophyllous shrubs. Groundstorey (e.g. litter depth and cover, coarse woody debris cover and density) and understorey features (e.g. shrub cover) are important elements affecting species occurrence in these woodlands, providing foraging, shelter and nesting resources (Yen et al. 2011). These vegetation types have been highly impacted by clearing for agriculture, causing a high-degree of habitat fragmentation and vegetation degradation, which have interacted with the pressures of Noisy Miners and climate-drying to substantially impact flowering and resource availability, and hence bird community composition, richness and breeding (Radford & Bennett 2007; Mac Nally et al. 2009; Bennett et al. 2014).

### **Region E: South-western Australia, Mallee, Mulga**

This region consists of seven bird communities, *South-western Forest, South-western Woodland, Mediterranean Mallee-heath, Open Mallee, Shrubby Mallee, Inland Woodland with Eucalypt,* and *Mulga*. *Mediterranean Mallee-heath, Open Mallee,* and *Shrubby Mallee* all occurred frequently in the mallee woodland and shrublands and Eucalypt woodland Major Vegetation Groups (MVGs). The *South-western Forest* and *South-western Woodland* bird communities occurred most in forest and woodland MVGs respectively. Five of the seven communities in this region are distributed across both eastern and western Australia, indicating that the conformity of these bird communities over broad scales outweighs the effect of species restricted to either the east or west.

Eucalyptus woodland associations predominate the bird communities in this region, with mallee vegetation communities prominent across the east and west. However, not all bird communities are associated with Eucalypt woodlands; the widespread *Mulga* bird community is more closely associated with Acacia dominant (predominantly *Acacia aneura*) shrublands and woodlands, and the *South-western Forest* bird community is closely associated with Eucalypt open forests and Eucalypt tall open forests. The banksia woodlands of the Swan Coastal Plain and Kwongan proteaceous heathlands of the south-west biodiversity hotspot are typified by a distinct nectarivore-rich bird assemblage but are not represented among the current typology.

The bird communities *Inland Woodland with Eucalypt* and *Mulga* were very similar in terms of species composition, with the main differences related to a subset of Eucalypt-associated bird species that were rare in the *Mulga* bird community (Australian Ringneck, Grey Fantail, Striated Pardalote and Weebill). Fire is less a driver of bird species composition in mulga woodlands as compared with dynamic rainfall patterns, but where fire does occur, it does impact the structural characteristics of Mulga woodlands, which then influences changes in bird communities (Leavesley et al. 2010).

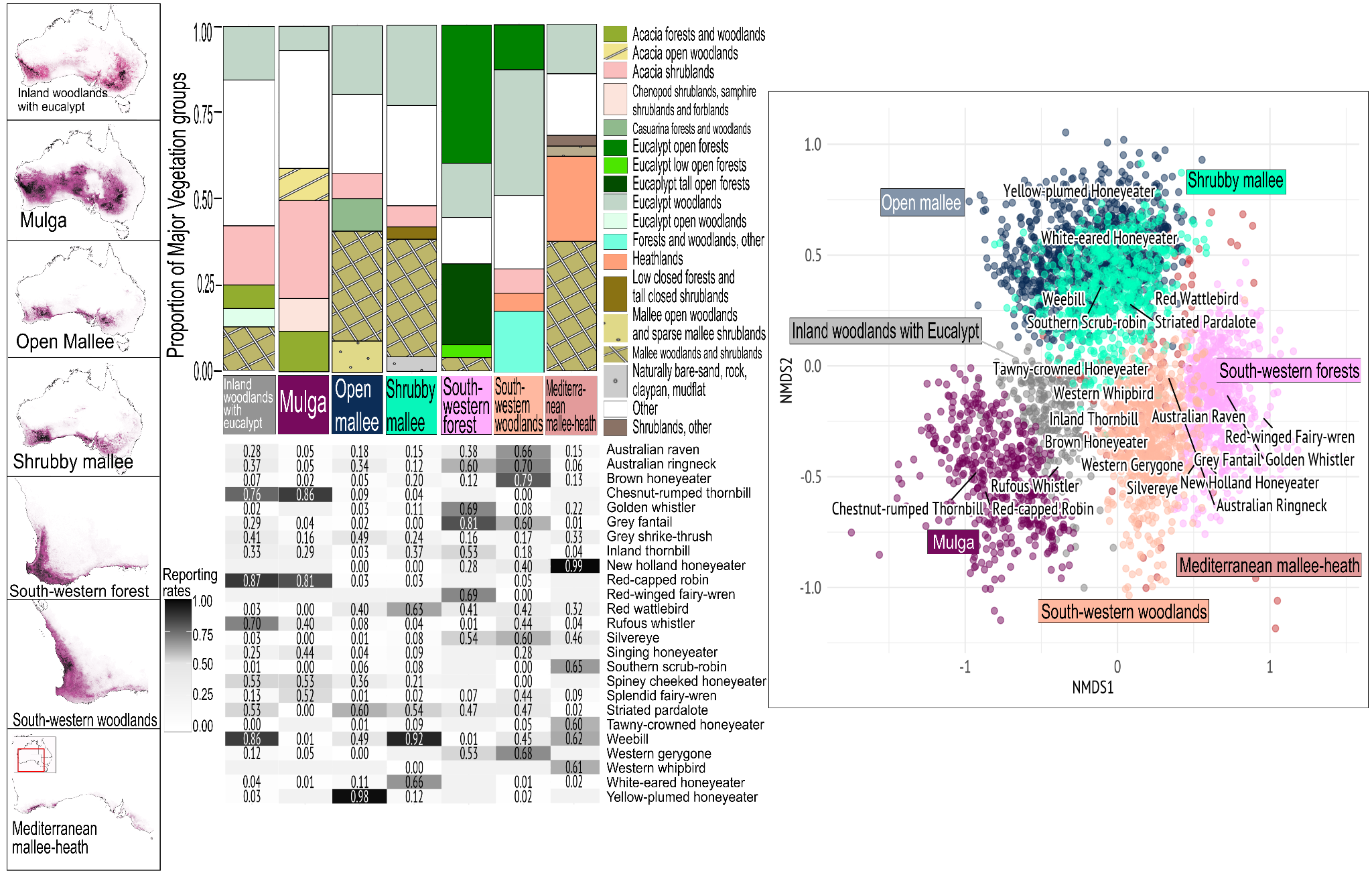

**Figure S1.7** From left to right: far left, maps show distribution of communities, darker shading represents higher likelihood of occurrence. Middle, top is the comparison of proportion of Major Vegetation Groups (MVG) and bottom is the comparison of Reporting Rates (RR) of the top 5 most commonly occurring species from each community. Far right, a non-metric multidimensional scaling (NMDS) plot depicting the different important species distinguishing communities.

**South-western Forest**
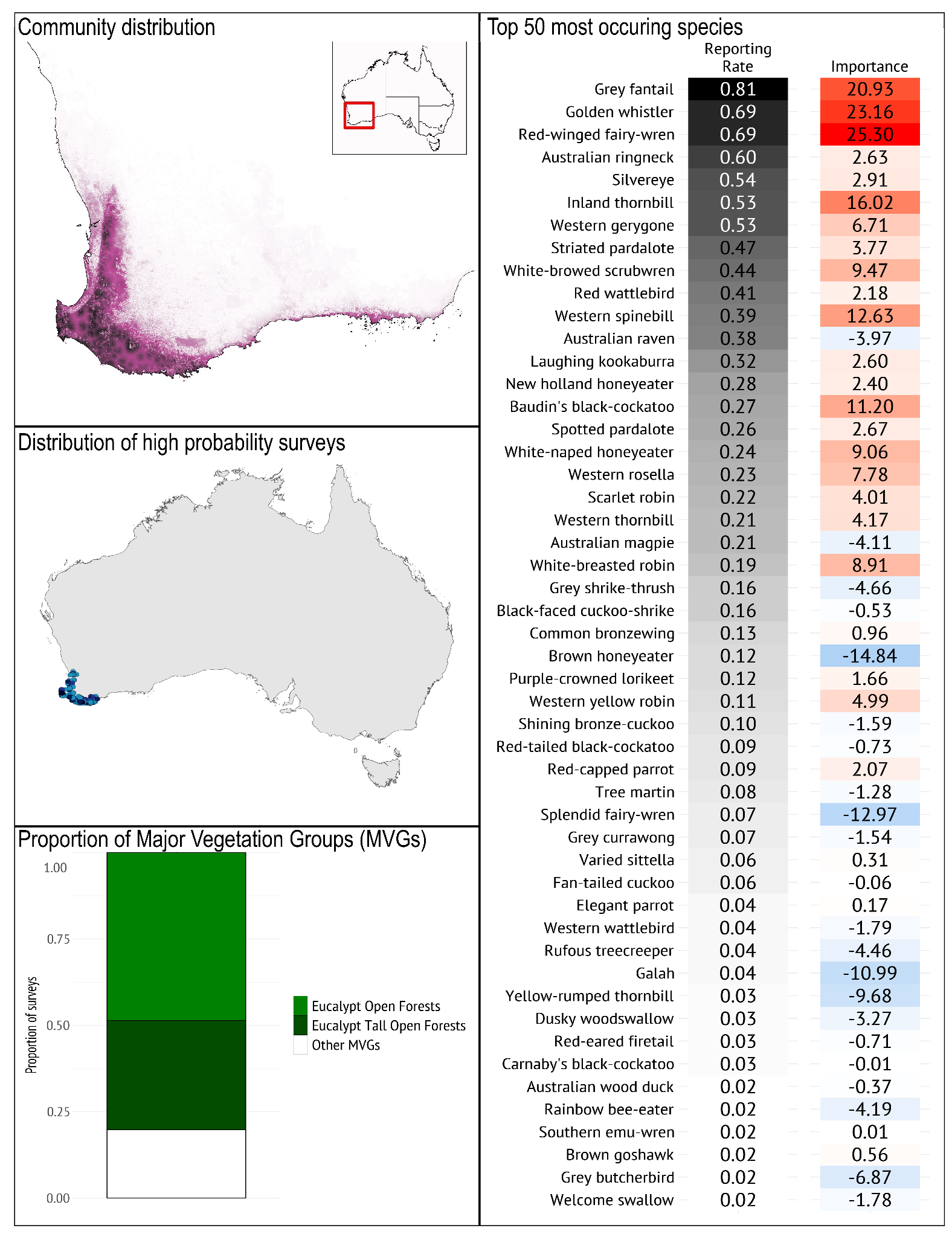

***Geographical occurrence***

The *South-western Forest* bird community occurs in the south-west corner of Western Australia, from Bindoon (north of Perth, WA), running around the western corner of WA with the main occurrence of the community ending at Cape Riche, and pockets of occurrence as far east as around Walpole-Albany and inland at the Porongorups. The community is most likely to be found in wetter areas within the range.

***Species composition***

*Typical species*: The *South-western Forest* bird community is typified by the prevalence of Grey Fantail, (Western) Golden Whistler, Red-winged Fairy-wren, Australian Ringneck, and Silvereye. It is best distinguished from other bird communities in the South-western Australia region by the prevalence of Western Gerygone, Golden Whistler, Baudin’s Black-Cockatoo, and Red-winged Fairy-wren, and the low prevalence of Rufous Whistler.

As the most mesic bird community in the cluster, *South-western Forests* had the highest rates of occurrence for the Grey Fantail and Golden Whistler; two species more commonly encountered in mesic regions than in the arid-zone. In addition, this is the only bird community in which Red-winged Fairy-wren (a mesic-habitat associated bird) is present. Notably, the Red-winged Fairy-wren is absent from the *South-western Woodlands* bird community, which also overlapped with this species’ distribution.

*Guild structure*: The *South-western Forest* bird community is dominated by small- and medium-bodied insectivores that forage in a range of strata from the ground to the canopy (e.g. Grey Fantail, Red-winged Fairy-wren, Golden Whistler), and many parrots and cockatoos (e.g. Australian Ringneck, Baudin’s Black-Cockatoo, Western Rosella, Purple-crowned Lorikeet) with relatively fewer honeyeater species than other communities.

*Temporal dynamics*: The mass flowering of marri, jarrah and karri are associated with a large influx of nectarivores especially Purple-crowned Lorikeets, Western Spinebill, Brown Honeyeater, New Holland Honeyeater and the Western Australian White-naped Honeyeater (now considered its own species: Gilbert’s Honeyeater).

*Geographical variation*: The limited distribution of this community means that its common species all occur throughout its extent.

***Vegetation/habitat associations***

The *South-western Forest* bird community occurs primarily in Eucalypt open forests, and Eucalypt tall open forests Major Vegetation Groups (MVGs). The dominant tree species in these forests include Karri *Eucalyptus diversicolor,* Jarrah *E. marginata* and Marri C*orymbia calophylla*. The understorey is often dense with intermittent occurrences of heathlands and granite-outcrop associated vegetation. The forests in the south-west are typically taller, with a denser understory, than those in the north and east. Karri forests occur only in the south-west of this area, while Jarrah and Marri forests are more widespread. Across the region, the forests have been heavily impacted by logging activities, as well as clearing for agriculture and mining in some areas, particularly in the north (Vasuki et al. 2019).

#### **South-western Woodland**

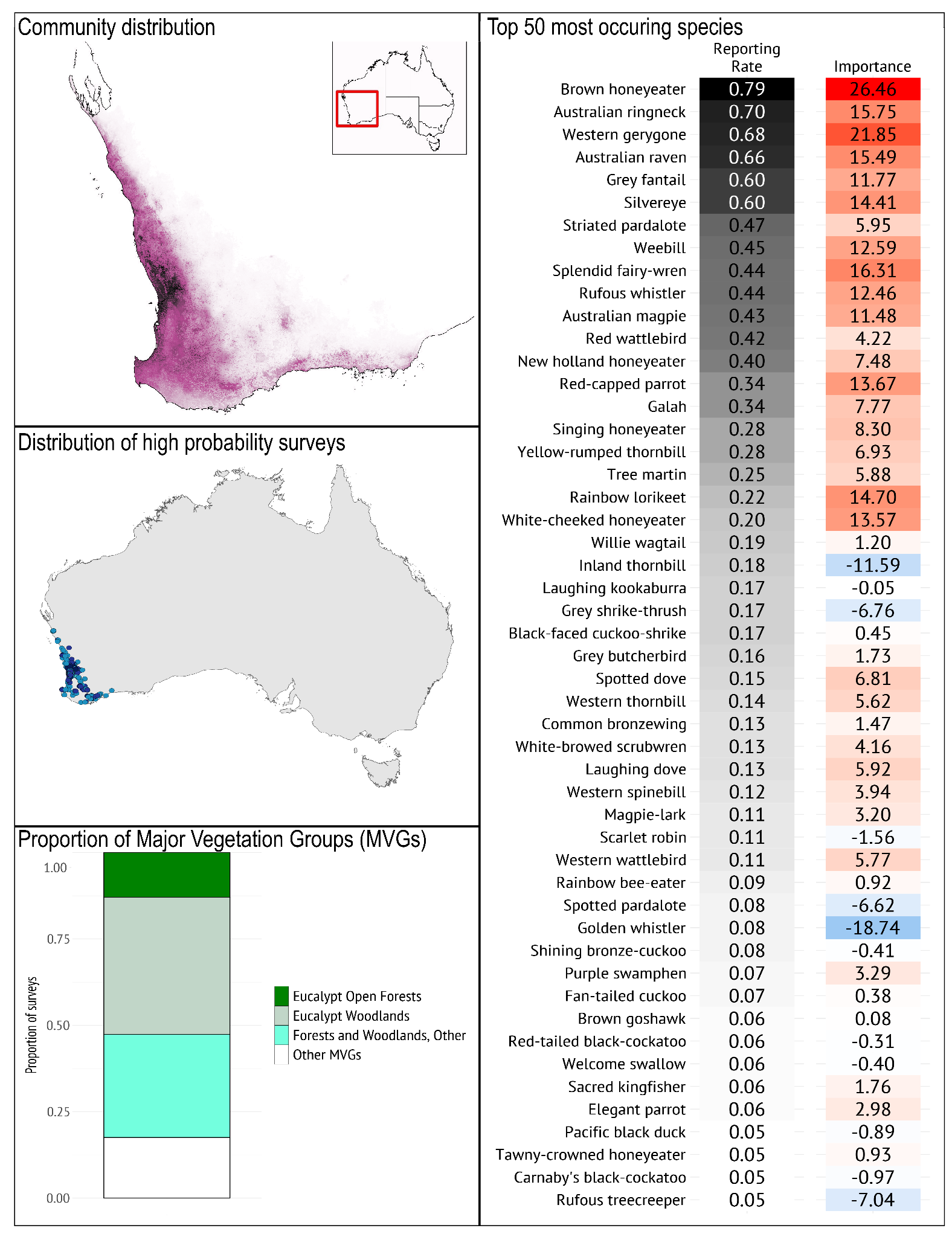

***Geographical occurrence***

The *South-western Woodland* bird community occurs in the south-western corner of Western Australia, extending from near Shark Bay in the north, around the south-west corner and finishing (sparsely) around Cape Arid.

***Species composition***

*Typical species*: The *South-western Woodland* bird community is typified by the prevalence of Brown Honeyeater, Australian Ringneck, Western Gerygone, Australian Raven, and Grey Fantail. It is best distinguished from other bird communities in the region by the high prevalence of Brown Honeyeater. The Splendid Fairy-wren is common in the *South-western Woodland* bird community (in contrast to the Red-winged Fairy-wren in the *South-western Forest* community). In addition, the *South-western Woodland* bird community is distinguished from the *South-western Forest* community by the prevalence of the Weebill, Rufous Whistler, and Singing Honeyeater. Whilst the above differences split these two bird communities, they share similarly high prevalence of Western Gerygone, Silvereye, Grey Fantail and Australian Ringneck.

*Guild structure*: The *South-western Woodland* bird community has few Western Australian endemic species. The species with the highest reporting rate in this community have wide distributions. There is a mix of insectivores, honeyeaters and granivores, with more honeyeaters represented than in the *South-western Forest* bird community.

*Temporal dynamics*: There are some distinct seasonal movements of species endemic to this vegetation association, the most obvious being Carnaby’s Black-Cockatoo which feeds, breeds and roosts throughout this region. It is seen in large flocks in late summer and autumn and relies on banksia woodlands for feeding. The mass flowering of banksia and wandoo in late winter and spring drive an influx of nectarivores particularly the Purple-crowned Lorikeet (in wandoo woodlands) and Western Spinebill, White-cheeked Honeyeater and Tawny-crowned Honeyeater (banksia woodlands).

*Geographical variation*: The banksia woodlands of the Swan Coastal Plain and Kwongan proteaceous heathlands of the south-west biodiversity hotspot are typified by a distinct nectarivore-rich bird assemblage that is not well represented by the species composition of the *South-western Woodland* community.

***Vegetation/habitat associations***

The *South-western Woodland* bird community occurs primarily in the drier and semi-arid Eucalypt woodlands and banksia woodlands and other forests and woodlands Major Vegetation Groups (MVGs). Dominant woodland species include York Gum (*E. loxophleba*) and wandoo (*E. wandoo*). The coastal distribution is dominated by banksia woodlands (*Banksia menziesii*, *B. attenuata* and *B. prionotes*) including the Swan Coastal Plain of Perth which contains extensive banksia woodlands with pockets of Tuart forest and mixed occurrences of jarrah and marri. The northern and eastern parts of this vegetation complex are drier with lower woodlands and differ to the taller woodlands in the higher rainfall zones of the south-west. An iconic species that is almost entirely restricted to this vegetation system is Carnaby’s Black-Cockatoo that breeds and feeds entirely in South-western Woodlands (Saunders 1990).

#### **Mediterranean Mallee-heath**

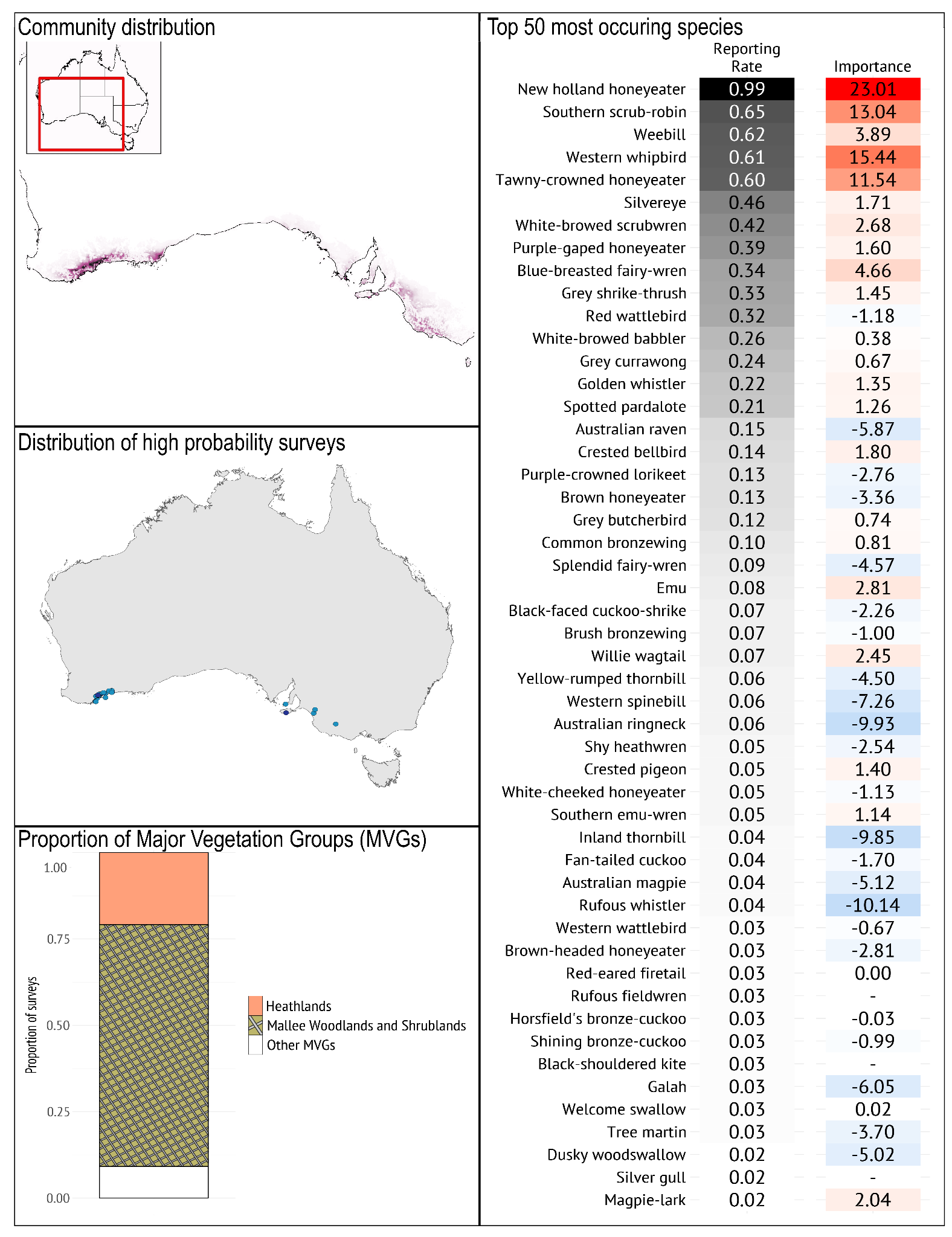

***Geographical occurrence***

The *Mediterranean Mallee-heath* bird community has a very restricted range, occurring from around Albany (WA) in the west to Jerdacuttup, then again in and around Cape Arid National Park (WA), and in SA on Kangaroo Island near Cape Gantheaume and Dudley, and on the mainland in and around Ngarkat Conservation Park, along the eastern edge of the Stirling Ranges, and in the patchy remnant vegetation near the SA/Vic border.

***Species composition***

*Typical species*: The *Mediterranean Mallee-heath* bird community is typified by the prevalence of New Holland Honeyeater, Western Whipbird, Southern Scrub-robin and Tawny-crowned Honeyeater. It is also distinguished from other communities in this region by the low prevalence of Rufous Whistler and Inland Thornbill (associated with drier regions), and the Australian Ringneck (associated with more forested regions).

*Guild structure*: The *Mediterranean Mallee-heath* bird community uniquely has high rates of occurrence for mesic associated shrub-dependent nectarivores such as Tawny-crowned Honeyeater and New Holland Honeyeater. In addition, mallee/heath-associated species such as Southern Scrub-robin and Western Whipbird were common in this community. In general, this community consists of species associated with dense low vegetation and eucalyptus shrublands, but not associated with taller eucalyptus woodlands and forests.

*Temporal dynamics*: Fires and post-fire succession are important in the vegetation that supports this community. Wildfires are often severe and stand-replacing (Burbidge et al. 2007). Due to the strong influence of such fires on vegetation structure and food resources, the composition of the community varies with time since the last fire (Rainsford et al. 2021). The first few years post-fire are dominated by species associated with open vegetation, whereas shrub-associated species increase in occurrence after this time, once low and dense vegetation has returned (Makdissi et al. 2024).

*Geographical variation*: There is some geographical variation for this community due to a distribution that spans western Australia, eastern Australia and Kangaroo Island (e.g. the Blue-breasted Fairy-wren is restricted to Western Australia). The Western Ground Parrot is an extremely rare and range restricted species found around Cape Arid National Park. Although this species was too rare and secretive to influence the community typology, it is worth noting that the Western Ground Parrot is probably part of this community (Burbidge et al. 2007).

***Vegetation/habitat associations***

The *Mediterranean Mallee-heath* bird community occurs primarily in mallee woodlands, and heathlands Major Vegetation Groups (MVGs). The typical species in this community prefer low dense vegetation, diverse flowering shrub layer and the presence of shrubby (mallee) eucalyptus species. The vegetation and bird bird species associated with this bird community are distinctly more mesic than similar communities in this region (e.g. Shrubby Mallee).

#### **Open Mallee**

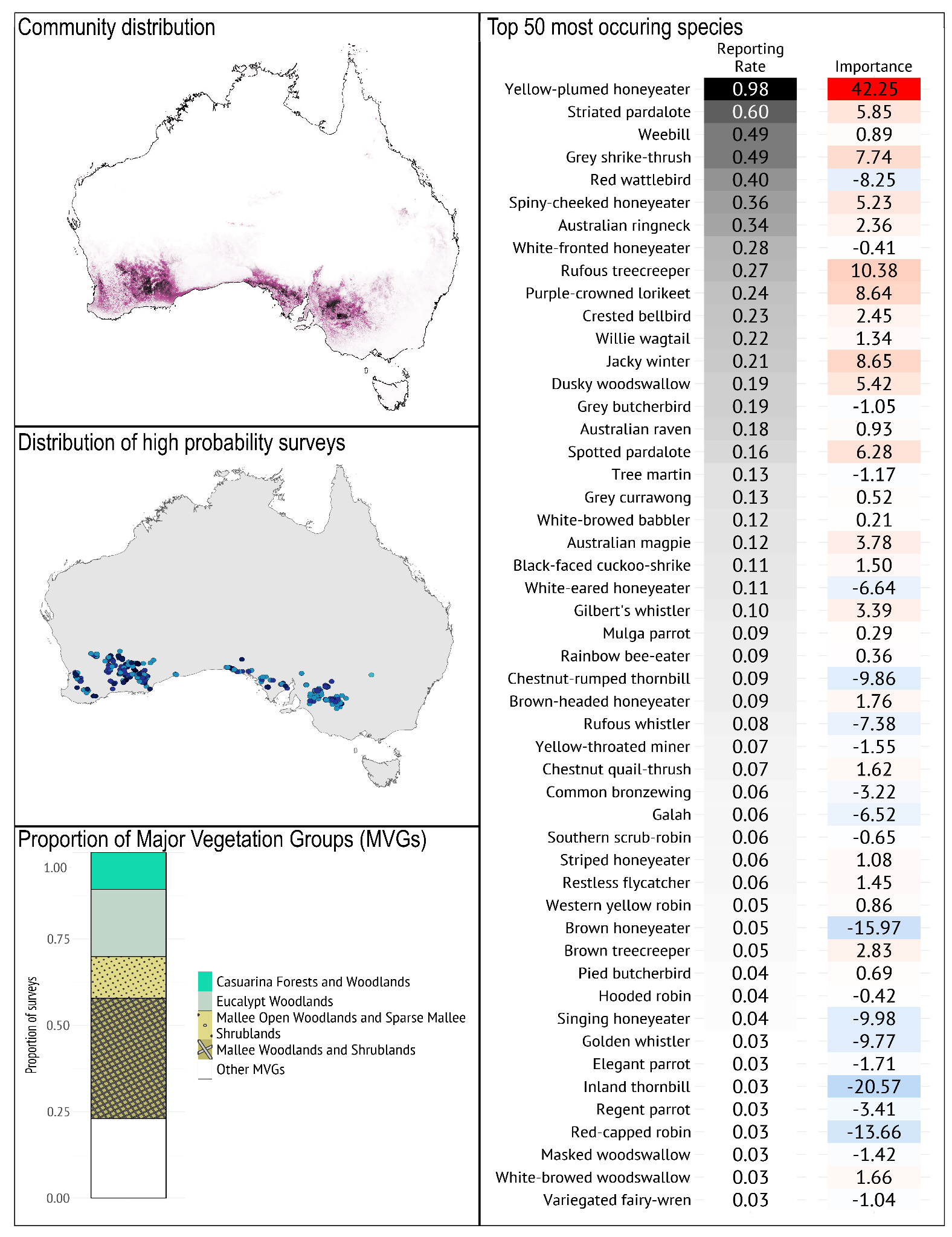

***Geographical occurrence***

The *Open Mallee* bird community (like the similar *Shrubby Mallee* bird community) stretches from south-west Western Australia, to western Victoria and south-west New South Wales, with its range split by the Nullarbor Plain. This community occurs in semi-arid regions with an average annual rainfall of 300-400mm. The areas with the highest probability of this community occurring include areas where more remnant vegetation remains such as Norseman and surrounds (WA), the Bookmark Biosphere Reserve (SA) and Murray Sunset National Park (Victoria).

***Species composition***

Typical species: When intact, this community is distinguished by the presence of the closely mallee-associated Yellow-plumed Honeyeater (by far the most important indicator for this community), Rufous Treecreeper, Jacky Winter, Purple-crowned Lorikeet and Grey Shrike-thrush.

*Guild structure*: Although *Shrubby Mallee* and *Open Mallee* bird communities had similar distributions, their guild structures differed substantially. *Open Mallee* had higher rates of occurrence for species associated with taller mallee woodlands (Yellow-plumed Honeyeater, Striated Pardalote, Grey Shrike-thrush, Jacky Winter, Spiny-cheeked Honeyeater and Rufous Treecreeper).

*Temporal dynamics*: Fires and post-fire succession are important in the vegetation that supports this community (Taylor et al. 2012; Gosper et al. 2019). Wildfires are often severe and stand-replacing (Clarke et al. 2021). Due to the strong influence of such fires on vegetation structure and food resources, the composition of the community varies with time since the last fire (Gosper et al. 2019). Regenerating mallee vegetation can form a low and dense shrubland in the first 10 years after fire, resulting in an increased prevalence in shrub-associated bird species (Rainsford et al. 2020, 2023). Whereas in mature Open Mallee, there is a mostly open understorey and taller mallee Eucalyptus canopy, with a mix of bareground and deep leaf litter around the bases of trees. As a result, litter foraging, bark foraging and canopy foraging species are more common in mature Open Mallee (Gosper et al. 2019).

*Geographical variation*: In the west, the Rufous Treecreeper is common but this is replaced by Brown Treecreeper in the east. The Brown Treecreeper remains a positive indicator of this community despite its overall low rate of occurrence. Although not shown in the typology, the Open Mallee bird community is likely to comprise multiple rare and threatened mallee specialist bird species in the eastern mallee (Boulton et al. 2021; Verdon & Clarke 2022).

***Vegetation/habitat associations***

The *Open Mallee* bird community occurs primarily in mallee woodlands, and Eucalypt woodlands Major Vegetation Groups (MVGs), where the understorey is moderately open, comprising shrubs and hummock-grass (*Triodia* spp.) and the ground layer comprises a mix of bareground and deep litter at the bases of trees. As a result, most species associated with this community forage in the canopy and bark of taller mallee trees, with occasional ground foraging species associated with leaf litter (e.g. Chestnut Quail-thrush).

#### **Shrubby Mallee**

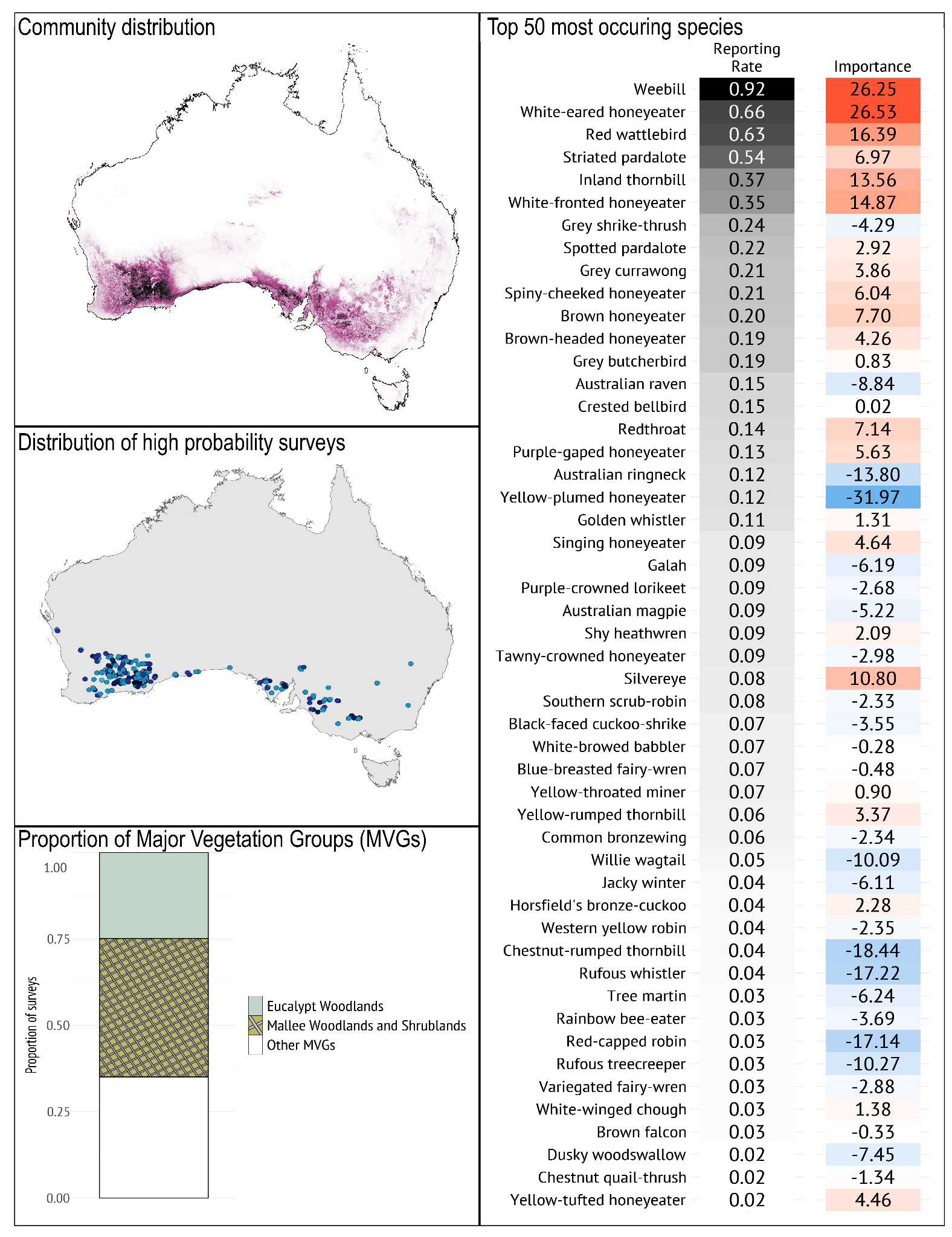

***Geographical occurrence***

The *Open Mallee* and *Shrubby Mallee* bird communities both stretch from south-west Western Australia, to western Victoria and south-west New South Wales, interrupted by the Nullarbor Plain. They occur in semi-arid regions with an annual average of 300-400mm of rain. The Shrubby Mallee community is less likely to occur in the eastern extent of its range than the Open Mallee community and has a higher occurrence in Norseman and surrounds. In its eastern extent, the Shrubby Mallee community occurs further south than the Open Mallee community.

***Species composition***

*Typical species*: The *Shrubby Mallee* bird community is typified by high prevalence of Weebill, White-eared Honeyeater, Red Wattlebird, Inland Thornbill and White-fronted Honeyeater. The relatively low rate of occurrence of Yellow-plumed Honeyeater (compared to Open Mallee) is an important distinguishing feature of the Shrubby Mallee community.

*Guild structure*: Although *Shrubby Mallee* and *Open Mallee* bird communities had similar distributions, their guild structures differed substantially. *Shrubby Mallee* had higher rates of occurrence for shrub associated species (e.g. White-eared Honeyeater, Inland Thornbill and White-fronted Honeyeater) and lower rates for many of the woodland associated species that define the Open Mallee community (e.g. Grey Shrike-thrush, Australian Ringneck, Rufous Whistler, Rufous Treecreeper). The Redthroat is another uncommon shrub-dwelling species that is more likely to occur in this community than *Open Mallee*.

*Temporal dynamics*: Fires and post-fire succession are important in the vegetation that supports this community. Wildfires are often severe and stand-replacing (Clarke et al. 2021). Due to the strong influence of such fires on vegetation structure and food resources, the composition of the community varies with time since the last fire (Makdissi et al. 2024). After a brief period that favours open habitat species such as the Yellow-rumped Thornbill, regenerating Shrubby Mallee supports a diverse and often dense shrub layer with many species providing important nectar resources. This regenerating vegetation can form a low and dense shrubland in the first 10 years after fire, resulting in an increased prevalence in shrub-associated bird species (Clarke et al. 2021). Mature Shrubby Mallee still supports a diverse bird assemblage. However, after longer fire-free periods (e.g. > 40 yrs), the vegetation can senesce, becoming open and less diverse, resulting in decreased bird abundance and richness (Verdon et al. 2024; Makdissi et al. 2024). Alternatively, the absence of fire can also allow monotypic stands of the inland pine *Callitris spp.* to develop, resulting in a bird community with different characteristics and generally lower richness (Bradstock et al. 2006).

*Geographical variation*: Although this community spans Western and Eastern Australia, most important species occur on both sides of the Nullarbor Plain, resulting in limited geographic variation in the Shrubby Mallee community. Some rare, range restricted species are associated with this community in eastern Australia. Although they are not represented in the typology due to their rarity (e.g. Mallee Emu-wren and Red-lored Whistler; Verdon et al. 2024).

***Vegetation/habitat associations***

The *Shrubby Mallee* bird community (like the *Open Mallee* bird community) occurs primarily in mallee woodlands, and Eucalypt woodlands Major Vegetation Groups (MVGs) with mallee-type trees and a well-developed shrub layer. In general the mallees form a low shrubland or open shrubland rather than a taller woodland. In addition, *Shrubby Mallee* supports a more diverse and structurally important layer of flowering shrub species, resulting in a greater prevalence of nectarivorous Honeyeaters (e.g. White-fronted Honeyeater and Purple-gaped Honeyeater).

#### **Inland Woodland with Eucalypt**

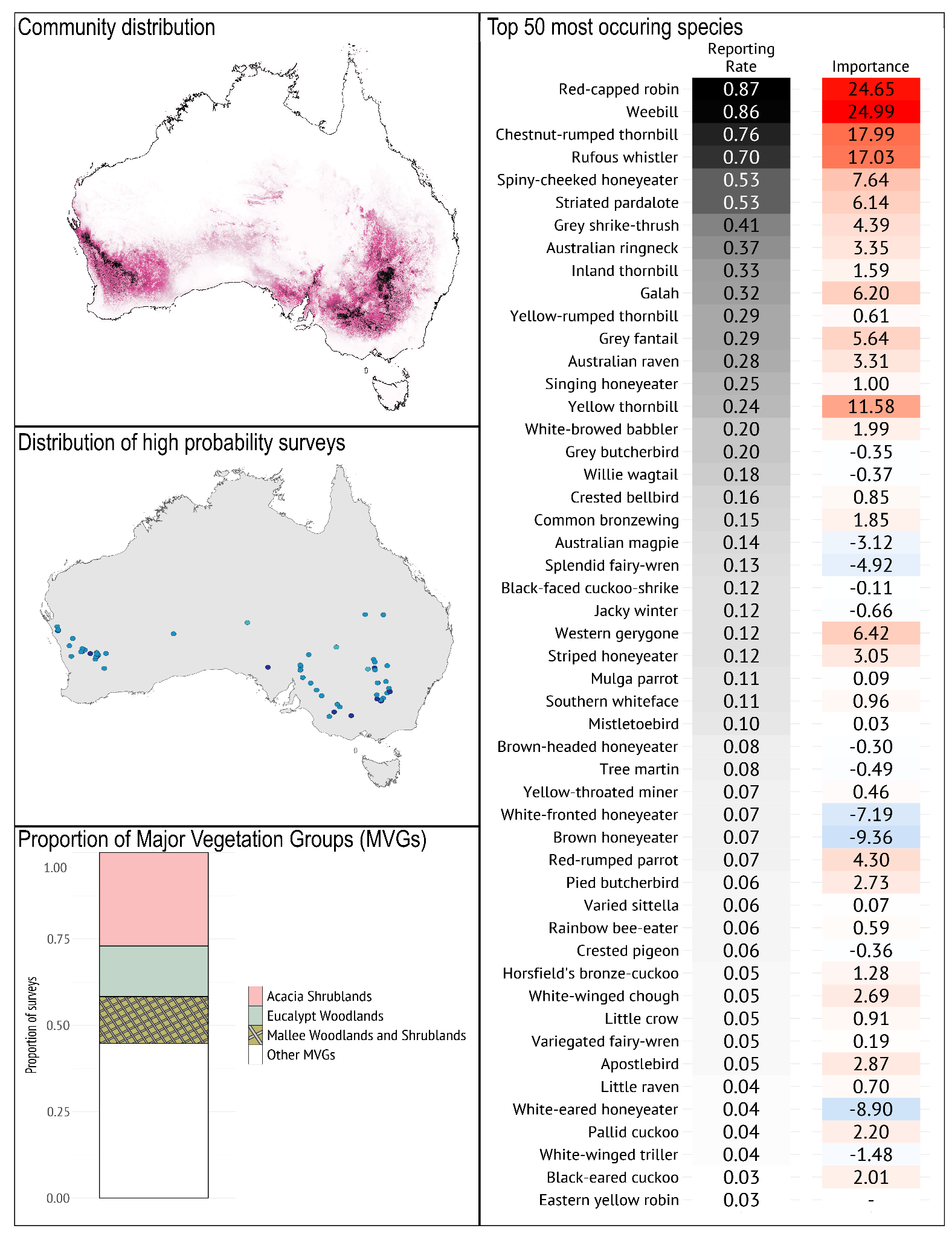

***Geographical occurrence***

The *Inland Woodland with Eucalypt* bird community has a similar pattern of occurrence to the *Mulga* bird community, spanning areas with between 300 and 600mm rainfall, primarily in winter, as high as Carnarvon (WA) and Jundah (QLD). The highest probability of occurrence for this community is around the junction of the Darling and Bogan Rivers (NSW) and the Murray-Sunset National Park (Vic), and along the series of almost-connecting lakes and waterways along the eastern fringe of the WA Wheat/Sheep belt (WA).

***Species composition***

*Typical species*: The *Inland Woodland with Eucalypt* bird community is typified by the prevalence of Red-capped Robin, Weebill, Chestnut-rumped Thornbill, Rufous Whistler and Spiny-cheeked Honeyeater. Its species composition is most similar to the *Mulga* bird community, sharing the distinctive high prevalence of Chestnut-rumped Thornbill, Red-capped Robin, and Spiny-cheeked Honeyeater. It is distinct from the *Mulga* bird community in having a higher prevalence of Weebill, Striated Pardalote, Rufous Whistler, and Grey Shrike-thrush and lower prevalence of Splendid Fairy-wren.

*Guild structure*: The *Inland Woodland with Eucalypt* is dominated by small to medium insectivores, with a relatively low diversity and prevalence of nectarivores and few granivores.

*Temporal dynamics*: In northern NSW and southern QLD, variable summer rainfall is a primary driver of sparse *Acacia* dominated Mulga vegetation community, while infrequent fire (inter-fire interval 40 – 50 years) is a driver of eucalypt-dominated systems in more Mediterranean parts of this distribution. When fire-sensitive Acacia shrublands are burned, the bird community typical of inland Eucalypt woodlands is likely to shift significantly, with a greater representation of generalist species for 10–15 years post-fire (Woinarski 1999).

*Geographical variation*: The majority of the important species in this typology span eastern and western Australia. The Yellow Thornbill and Red-rumped Parrot are representative of the eastern taxa.

***Vegetation/habitat associations***

The *Inland Woodland with Eucalypt* bird community occurs primarily in Acacia shrublands, and Eucalypt woodlands Major Vegetation Groups (MVGs). Locations situated in the drier regions of the geographical range are characterised by multi-stemmed, low *Acacia* shrubs (<2 m), while in more mesic parts of the distribution give way to single-stemmed acacia of greater height (<6 m). In eucalypt-dominated systems, trees are widely spaced with a ground layer that is dominated by tussock grasses with forbs in south-eastern Australia, while being relatively sparse in south-western Australia. *Acacia* or *Callitris* may occur in a subcanopy, providing key resources for gleaning insectivores (Pavey & Nano 2009).

#### **Mulga**

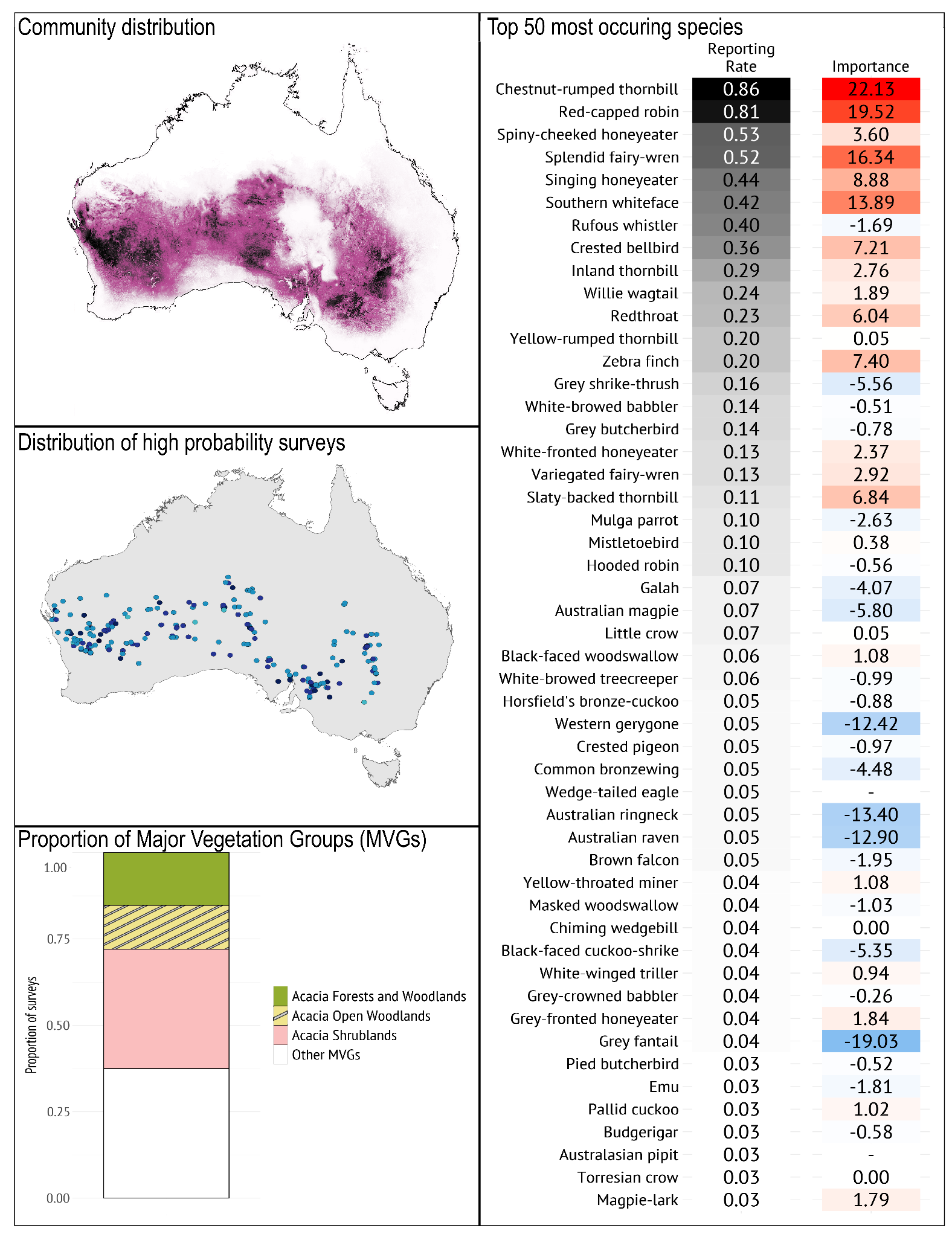

***Geographical occurrence***

The distribution of the *Mulga* bird community corresponds with the mapped distribution of mulga *Acacia aneura* shrublands (Eamus et al. 2016). The Mulga bird community has a similar pattern of occurrence to the *Inland Woodland with Eucalypt* bird community, but includes areas further north. It spans areas with between 200 and 600 mm rainfall, primarily in winter, as far north as Exmouth (WA) and McKinlay (QLD). It has the same occurrence hotspots to the *Inland Woodland with Eucalypt bird* community, but extending in a wider radius around the same points: the junction of the Darling and Bogan Rivers (NSW) and the Murray-Sunset National Park (Vic), and along the series of almost-connecting lakes and waterways along the eastern fringe of the WA Wheat/Sheep belt (WA). It also has a high occurrence in the Maralinga Tjarutja lands and surrounds (SA).

***Species composition***

*Typical species*: The *Mulga* bird community is typified by the prevalence of Chestnut-rumped Thornbill, Red-capped Robin, Spiny-cheeked Honeyeater, Splendid Fairy-wren, and Singing Honeyeater. Its species composition is most similar to the *Inland Woodland with Eucalypt* bird community, sharing distinctive high prevalence of Chestnut-rumped Thornbill, Red-capped Robin, and Spiny-cheeked Honeyeater. It is distinct from the *Inland Woodland with Eucalypt* bird community in its higher prevalence of Splendid Fairy-wren and Inland Thornbill and lower prevalence of Weebill, Striated Pardalote, Rufous Whistler, and Grey Shrike-thrush. Redthroat and Slaty-backed Thornbill are characteristic species of more intact sites.

*Guild structure*: The community is dominated by a diversity of small- to large-bodied insectivores that forage at various strata from the ground to the canopy, which is a consistent finding with avifauna studies in mulga habitats (Recher & Davis Jr 1997; Recher 2018); the common honeyeaters (Spiny-cheeked and Singing) are quite insectivorous.

*Temporal dynamics*: The *Mulga* bird community appears to exhibit relatively little seasonal variation across a year as compared with other dryland bird communities (Cody 1994; Reid et al. 2024). But is sensitive to large climatic changes, with wet conditions driving an influx of migratory and nomadic birds which are largely absent in dry conditions (Recher 2018).

*Geographical variation*: Species composition varies slightly depending on the available species pool in the area, variation in mulga stands and their position in the landscape (Burbidge et al. 2000, 2010). Slaty-backed Thornbill and White-browed Treecreeper, for example, have a narrower distribution than that of the community.

***Vegetation/habitat associations***

The *Mulga* bird community occurs primarily in Acacia shrublands, and Acacia forests and woodlands Major Vegetation Groups (MVGs) - essentially, those dominated by Mulga *A. aneura*.

**References**

Abbott I. 1972. Land bird migration across Bass Strait. Page Birds in Bass Strait. A. H. & A. W. Reed Pty Ltd.

Antos MJ, Bennett AF. 2005. How important are different types of temperate woodlands for ground-foraging birds? Wildlife Research 32:557.

Antos MJ, Bennett AF. 2006. Foraging ecology of ground-feeding woodland birds in temperate woodlands of southern Australia. Emu - Austral Ornithology 106:29–40.

Armstrong JA. 1979. Biotic pollination mechanisms in the Australian flora — a review. New Zealand Journal of Botany 17:467–508.

Arnold G. 1988. The Effects of Habitat Structure and Floristics on the Densities of Bird Species in Wandoo Woodland. Wildlife Research 15:499.

Atlas Of Living Australia. 2024. Occurrence download records-2024-12-19DOI: 10.26197/ALA.CEA103AD-A447-4EA3-93BD-FCD6589D5BCE. Atlas Of Living Australia. Available from https://doi.ala.org.au/doi/cea103ad-a447-4ea3-93bd-fcd6589d5bce (accessed December 19, 2024).

Bennett AF, Ford LA. 1997. Land use, habitat change and the conservation of birds in fragmented rural environments: a landscape perspective from the Northern Plains, Victoria, Australia. Pacific Conservation Biology 3:244.

Bennett AF, Haslem A, Garnett ST, Loyn RH, Woinarski JCZ, Ehmke G. 2024. Declining but not (yet) threatened: a challenge for avian conservation in Australia. Emu - Austral Ornithology 124:123–145.

Bennett JM, Nimmo DG, Clarke RH, Thomson JR, Cheers G, Horrocks GFB, Hall M, Radford JQ, Bennett AF, Mac Nally R. 2014. Resistance and resilience: can the abrupt end of extreme drought reverse avifaunal collapse? Diversity and Distributions 20:1321–1332.

Boulton RL, Clarke R, Verdon SJ, Hedger C, Ireland L, Garnett ST. 2021. Black-eared Miner: Manorina melanotis Wilson, 1911: Meliphagidae. Page in Garnett S, Baker GB, editors. The Action Plan for Australian Birds 2020. CSIRO Publishing, Clayton South, VIC.

Bradstock RA, Bedward M, Cohn JS. 2006. The modelled effects of differing fire management strategies on the conifer Callitris verrucosa within semi‐arid mallee vegetation in Australia. Journal of Applied Ecology 43:281–292.

Brereton R, Taylor R. 2000. Composition, seasonal occurrences and habitat use of bird assemblages in wet forests on the Central Plateau of Tasmania. Papers and Proceedings of the Royal Society of Tasmania 134:35–43.

Brooker MG, Ridpath MG, Estbergs AJ, Bywater J, Hart DS, Jones MS. 1979. Bird Observations on the North-Western Nullarbor Plain and Neighbouring Regions, 1967–1978. Emu - Austral Ornithology 79:176–190.

Burbidge AA, Fuller PJ. 2007. Gibson Desert birds: responses to drought and plenty. Emu - Austral Ornithology 107:126–134.

Burbidge AH, Johnstone RE, Fuller PJ, Stone P. 2000. Terrestrial birds of the southern Camarvon Basin, Western Australia: contemporary patterns of occurrence. Records of the Western Australian Museum, Supplement 60:449.

Burbidge AH, Johnstone RE, Pearson DJ. 2010. Birds in a vast arid upland: avian biogeographical patterns in the Pilbara region of Western Australia. Records of the Western Australian Museum, Supplement 78:247.

Burbidge AH, Rolfe J, McNee S, Newbey B, Williams M. 2007. Monitoring population change in the cryptic and threatened Western Ground Parrot in relation to fire. Emu - Austral Ornithology 107:79–88.

Cardillo M, Pratt R. 2013. Evolution of a hotspot genus: geographic variation in speciation and extinction rates in Banksia (Proteaceae). BMC Evolutionary Biology 13:155.

Chapman A, Kofron CP. 2010. Tropical Wet Sclerophyll Forest and Bird Diversity in North-east Queensland, Australia. Pacific Conservation Biology 16:20.

Clarke MF et al. 2021. Fire and Its Interactions With Other Drivers Shape a Distinctive, Semi-Arid ‘Mallee’ Ecosystem. Frontiers in Ecology and Evolution 9:647557.

Cody ML. 1994. Mulga bird communities. I. Species composition and predictability across Australia. Australian Journal of Ecology 19:206–219.

Crates R, Terauds A, Rayner L, Stojanovic D, Heinsohn R, Ingwersen D, Webb M. 2017. An occupancy approach to monitoring regent honeyeaters. The Journal of Wildlife Management 81:669–677.

Cunningham RB, Lindenmayer DB, Crane M, Michael DR, Barton PS, Gibbons P, Okada S, Ikin K, Stein JAR. 2014. The law of diminishing returns: woodland birds respond to native vegetation cover at multiple spatial scales and over time. Diversity and Distributions 20:59–71.

Davis RA, Valentine LE, Craig MD. 2022. Do bird communities differ with post-fire age in Banksia woodlands of south-western Australia. International Journal of Wildland Fire 31:621–633.

Davis RA, Valentine LE, Craig MD, Wilson B, Bancroft WJ, Mallie M. 2014. Impact of Phytophthora-dieback on birds in Banksia woodlands in south west Western Australia. Biological Conservation 171:136–144.

De La Fuente A, Navarro A, Williams SE. 2023. The climatic drivers of long‐term population changes in rainforest montane birds. Global Change Biology 29:2132–2140.

Eamus D, Huete A, Cleverly J, Nolan RH, Ma X, Tarin T, Santini NS. 2016. Mulga, a major tropical dry open forest of Australia: recent insights to carbon and water fluxes. Environmental Research Letters 11:125011.

Eyre TJ, Maron M, Mathieson MT, Haseler M. 2009. Impacts of grazing, selective logging and hyper‐aggressors on diurnal bird fauna in intact forest landscapes of the Brigalow Belt, Queensland. Austral Ecology 34:705–716.

Ford HA, Barrett GW, Saunders DA, Recher HF. 2001. Why have birds in the woodlands of Southern Australia declined? Biological Conservation 97:71–88.

Franklin DC, Menkhorst PW, Robinson JL. 1989. Ecology of the Regent Honeyeater Xanthomyza phrygia. Emu - Austral Ornithology 89:140–154.

Gosper CR et al. 2019. Fire‐mediated habitat change regulates woodland bird species and functional group occurrence. Ecological Applications 29:e01997.

Grey MJ, Clarke MF, Loyn RH. 1998. Influence of the Noisy Miner Manorina melanocephala on avian diversity and abundance in remnant Grey Box woodland. Pacific Conservation Biology 4:55.

Griffioen PA, Clarke MF. 2002. Large-scale bird-movement patterns evident in eastern Australian atlas data. Emu - Austral Ornithology 102:99–125.

Haslem A, Nimmo DG, Radford JQ, Bennett AF. 2015. Landscape properties mediate the homogenization of bird assemblages during climatic extremes. Ecology 96:3165–3174.

Hernandez S, Adams VM, Duce S. 2024. The hidden impact of policy changes on remnant vegetation in Queensland, Australia. Land Use Policy 139:107064.

Hutley LB, Beringer J, Isaac PR, Hacker JM, Cernusak LA. 2011. A sub-continental scale living laboratory: Spatial patterns of savanna vegetation over a rainfall gradient in northern Australia. Agricultural and Forest Meteorology 151:1417–1428.

Jordan R, James AI, Moore D, Franklin DC. 2017. Boom and bust (or not?) among birds in an Australian semi-desert. Journal of Arid Environments 139:58–66.

Kikkawa J. 1968. Ecological association of bird species and habitats in eastern Australia; similarity analysis. The Journal of Animal Ecology:143–165.

Kutt AS, Vanderduys EP. 2017. Bird assemblage changes along a savannarainforest gradient in north-eastern Australia. Australian Zoologist 38:552–561.

Kutt AS, Vanderduys EP, Perry JJ, Mathieson MT, Eyre TJ. 2016. Yellow-throated miners M anorina flavigula homogenize bird communities across intact and fragmented landscapes: Homogenization of Bird Communities. Austral Ecology 41:316–327.

Leavesley AJ, Cary GJ, Edwards GP, Gill AM. 2010. The effect of fire on birds of mulga woodland in arid central Australia. International Journal of Wildland Fire 19:949.

Loyn RH. 1985. Ecology, distribution and density of birds in Victorian forests. Page Birds of Eucalypt Forests and Woodlands: Ecology, Conservation, Management. RAOU and Surrey Beatty & Sons, Sydney, NSW.

Loyn RH. 1987. Effects of patch area and habitat on bird abundances, species numbers and tree health in fragmented Victorian forests. Page in Saunders DA, Arnold GW, Burbidge AA, Hopkins AJM, editors. Nature Conservation: the Role of Remnants of Native Vegetation. Surrey Beatty & Sons, Chipping Norton, NSW.

Lunn TJ, Gerwin M, Buettel JC, Brook BW. 2018. Impact of intense disturbance on the structure and composition of wet-eucalypt forests: A case study from the Tasmanian 2016 wildfires. PLOS ONE 13:e0200905.

Lunt ID, Bennett AF. 2000. Temperate woodlands in Victoria: distribution, composition and conservation. Pages 17–31 Temperate Eucalypt Woodlands in Australia: Biology, Conservation, Management and Restoration. Surrey Beatty & Sons, Chipping Norton, NSW.

Mac Nally R. 1995. On Large-Scale Dynamics and Community Structure in Forest Birds: Lessons from Some Eucalypt Forests of Southeastern Australia. Philosophical Transactions: Biological Sciences 350:369–379. Royal Society.

Mac Nally R, Bennett AF, Thomson JR, Radford JQ, Unmack G, Horrocks G, Vesk PA. 2009. Collapse of an avifauna: climate change appears to exacerbate habitat loss and degradation. Diversity and Distributions 15:720–730.

Mac Nally R, Bowen M, Howes A, McAlpine CA, Maron M. 2012. Despotic, high‐impact species and the subcontinental scale control of avian assemblage structure. Ecology 93:668–678.

Mac Nally R, Horrocks G. 2000. Landscape-scale conservation of an endangered migrant:the Swift Parrot (Lathamus discolor) in its winter range. Biological Conservation 92:335–343.

Mac Nally R, Horrocks GFB, Bennett JM, Yen JDL, Selwood KE, Thomson JR, Lada H. 2020. Ecological and life‐history traits may say little about birds’ vulnerability to high‐amplitude climatic fluctuations. Austral Ecology 45:880–895.

Mac Nally R, Kutt AS, Eyre TJ, Perry JJ, Vanderduys EP, Mathieson M, Ferguson DJ, Thomson JR. 2014. The hegemony of the ‘despots’: the control of avifaunas over vast continental areas. Diversity and Distributions 20:1071–1083.

Mac Nally R, McGoldrick JM. 1997. Landscape Dynamics of Bird Communities in Relation to Mass Flowering in Some Eucalypt Forests of Central Victoria, Australia. Journal of Avian Biology 28:171.

Mac Nally R, Timewell CAR. 2005. Resource Availability Controls Bird-Assemblage Composition Through Interspecific Aggression. The Auk 122:1097–1111.

Makdissi R, Verdon SJ, Radford JQ, Bennett AF, Clarke MF. 2024. The impact of plant‐derived fire management prescriptions on fire‐responsive bird species. Ecological Applications:e3036.

Maron M, Grey MJ, Catterall CP, Major RE, Oliver DL, Clarke MF, Loyn RH, Mac Nally R, Davidson I, Thomson JR. 2013. Avifaunal disarray due to a single despotic species. Diversity and Distributions 19:1468–1479.

McGoldrick JM, Mac Nally R. 1998. Impact of flowering on bird community dynamics in some central Victorian eucalypt forests. Ecological Research 13:125–139.

Milledge DR, Recher HF. 1985. A comparison of forest bird communities on the New South Wales south and mind-north coasts. Pages 47–52 in Keast A, Recher HF, Saunders D, editors. Birds of Eucalypt Forests and Woodlands: Ecology, Conservation, Management,. Surrey Beatty & Sons, Sydney, NSW.

Mollenmans FH, Reid JRW, Thompson MB, Alexander L, Pedler LP. 1984. Biological survey of the Cooper Creek Environmental Association (8.4.4), North Eastern South Australia. Consultancy Report. NPWS Department of Environment and Planning, Adelaide.

Paton DC, Prescott AM, Davies RJ-P, Heard LM. 1999. The distribution, status and threats to temperate woodlands in South Australia. Page Temperate Eucalypt Woodlands in Australia: Biology, Conservation, Management and Restoration. Surrey Beatty & Sons, Chipping Norton, NSW.

Pavey CR, Nano CEM. 2009. Bird assemblages of arid Australia: Vegetation patterns have a greater effect than disturbance and resource pulses. Journal of Arid Environments 73:634–642.

Radford JQ, Bennett AF. 2005. Terrestrial avifauna of the Gippsland Plain and Strzelecki Ranges, Victoria, Australia: insights from Atlas data. Wildlife Research 32:531.

Radford JQ, Bennett AF. 2007. The relative importance of landscape properties for woodland birds in agricultural environments. Journal of Applied Ecology 44:737–747.

Radford JQ, Bennett AF, Cheers GJ. 2005. Landscape-level thresholds of habitat cover for woodland-dependent birds. Biological Conservation 124:317–337.

Rainsford FW, Giljohann KM, Bennett AF, Clarke MF, MacHunter J, Senior K, Sitters H, Watson S, Kelly LT. 2023. Ecosystem type and species’ traits help explain bird responses to spatial patterns of fire. Fire Ecology 19:59.

Rainsford FW, Kelly LT, Leonard SWJ, Bennett AF. 2020. Post‐fire development of faunal habitat depends on plant regeneration traits. Austral Ecology 45:800–812.

Rainsford FW, Kelly LT, Leonard SWJ, Bennett AF. 2021. Post-fire habitat relationships for birds differ among ecosystems. Biological Conservation 260:109218.

Rainsford FW, Kelly LT, Leonard SWJ, Bennett AF. 2022. Fire and functional traits: Using functional groups of birds and plants to guide management in a fire‐prone, heathy woodland ecosystem. Diversity and Distributions 28:372–385.

Recher HF. 2018. Foraging behaviour of mulga birds in Western Australia. II. Community structure and conservation. Pacific Conservation Biology 24:87.

Recher HF, Davis Jr WE. 1997. Foraging Ecology of a Mulga Bird Community. Wildlife Research 24:27.

Recher HF, Gowing G, Kavanagh R, Shields J, Rohan-Jones WD. 1983. Birds, resources and time in a tablelands forest. Pages 101–123 Mountain ecology in the Australian region: proceedings of a symposium held at Canberra, 8-9 May, 1982. Ecological Society of Australia.

Recher HF, Holmes RT, Schulz M, Shields J, Kavanagh R. 1985. Foraging patterns of breeding birds in eucalypt forest and woodland of southeastern Australia. Australian Journal of Ecology 10:399–419.

Recher HF, Kavanagh RP, Shields JM, Lind P. 1991. Ecological association of habitats and bird species during the breeding season in southeastern New South Wales. Australian Journal of Ecology 16:337–352.

Reid J. 1990. 14: Birds. Page Natural History of the North East Deserts. Royal Society of South Australia.

Reid J, Smith R, Scott L, Reid N. 2024. The stability of bird assemblages across time and the reliability of snapshot surveys. Austral Ecology 49:e13516.

Reid JRW. 1999. Threatened and declining birds in the New South Wales sheep-wheatbelt: 1. Diagnosis, characteristics and management. Page Report to NSW National Parks and Wildlife Service. CSIRO Sustainable Ecosystems, Canberra, ACT.

Rldpath MG, Moreau RE. 1966. The birds of Tasmania: ecology and evolution. Ibis 108:348–393.

Robinson D, Trail B. 1996. Conserving woodland birds in the wheat and sheep belts of southern Australia. RAOU Conservation Statement No. 10. RAOU (Birds Australia), Melbourne, VIC.

Saunders DA. 1990. Problems of survival in an extensively cultivated landscape: the case of Carnaby’s cockatoo Calyptorhynchus funereus latirostris. Biological Conservation 54:277–290.

Selwood KE, Clarke RH, Cunningham SC, Lada H, McGeoch MA, Mac Nally R. 2015a. A bust but no boom: responses of floodplain bird assemblages during and after prolonged drought. Journal of Animal Ecology 84:1700–1710.

Selwood KE, Thomson JR, Clarke RH, McGeoch MA, Mac Nally R. 2015b. Resistance and resilience of terrestrial birds in drying climates: do floodplains provide drought refugia? Global Ecology and Biogeography 24:838–848.

Serong M, Lill A. 2012. Changes in bird assemblages during succession following disturbance in secondary wet forests in south-eastern Australia. Emu - Austral Ornithology 112:117–128.

Specht RL. 1970. Vegetation. Pages 44–67 in Leeper GW, editor. Australian Environment, 4th edition. Melbourne University Press, Melbourne.

Spessa A, McBeth B, Prentice C. 2005. Relationships among fire frequency, rainfall and vegetation patterns in the wet–dry tropics of northern Australia: an analysis based on NOAA‐AVHRR data. Global Ecology and Biogeography 14:439–454.

Szabo JK, Vesk PA, Baxter PWJ, Possingham HP. 2011. Paying the extinction debt: woodland birds in the Mount Lofty Ranges, South Australia. Emu - Austral Ornithology 111:59–70.

Taylor RS, Watson SJ, Nimmo DG, Kelly LT, Bennett AF, Clarke MF. 2012. Landscape‐scale effects of fire on bird assemblages: does pyrodiversity beget biodiversity? Diversity and Distributions 18:519–529.

Tischler M, Dickman CR, Wardle GM. 2013. Avian functional group responses to rainfall across four vegetation types in the S impson D esert, central A ustralia. Austral Ecology 38:809–819.

Tulloch AIT, Healy A, Silcock J, Wardle GM, Dickman CR, Frank ASK, Aubault H, Barton K, Greenville AC. 2023. Long‐term livestock exclusion increases plant richness and reproductive capacity in arid woodlands. Ecological Applications 33:e2909.

Vasuki Y, Yu L, Holden E-J, Kovesi P, Wedge D, Grigg AH. 2019. The spatial-temporal patterns of land cover changes due to mining activities in the Darling Range, Western Australia: A Visual Analytics Approach. Ore Geology Reviews 108:23–32.

Verdon SJ, Clarke MF. 2022. Can fire‐age mosaics really deal with conflicting needs of species? A study using population hotspots of multiple threatened birds. Journal of Applied Ecology 59:2128–2141.

Verdon SJ, Makdissi R, Mitchell WF, Boulton RL, Radford JQ. 2024. Benefits of modelling abundance for rare species conservation: a case study with multiple birds across one million hectares. Diversity and Distributions.

Watson DM. 2011. A productivity-based explanation for woodland bird declines: poorer soils yield less food. Emu - Austral Ornithology 111:10–18.

Williams SE, De La Fuente A. 2021. Long-term changes in populations of rainforest birds in the Australia Wet Tropics bioregion: A climate-driven biodiversity emergency. PLOS ONE 16:e0254307.

Williams SE, Shoo LP, Henriod R, Pearson RG. 2010. Elevational gradients in species abundance, assemblage structure and energy use of rainforest birds in the Australian Wet Tropics bioregion. Austral Ecology 35:650–664.

Woinarski J. 1985. Foliage-gleaners of the treetops, the pardalotes. Page in Keast A, Recher HF, Ford HA, Saunders D, editors. Birds of Eucalypt Forests and Woodlands: Ecology, Conservation, Management. Surrey Beatty & Sons, Chipping Norton, NSW.

Woinarski JCZ. 1999. Fire and Australian birds: a review. Page in Gill AM, Woinarski JCZ, York A, editors. Australia’s Biodiversity: Responses to Fire: Plants, Birds and Invertebrates. Environment Australia, Canberra, ACT.

Wood SW, Murphy BP, Bowman DMJS. 2011. Firescape ecology: how topography determines the contrasting distribution of fire and rain forest in the south-west of the Tasmanian Wilderness World Heritage Area: Topography, fire and rain forest in south-west Tasmania. Journal of Biogeography 38:1807–1820.

Yen JDL, Thomson JR, Vesk PA, Mac Nally R. 2011. To what are woodland birds responding? Inference on relative importance of in‐site habitat variables using several ensemble habitat modelling techniques. Ecography 34:946–954.
