## Supplementary material for "A typology of Australian terrestrial bird communities": Supplementary Material S3 MaxentVariableTable.docx

**Supplementary Material S3: Community descriptions**

Table S3. The 11 environmental and spatial variables used in the final Maxent models to predict community distributions, their units, and any adjustments made to them in ArcMap 10.8.2 for model processing. All units are continuous.

| **Source** | **Variable** | **Units** | **Adjustments** |
| --- | --- | --- | --- |
| WorldClim | Precipitation in the driest quarter | Millimetres (mm) | Clipped to Australian border shapefile. |
|  | Annual precipitation | Millimetres (mm) |  |
|  | Precipitation in the wettest quarter | Millimetres (mm) |  |
|  | Precipitation seasonality | Coefficient of variation of monthly total precipitation |  |
|  | Minimum temperature of the coldest month | Degrees Celsius (°C) |  |
|  | Mean annual temperature | Degrees Celsius (°C) |  |
|  | Maximum temperature of the warmest month | Degrees Celsius (°C) |  |
|  | Temperature seasonality | Standard deviation of mean monthly temperature x100 |  |
| TERN | Total vegetation cover | (Percentage x100)+1 | Resampled to same resolution (30 seconds, ~1km) and extent as WorldClim variables. |
| Geoscience Australia | Elevation | Meters (m) | Resampled to same resolution (30 seconds, ~1km) and extent as WorldClim variables. |
|  | Distance from Water | Decimal degrees | Created using Euclidean distance from the Digital Earth Australia Waterbodies v2 layer; resolution and extent matched to WorldClim variables. |
